# Reductive death is averted by an ancient metabolic switch

**DOI:** 10.1101/2025.11.24.690166

**Authors:** Fasih M. Ahsan, Jen F. Rotti, Armen I. Yerevanian, Sinclair W. Emans, Nicole L. Stuhr, Jose A. Aceves-Salvador, Wei Wang, Daniel J. Baker, Dimitra Pouli, Owen S. Skinner, Michael D. Blower, Alexander A. Soukas

## Abstract

Biguanides, including metformin, the world’s most prescribed oral hypoglycemic, extend health-span and lifespan in vertebrates and invertebrates. Given the widespread use and apparent safety of metformin, it is assumed that its effects are not associated with toxicity, except when in marked excess. Here we determine that accumulation of damaging reducing equivalents is an unanticipated toxicity associated with biguanides, the defense against which requires post-transcriptional protection of *de novo* fatty acid biosynthesis. We demonstrate that biguanide treatment during impaired fatty acid biosynthesis drives NADPH toxicity, leading to catastrophic elevation of NADH/GSH reducing equivalents and accelerated death across metazoans. Multiple NADPH-generating interventions require fatty acid biosynthesis to prevent markedly shortened survival, indicating that this defense mechanism is broadly leveraged. We propose that fatty acid biosynthesis is a tunable rheostat which can minimize biguanide-induced reductive stress whilst maximizing its pro-longevity outcomes and serve as an exploitable vulnerability in reductive stress sensitive cancers.

**HIGHLIGHTS:**

- Biguanides inhibit cytosolic mRNA translation to extend lifespan in *C. elegans*.
- Fatty acid synthesis is translationally protected by eIF3 complex subunits.
- *pod-2*/*fasn-1* inactivation amplifies biguanide-induced reductive stress and death.
- NADPH-generating insults require fatty acid synthesis to buffer reductive stress.

## INTRODUCTION

Repurposing of biguanides, including metformin, the most widely prescribed drug for the management of type 2 diabetes mellitus, has been suggested for the treatment of various cancers, cardiovascular disease, inflammation, neurodegenerative illnesses, and other age-related disorders^1–6^. Biguanides have been shown to extend the lifespan and various health-span metrics of metazoan models, including *Caenorhabditis elegans*, *Drosophila melanogaster*, *Mus musculus*, and non-human primates^4,7–16^. Strikingly, human epidemiological studies have suggested that metformin use correlates with increased maximal survival and lowering of all-cause mortality when controlling for diabetes incidence, though these findings have more recently been called into question^4,6,7,17^. Nonetheless, these prior studies have generated interest in evaluating metformin as a geroprotective therapeutic through the Metformin in Longevity Study (MILES) and Targeting Aging with Metformin (TAME) clinical trials^4,6,7^. Thus, identification of the genetic factors that underlie the geroprotective mechanisms of biguanides are required to maximize the drugs’ therapeutic potential.

The molecular targets of biguanides in aging have not been conclusively assigned. While several studies have implicated mitochondrial electron transport chain (ETC) complex I as the major molecular target of biguanides, other studies have suggested alternative targets, including the gut microbiome, mitochondrial glycerol-3-phosphate dehydrogenase (mGPDH), ETC complex IV, and lysosomal membrane protein presenilin enhancer 2 (PEN2)^1,6,8,12,14,18–26^. Additionally, the effects of biguanides on downstream metabolic pathways depend on the circulating dose, severity of accumulation, and the tissue of exposure, among others^9,12,24,26,27^. Of note, the effects of biguanides on cellular redox state depend on their proposed biological target(s). Targeting mitochondrial electron transport chain (ETC) complex I inhibits NADH oxidation and ubiquinone reduction, raising the ratio of NADH/NAD+ and lowering the ratio of ATP/AMP^6,28^. Similarly, targeting mGPDH or ETC complex IV enhances cytosolic NADH/NAD+ and AMP/ATP ratios, and other studies suggest biguanides directly elevate hepatic reduced glutathione (GSH) pools to suppress gluconeogenesis^18,20–22,26^. Thus, major proposed mechanisms for biguanide action converge on elevation of distinct reducing equivalent pools. While biguanide-mediated elevation of reducing equivalents may correlate with beneficial health outcomes, elevated electron pressure is known to induce maladaptive reductive stress, leading to mitochondrial dysfunction, enhanced ATP exhaustion, tissue-wide inflammation, and the onset of cardiomyopathies and cancers^29–31^. It remains unknown whether specific compensatory mechanisms are activated to prevent unchecked reductive stress and ATP exhaustion during biguanide action, and whether such a mechanism is required to enable its pro-longevity outcomes.

Biguanides and mRNA translation inhibition are geroprotective therapeutic strategies converging upon overlapping cellular hallmarks of longevity^32–34^. In *C. elegans* and *Saccharomyces cerevisiae*, translation inhibition has been shown to extend lifespan and health-span outcomes^32–34^. Additionally, multiple pro-longevity paradigms, including reduced insulin/IGF-1 signaling, mitochondrial ETC disruption, and dietary restriction mimicry all result in dampened mRNA translation and reduced protein synthesis, suggesting that altered mRNA translation may function as a downstream effector of multiple pro-longevity outcomes^35–42^. High-dose metformin is shown to decrease protein synthesis and mRNA translation rates in cancer cell models downstream of inhibition of the mammalian target of rapamycin complex I (mTORC1)^43,44^. However, comparison of differentially translated mRNAs in metformin-versus mTOR inhibitor-treated cells reveals disparate effects on the translatome, indicating that metformin may also shape translation through mTOR-independent mechanisms^43^. Moreover, it is unclear if the pro-longevity mechanisms of biguanides mirror the loss of protein synthesis seen in cancer models, as biguanide responsive *C. elegans* lack known orthologs of translation-driving mTORC1 effector 4E-BP1 and regulator TSC1/TSC2^45^. Given the differential protein synthesis demands across lifespan, there is a need to identify the mechanisms by which translation inhibition promotes healthy aging, and the intersection of longevity attributed to translation inhibition and biguanide action.

Here, we leveraged functional genetic, translatomic, and metabolomic strategies in *C. elegans* to identify that *de novo* fatty acid biosynthesis critically mitigates reductive stress, an unanticipated biguanide-induced toxicity. Lifespan extending doses of the biguanide phenformin in nematodes inhibit cytosolic translation initiation. Despite this global dampening of cytosolic mRNA translation, biguanide treatment translationally protects fatty acid synthase (*fasn-1/CeFASN*) and acetyl-CoA carboxylase (*pod-2/CeACC*) transcripts to prevent a catastrophic synthetic lethal interaction. We find that EGL-45/CeEIF3A and EIF-3.B/CeEIF3B, members of the eIF3 translation initiation complex, protect biguanide-induced *pod-2* mRNA translation and pro-longevity outcomes. We further demonstrate that synthetic lethality resulting from combined biguanide treatment and loss of fatty acid synthetic capacity is not due to a loss of palmitate production, but rather due to a loss of NADPH consumption, resulting in an elevated NADPH, NADH and thiol reductive stress state coupled to ATP exhaustion. Indicating broad mechanistic conservation, NADPH-generating metabolic interventions require *de novo* fatty acid biosynthesis to buffer reductive stress, acting as a major sink to mitigate NADPH toxicity. We demonstrate that this synthetic lethality is mirrored across multiple metazoans and is conserved all the way to humans, suggesting an ancient, conserved role for fatty acid biosynthesis in mitigating reductive stress toxicity, while reciprocally enabling beneficial health and pro-longevity outcomes with biguanide administration. In sum, we find that modulating *de novo* fatty acid biosynthesis may act as a tunable metabolic vulnerability in reductive stress sensitive disorders, including various cancers.

## RESULTS

### Genome-wide functional analysis implicates attenuation of mRNA translation in biguanide-prompted longevity

To identify additional effectors of biguanide-mediated pro-longevity outcomes, we expanded our previous RNA interference (RNAi) screens by targeting 86% of all *C. elegans* coding sequences (CDS). Based on our prior observations that genes involved in biguanide-mediated growth inhibition also participate in its pro-longevity responses, we selected knockdowns that mediate the growth inhibitory properties of 150 mM metformin in *C. elegans* (Figure 1A, Table S1)^14,15^. This analysis identified genes that phenocopied metformin’s growth inhibitory effects. GO analysis of these genes revealed enrichment in cytosolic ribosomal subunits and mRNA translational machinery, suggesting that metformin may inhibit growth by targeting mRNA translation (Figure 1B).

**Figure 1.**
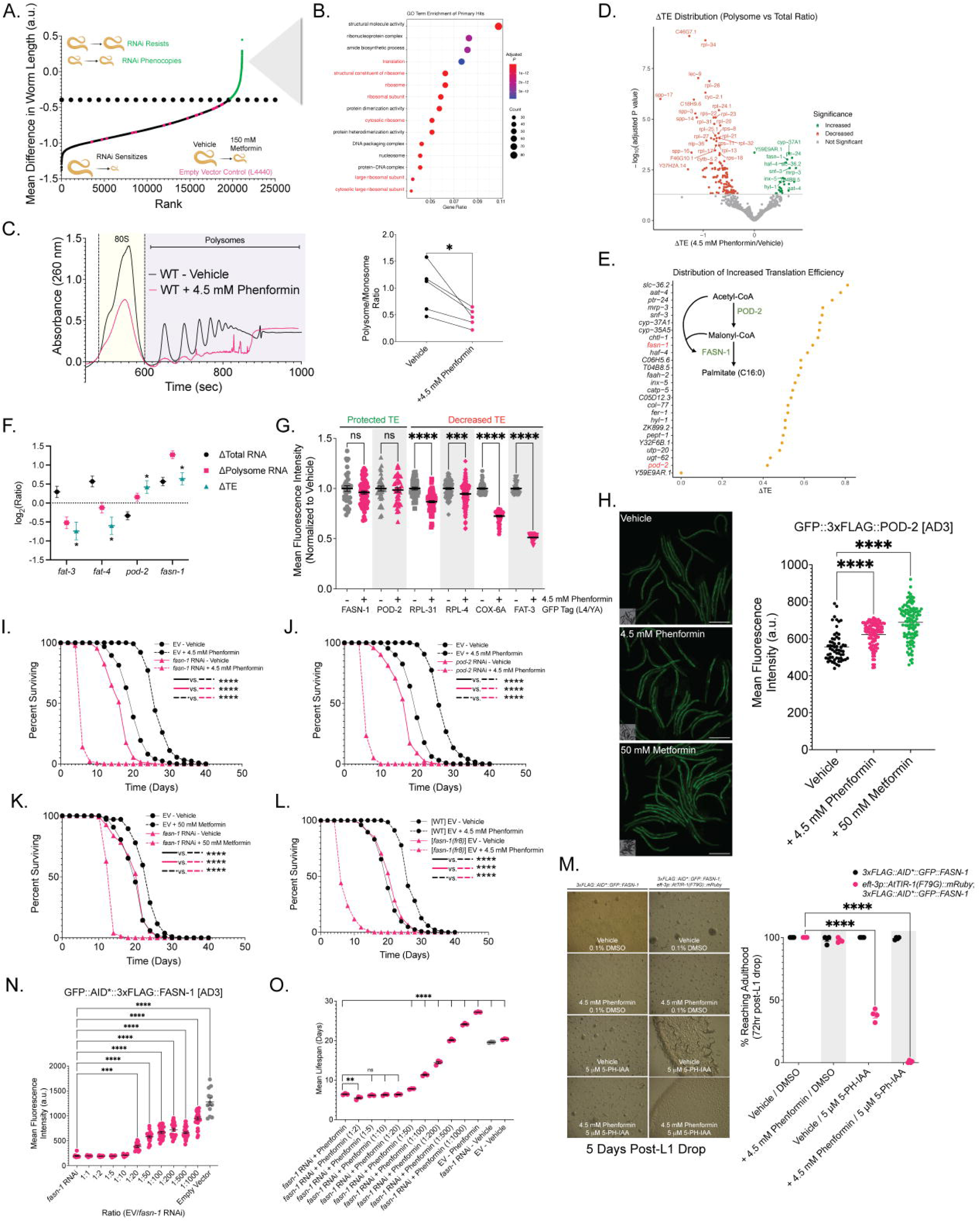
Biguanides protect translation efficiency of *de novo* fatty acid biosynthesis genes *fasn-1* and *pod-2* to prevent synthetic lethality. (A) Genome-wide RNAi screen for metformin-mediated growth inhibition. Y-axis shows the mean difference in worm length between 150 mM metformin and vehicle (n = 2 biological replicates). (B) GO term enrichment of genes whose RNAi phenocopied metformin-mediated growth inhibition (BH adjusted *p* < 0.05). (C) Representative polysome traces of wild-type *C. elegans* (N2 - WT) treated ± 4.5 mM phenformin from L1 to L4/young adulthood (YA). Inset shows polysome/monosome AUC with phenformin (n = 5 biological replicates). (D) Translation efficiency shifts with 4.5 mM phenformin, quantified as polysome occupancy versus total RNA expression (DESeq2 adjusted *p* value < 0.05) (n = 3 biological replicates). (E) Phenformin-induced translation efficiency of 26 genes. Inset: *de novo* fatty acid biosynthesis pathway with *pod-2* and *fasn-1* in green. (F) Expression changes for indicated genes in total RNA, polysomal RNA, or ΔTE scores. (G) GFP quantification of endogenously GFP-tagged *C. elegans* strains treated with 4.5 mM phenformin from L1 to L4/YA. (H) Representative micrographs and quantification of GFP::3xFLAG:POD-2 animals treated with 4.5 mM phenformin or 50 mM metformin from L4 to AD3. Note that the vehicle measurements in these graphs are identical to those shown in 5K and 6F, as these quantifications were performed in parallel. Scale bar: 500 µM. (I-K) Lifespan analyses of animals treated from L4 with RNAi against *fasn-1* (I,K), *pod-2* (J), and co-treated with 4.5 mM phenformin (I-J) or 50 mM metformin (K). (L) Lifespan analyses of WT or hypomorphic *fasn-1(fr8)* mutation bearing animals treated with 4.5 mM phenformin from L4. (M) Representative micrographs of conditional FASN-1 depleted animals imaged 5 days after L1 treatment with 4.5 mM phenformin or 5 µM 5-PH-IAA. Quantification is the % of animals reaching egg-laying adulthood 72 hour after indicated treatments (n = 4 biological replicates). (N) GFP quantification of GFP::AID*::3xFLAG::FASN-1 animals treated with a *fasn-1* RNAi dose dilution curve series shown compared to EV control. Data represent the mean ± SEM of at least 10 animals aggregated across 2 biologically independent experiments. (O) Mean lifespan of WT *C. elegans* treated with 4.5 mM phenformin and an RNAi dose dilution curve of *fasn-1* RNAi mixed with EV RNAi. Lifespans are representative of 3 biological replicates (Table S3). Data represent the mean ± SEM. Imaging data are representative of at least 30 animals aggregated from 3 biologically independent experiments per condition, unless otherwise noted. Significance was assessed using (B) clusterProfiler overrepresentation tests, (C) paired t-tests, (D-F) DESeq2 likelihood ratio tests, (G, M) two-way ANOVA followed by correction for multiple comparisons, (I-L) log rank analysis, and (H, N-O) one-way ANOVA followed by correction for multiple comparisons. ns p > 0.05, * p < 0.05, ** p < 0.01, *** p < 0.001, **** p < 0.0001.

Previous studies proposed that attenuating cytosolic translation unidirectionally impacts mitochondrial translation^46^. However, RNAi knockdown of cytosolic, but not mitochondrial, ribosomal subunits phenocopied and acted non-additively with metformin-mediated growth inhibition (Figure S1A-B). Moreover, polysome profiling of *C. elegans* treated with a longevity-promoting dose of the biguanide phenformin revealed that phenformin significantly attenuated the polysome-to-monosome ratio, indicating decreased mRNA translation initiation (Figure 1C). Consistently, we found that biguanides extend lifespan in a manner that operates non-additively with loss of canonical cytosolic mRNA translation inhibition paradigms (Figure S1C-F)^32,34,47^.

To assess the impact of biguanides on cytosolic mRNA translation, we performed RNA-sequencing on the total, polysomal and monosomal RNA fractions of vehicle and phenformin treated animals. Principal component analysis revealed distinct striation of samples on PC1 by treatment and PC2 by ribosomal fraction, suggesting robust fidelity in separation of actively translating ribosomal pools (Figure S1G). In line with our polysome-to-monosome analysis, we found that polysome associated mRNAs are globally depleted upon phenformin treatment (Figure S1H). Using protein subunit complex analysis^48^, we observed significant reduction in the translation efficiency (TE) of cytoplasmic ribosome, eEF1 translation elongation complex, and electron transport chain (ETC) complex IV components (Figure S1I, Table S2). We consequently observed that phenformin reduced the TE of cytosolic, but not mitochondrial, ribosomal subunits (Figure S1J). Interestingly, we observed that phenformin preferentially reduced the TE of ETC complex IV subunits, consistent with studies suggesting ETC complex IV as the molecular target of the drug (Figure S1I)^18^. We thus hypothesized that reduced production of complex IV subunits would mimic biguanide-mediated lifespan extension. In support of this, post-developmental RNAi knockdown of *cox-7C*, a complex IV subunit displaying the greatest reduction in translation efficiency upon phenformin treatment (ΔTE = - 0.71, adjusted *p* value = 0.0031) extended lifespan non-additively with phenformin treatment (Figure S1K, for all lifespan statistics see Table S3). Combined, these data suggest that biguanides attenuate cytoplasmic mRNA translation and inhibit cytoplasmic ribosomal and ETC complex IV TE to partially exert its favorable pro-longevity effects in *C. elegans*.

### Biguanides preferentially protect the translation efficiency of mRNAs encoding fatty acid biosynthetic enzymes fasn-1 and pod-2

Our polysome profiling identified 26 transcripts with increased TE upon phenformin treatment, suggesting that their protected polysome engagement in the face of attenuated translation may be required to promote biguanide action (Figure 1D-E). Systematic RNAi lifespan determination of these 26 genes in RNAi-hypersensitive *eri-1(mg366)* animals identified two factors, *pod-2/CeACC* and *fasn-1/CeFASN*, that abrogated phenformin-mediated lifespan extension, surpassing factors known to be epistatic to biguanide action in aging (*npp-3*, *npp-21*, *fard-1*) (Figure S1L)^14,15^. Unlike downstream fatty acid desaturases (*fat-3, fat-4*), *fasn-1* and *pod-2* transcripts remain translationally protected, suggesting a concerted effort to sustain fatty acid biosynthesis during attenuation of global mRNA translation (Figure 1F). We validated the translational protection of *fasn-1* and *pod-2*, quantifying endogenous GFP-tagged levels of FASN and POD-2 relative to RPL-31, RPL-4, COX-6A, and FAT-3, predicted in our polysome profiling studies to have reduced TE with phenformin treatment by L4 of development (Figure 1G)^49^. We found that endogenous levels of POD-2 are significantly increased upon administration of metformin or phenformin to adult day 3, suggesting that early translational protection of POD-2 manifests later in life with increased protein levels (Figure 1H).

### Biguanides require fasn-1 and pod-2 to prevent accelerated death outcomes in C. elegans

We assessed the impact of post-developmental *fasn-1* or *pod-2* inactivation on biguanide-mediated lifespan extension. We first validated that post-developmental RNAi robustly depletes endogenous protein levels of FASN-1 and POD-2, and, in the case of *fasn-1*, has similar transcript knockdown efficiency regardless of phenformin treatment (Figure S1M-O). Compellingly, post-developmental RNAi knockdown of either *fasn-1* or *pod-2* not only completely abrogated phenformin-mediated lifespan extension, but synergistically reduced survival, resulting in an accelerated death phenotype we term **BIG**uanide-induced **S**ynergistic **A**cceleration of **D**eath (**BIG-SAD**) (Figure 1I-J). Post-developmental *fasn-1* RNAi induced BIG-SAD with metformin treatment, suggesting that both biguanides require fatty acid biosynthesis to prevent toxicity (Figure 1K). We verified that BIG-SAD is not confounded by chemical sterility agents, as lifespans performed without the use of the thymidylate synthase inhibitor 5-fluoro-2′-deoxyuridine (FUdR) showed similar rates of accelerated death (Figure S1P). Treatment of animals with phenformin from either the L1 or L4 stage did not alter BIG-SAD (Figure S1P). To verify that BIG-SAD is not restricted to RNAi-based approaches, we leveraged nematodes bearing a hypomorphic G723A mutation in *fasn-1* (*fasn-1(fr8)*), finding that *fasn-1(fr8)* animals treated post-developmentally with phenformin results in BIG-SAD, independent of indirect effects on bacterial metabolism (Figure 1L, S1Q)^11,13,50,51^. We ruled out enhanced drug uptake as a mechanism for BIG-SAD by performing LC-MS metabolomics to quantify absolute levels of phenformin, observing no significant increase in phenformin levels with *fasn-1* RNAi at any dose of phenformin tested (Figure S1R). A 0.5 mM phenformin dose was sufficient to accelerate the death of *fasn-1* RNAi treated animals by ∼50%, without a concomitant ∼50% increase in phenformin uptake in our LC-MS analysis, as would be expected if enhanced uptake was the sole mechanism by which BIG-SAD is mediated (Figure S1S). To further corroborate that BIG-SAD was the consequence of *fasn-1* loss of function, we constructed a conditional depletion model of FASN-1 using the auxin-induced degron (AID) system, observing that auxin-induced degradation of somatic FASN-1 from L1 results in a highly penetrant arrest phenotype when combined with phenformin treatment, akin to BIG-SAD (Figure 1M).

Finally, we hypothesized that if translational protection of FASN-1 production is required to prevent BIG-SAD, then titrating the amount of FASN-1 available may modulate its onset. Indeed, titrating the dosage of *fasn-1* RNAi by varying its dilution with empty vector RNAi was sufficient to linearly increase the average lifespan of phenformin co-treated animals; a 1:1000 dilution of *fasn-1* RNAi, which only moderately impacts *fasn-1* expression, was sufficient to restore phenformin-mediated lifespan extension (Figure 1N-O). In aggregate, these data suggest that protecting the translational efficiency of *fasn-1* and *pod-2* transcripts during biguanide treatment is necessary to prevent accelerated death and enable biguanide-induced lifespan extension.

### EGL45/CeEIF3A and EGL-45/CeEIF3B mediate the translational protection of pod-2 transcripts during phenformin treatment

We hypothesized that specific mRNA translation factors may coordinate selective mRNA translation and geroprotective action following biguanide treatment. To test this, we screened 48 translation initiation, 6 elongation, and 2 termination factors for post-developmental RNAi knockdowns that suppressed the activation of acyl-coenzyme A synthase ACS-2, a robust, established biomarker for the pro-longevity action of biguanides (Figure 2A)^13,14^. This identified two genes, *egl-45*/*Ce*EIF3A and *eif-3.B*/*Ce*EIF3B, conserved members of the eIF3 translation initiation complex, as necessary for biguanide-prompted activation of the transcriptional reporter *acs-2p::GFP*, mirroring the effects of *pod-2* RNAi (Figure 2B-C, S2A). Consistent with their increased activity, *egl-45* and *eif-3.B* transcripts are significantly upregulated upon phenformin treatment in a manner upstream of *fasn-1* inactivation (Figure 2D, Table S4). Importantly, RNAi of *eif-3.B* and *egl-45* does not mitigate biguanide-induced activation of the orthogonal transcriptional reporter *bigr-1p::bigr-1::mRFP3*, suggesting they do not act as transgene suppressors (Figure S2B)^13,14^. Furthermore, we observed that inactivation of *egl-45* or *eif-3.B* suppressed biguanide-induced mobilization of somatic neutral lipid stores and growth restriction, further suggesting that EGL-45 and EIF-3.B are mechanistically linked to the known metabolic outcomes of biguanide action (Figure 2E-G)^13,15^.

**Figure 2.**
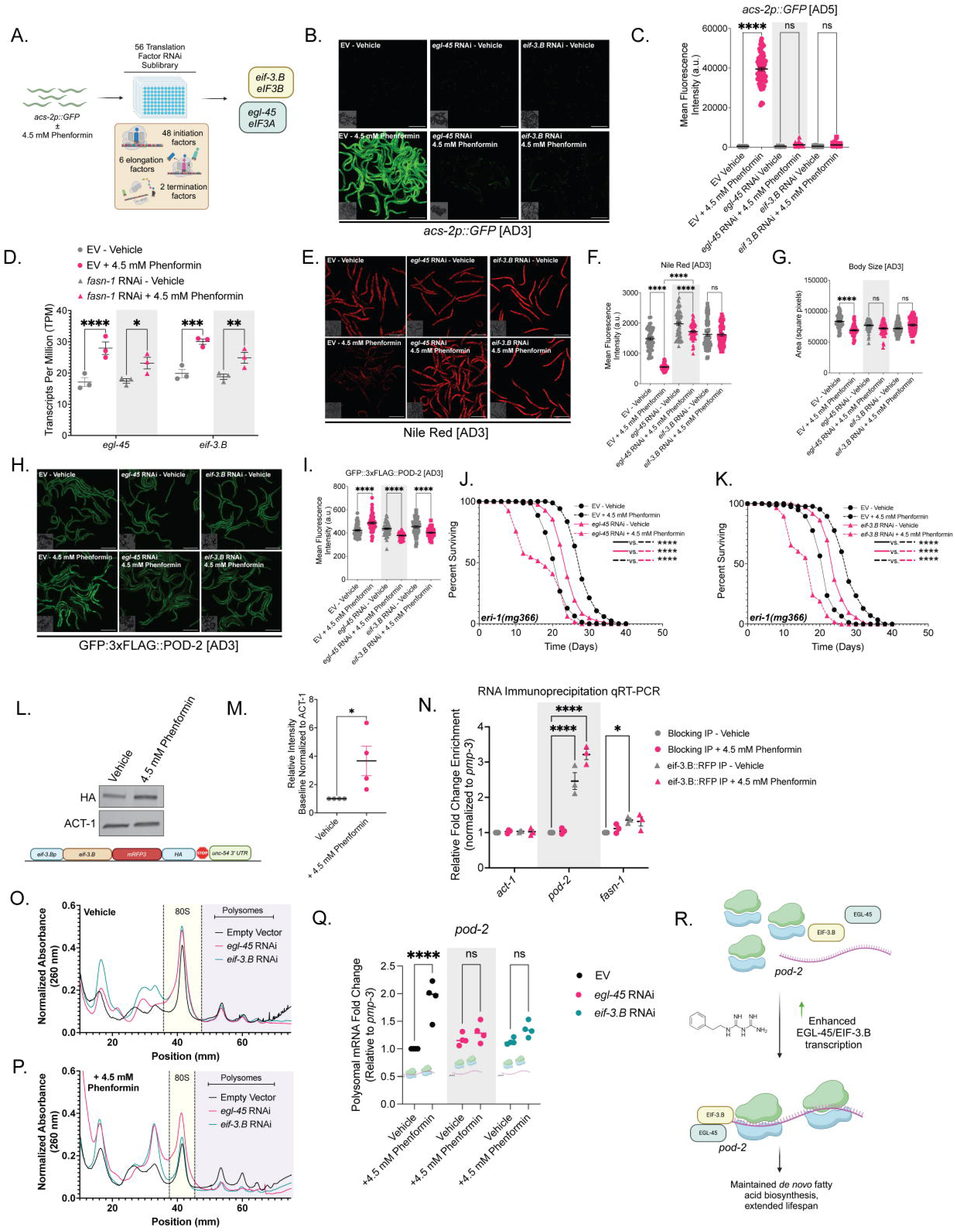
Translation efficiency of *pod-2* is coordinated by EIF3 subunit factors EGL-45/CeEIF3A and EIF-3.B/CeEIF3B during biguanide response. (A) Schematic of RNAi strategy used to identify two translation factors (*egl-45* and *eif-3.B*) that suppressed biguanide mediated induction of *acs-2p::GFP*. (B-C) Representative micrographs (B) and GFP (C) quantification of *acs-2p::GFP* levels in adult day 5 animals treated from L4 with *eif-3.B* or *egl-45* RNAi and 4.5 mM phenformin. Scale bar indicates 500 µM. (D) Normalized transcripts per million (TPM) of *egl-45* and *eif-3.B* with *fasn-1* RNAi or 4.5 mM phenformin treatment. Note that the vehicle measurements in 2D are identical to those in 6G, as these RNA-sequencing experiments were performed in tandem (n = 3 biological replicates). (E-G) Representative micrographs (E) and GFP (F) quantification of Nile-Red stained neutral lipid stores in adult day 3 animals treated from L4 with *eif-3.B* or *egl-45* RNAi and 4.5 mM phenformin. Body size area for each animal was assessed (G). Nile Red MFI was normalized to animal body area. Note that the EV measurements in 2F are identical to those shown in 3E as these quantifications were performed in parallel. (H-I) Representative micrographs (H) and fluorescence GFP (I) quantification of GFP::3xFLAG::POD-2 in adult day 3 animals treated from L4 with *eif-3.B* or *egl-45* RNAi, and 4.5 mM phenformin. Note that the EV measurements in 2I are identical to those shown in S1N, as these quantifications were performed in parallel. (J-K) Representative lifespan analyses of RNAi hypersensitive *eri-1(mg366)* animals treated from L4 with *eif-3.B* or *egl-45* RNAi, and 4.5 mM phenformin. (L) Representative western blot of *eif-3.Bp::eif-3.B::mRFP3::HA::unc-54 3’ UTR* array expressing animals treated with 4.5 mM phenformin from L1 to L4/YA. Lower panel displays the array structure schematic. (M) Quantification of western blots for induction of *eif-3.B* extrachromosomal array (n = 3 biological replicates). (N) RNA immunoprecipitation coupled with qRT-PCR (RIP-qRT-PCR) of *eif-3.Bp::eif-3.B::mRFP3::HA::unc-54* 3’ UTR array expressing animals treated with phenformin from L1 to L4/YA. Enrichment of *act-1*, *pod-2* and *fasn-1* is normalized to *pmp-3* and is compared relative to RNA isolated from IP of blocking beads in vehicle treated animals (n = 3 biological replicates). (O-P) Representative polysome profiles from WT *C. elegans* treated with *egl-45* RNAi or *eif-3.B* RNAi alone (O) or with 4.5 mM phenformin treatment (P) (n = 4 biological replicates). (Q) qRT-PCR of *pod-2* from pooled polysomal fractions of WT *C. elegans* treated as shown in (O-P) (n = 4 biological replicates). (R) Schematic interpretation of all results in Figure 2. Lifespans are representative of 3 biological replicates (Table S3). Data represent the mean ± SEM. Imaging data are representative of at least 30 animals aggregated from 3 biologically independent experiments per condition, unless otherwise noted. Scale bars indicate 500 µM. Significance was assessed by (D) DESeq2 Wald tests with Benjamini-Hochberg FDR correction, (C, F, G, I, N, Q) two-way ANOVA followed by correction for multiple comparisons, (J-K) log rank analyses, and (M) Student’s t-tests. ns p > 0.05, * p < 0.05, ** p < 0.01, *** p < 0.001, **** p < 0.0001.

We next quantitatively examined EGL-45 and EIF-3.B translational protection of *pod-2* or *fasn-1*. Quantitative fluorescence imaging of endogenously tagged POD-2 and FASN-1 CRISPR knock-in animals revealed that inactivation of *egl-45* or *eif-3.B* prevented the biguanide-mediated induction of POD-2, but not FASN-1, levels (Figure 2H-I, S2C). Given that *pod-2* RNAi induces BIG-SAD, we hypothesized that loss of factors that protect POD-2 translation upon biguanide treatment should mimic this response. Leveraging *eri-1(mg366)* animals, we found that post-developmental inactivation of *egl-45* or *eif-3.B* in combination with biguanide treatment results in an accelerated death curve, in contrast to the extended lifespan observed with RNAi inactivation of *egl-45* or *eif-3.B* alone (Figure 2J-K)^33^. We note that these accelerated death phenotypes as not as severe as those seen with direct inactivation of fatty acid biosynthesis, likely due to EGL-45/EIF-3.B selectively augmenting *pod-2* translation during biguanide treatment without affecting its basal levels (Figure 2H-K). Combined, these data suggest that *egl-45* and *eif-3.B* preserve translation of *pod-2* transcripts upon biguanide treatment in part to prevent BIG-SAD.

Emerging studies have shown that human EIF3A and EIF3B can selectively regulate polysome occupancy and translation initiation of specific mRNAs in a sequence-specific manner^52^. To explore this, we constructed transgenic nematodes overexpressing an EIF-3.B::mRFP3::HA fusion protein, validating increased expression upon phenformin treatment (Figure 2L-M). RNA immunoprecipitation qRT-PCR of *pod-2* transcripts indicated a ∼2-3 fold enrichment of *pod-2* transcripts associated with EIF-3.B::mRFP3 with either vehicle or phenformin treatment, and a significant but marginal enrichment of *fasn-1* transcripts, relative to *act-1* and *pmp-3* reference controls (Figure 2N, S2D), suggesting that EIF-3.B can bind directly to *pod-2* transcripts. Polysome profiling analyses indicated that inactivation of *eif-3.B* and *egl-45* combined with phenformin treatment results in a synthetic collapse of polysome occupancy, suggesting that presence of these two factors are required to maintain translation on polysomes with biguanide treatment (Figure 2O-P, S2E). qRT-PCR analysis of isolated polysomal fractions revealed that knockdown of *egl-45* and *eif-3.B* suppressed the enhanced polysome occupancy of *pod-2*, but not *fasn-1*, transcripts with phenformin treatment (Figure 2Q, S2F). Intriguingly, inactivation of *egl-45* and *eif-3.B* also partially reversed the decreased polysome occupancy of *fat-3* and *fat-4* transcripts upon biguanide treatment, suggesting that EGL-45 and EIF-3.B are in part responsible for polysomal selectivity upon exposure to phenformin in *C. elegans* (Figure S2G-H). Thus, translation factors EGL-45 and EIF-3.B are transcriptionally elevated by biguanide treatment, bind to *pod-2* transcripts to maintain polysome occupancy, and enable biguanide-mediated pro-longevity outcomes (Figure 2R).

### FASN-1 and POD-2 do not buffer BIG-SAD through production of a lipid product or modulation of lipid stores

To confirm increased POD-2 and FASN-1 activity upon biguanide treatment, we used gas chromatography-mass spectrometry (GC-MS) to analyze fatty acids from phenformin-treated nematodes (Table S5). Enhanced accumulation of palmitate and stearate was observed relative to controls, indicating increased fatty acid biosynthesis (Figure 3A-B)^51,53^. This effect was independent of whether the *E. coli* food source was live or inactivated, suggesting that biguanides directly induce palmitate and stearate production in the nematode (Figure 3A-B). Triglycerides (TAG) did not further decrease with combined *fasn-1* RNAi and phenformin treatment, implying no additional mobilization of lipid stores (Figure 3C-E). Additional GC-MS analyses revealed equivalent effects of biguanides on fatty acid composition regardless of *fasn-1* RNAi in either triglyceride or phospholipid fractions, indicating a lack of specific fatty acid remodeling (Figure 3F-G). Previous studies suggested that animals bearing loss of lipid biosynthetic transcription factor SBP-1 or mediator subunit MDT-15 result in accelerated death when administered compounds that alter metabolic rewiring, including dietary glucose treatment^54^. In contrast, we found that hypomorphic *sbp-1(ep79)* animals do not mimic the BIG-SAD observed with *fasn-1* deficiency (Figure S3A-B). Null mutations in *mdt-15* did accelerate the death of biguanide treated animals, but *fasn-1* RNAi accelerated their death even further, and a gain-of-function allele in *mdt-15* failed to rescue *fasn-1* RNAi induced BIG-SAD (Figure S3C-D)^55^.

**Figure 3.**
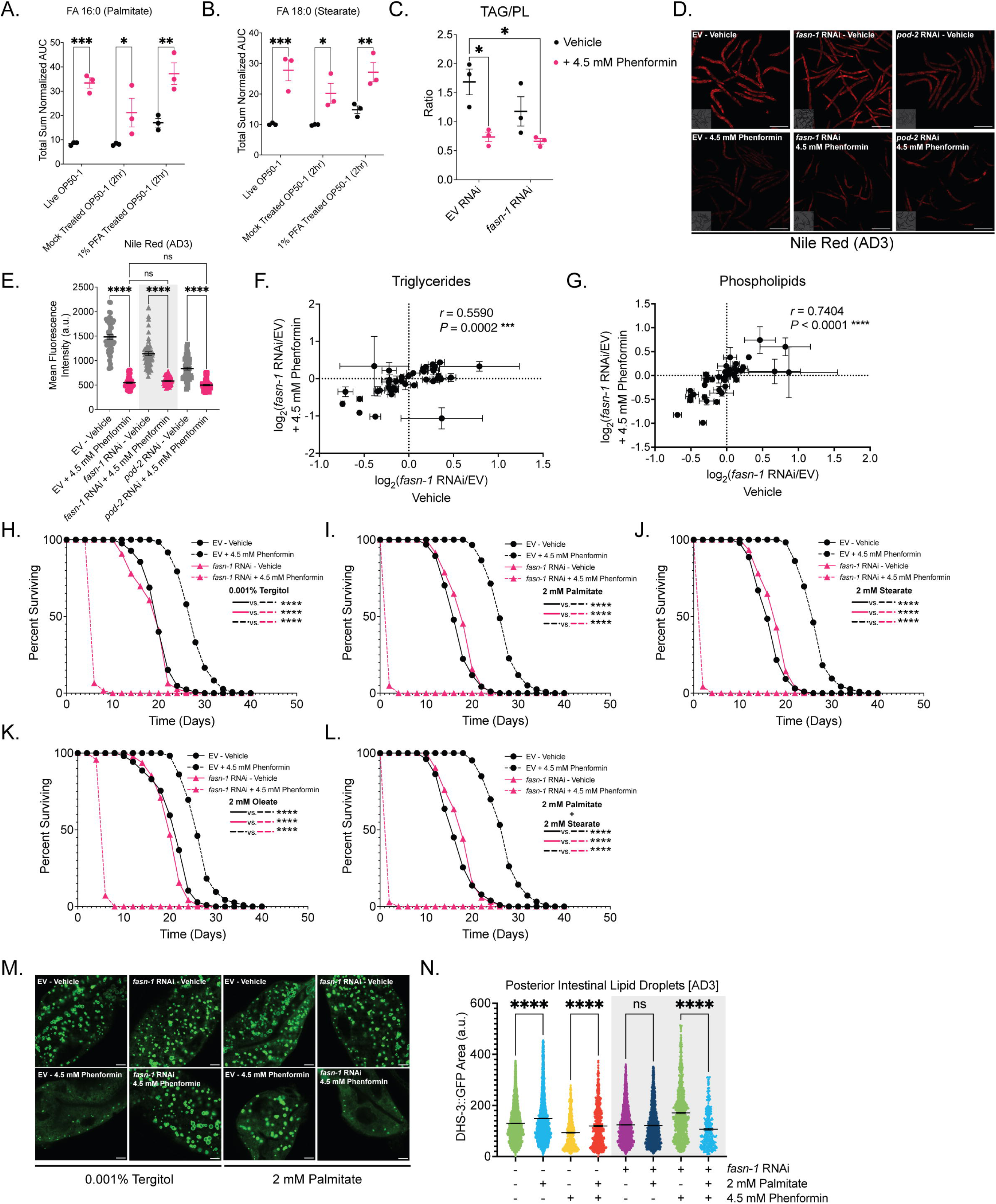
*fasn-1*/*pod-2* RNAi induced BIG-SAD is not mediated by shifts in lipid production or utilization. (A-B) GC-MS analysis of palmitate (A) or stearate (B) fatty acid methyl esters derivatized from animals treated with 4.5 mM phenformin to AD1 on the indicated *E. coli* food sources (n = 3 biological replicates). (C) Triglycerides (TAG) normalized to phospholipid (PL) stores in WT animals treated from L4 to AD3 with *fasn-1* RNAi or 4.5 mM phenformin (n = 3 biological replicates). (D-E) Representative micrographs (D) and GFP (E) quantification of Nile Red-stained neutral lipid stores in WT *C. elegans* treated from L4 to AD3 with *fasn-1* or *pod-2* RNAi or 4.5 mM phenformin. Nile Red MFI was normalized to animal body area. Note that the EV measurements in 3E are identical to those shown in 2F, as these quantifications were performed in parallel. Scale bar indicates 500 µM. (F-G) Scatterplots of TAG (F) or PL (G) associated fatty acids in WT *C. elegans* treated as shown in (C). (n = 3 biological replicates). (H-L) Lifespan analyses of WT animals treated with *fasn-1* RNAi or 4.5 mM phenformin on plates supplemented with 0.001% tergitol control (H), 2 mM palmitate (I), 2 mM stearate (J), 2 mM oleate (K), or both 2 mM palmitate and 2 mM stearate (L). (M-N) Representative micrographs (M) and quantification (N) of DHS-3::GFP lipid droplet area in posterior intestinal cells treated with *fasn-1* RNAi, 4.5 mM phenformin, or 2mM palmitate from L4 to AD3. Note that the EV/*fasn-1* ± 2 mM palmitate measurements in 3N are identical to those shown in S6M, as these quantifications were performed in parallel. Data represent the mean ± SEM of lipid droplets combined from at least 10 animals per condition aggregated from 3 biologically independent replicates. Scale bar indicates 50 µM. Lifespans are representative of 3 biological replicates (Table S3). Data represent the mean ± SEM. Imaging data are representative of at least 30 animals aggregated from 3 biologically independent experiments per condition, unless otherwise noted. Significance was assessed by (A-C, E, N) two-way ANOVA followed by correction for multiple comparisons, (H-L) log rank analyses, and (F-G) Pearson’s correlation coefficient analysis. ns p > 0.05, * p < 0.05, ** p < 0.01, *** p < 0.001, **** p < 0.0001.

Given that palmitate is the primary product of *de novo* fatty acid biosynthesis, we tested whether its loss was the causal factor driving *fasn-1*/*pod-2* RNAi-induced BIG-SAD. Supplementation with elevated concentrations of palmitate, (FA C16:0), stearate (FA C18:0), oleate (FA C18:1), or a combination of palmitate and stearate failed to rescue the BIG-SAD; in the case of palmitate and/or stearate treatment, lipid supplementation accelerated the onset of death even further (Figure 3H-L)^56^. These results suggested that it is not the product of fatty acid biosynthesis, but rather the process itself that is required to ensure viability upon treatment with biguanides. Accordingly, we hypothesized that exogenous FA supplementation exaggerated the BIG-SAD due to end-product inhibition of residual fatty acid biosynthetic capacity. Lipid droplet (LD) analysis revealed that palmitate supplementation enhanced LD size in vehicle and phenformin treated animals, but synergistically reduced LD number and size with combined *fasn-1* RNAi and phenformin treatment, corroborating our hypothesis (Figure 3M-N). Combined, these results suggest that *fasn-1*/*pod-2* RNAi induced-BIG SAD is not solely due to the lack of production of a major fatty acid product.

### Depletion of FASN-1 induces NADPH reductive stress and synergistically enhances biguanide-mediated NADH and glutathione reductive stress

Fatty acid biosynthesis is the major consumer of NADPH, utilizing 14 moles of NADPH per mole of palmitate produced (Figure 4A)^57,58^. Gene module association analyses revealed that fatty acid synthase activity positively correlates with oxidation-reduction process gene expression (Figure S4A)^59^. Given that palmitate supplementation was insufficient to prevent BIG-SAD, we hypothesized that the ability of *fasn-1* to consume excess NADPH reducing equivalents, rather than lipogenesis *per se*, is mobilized upon biguanide treatment. Indeed, LC-MS analysis revealed that phenformin treatment significantly increased NADPH/NADP+, suggesting that phenformin alone may prime the animals in a NADPH reductive state (Figure 4B). Inactivation of *fasn-1* further exaggerated the buildup of NADPH/NADP+ at any concentration of drug administered, as expected if a principal role of FASN-1/CeFASN is to consume excessively produced NADPH and mitigate reductive stress (Figure 4B).

**Figure 4.**
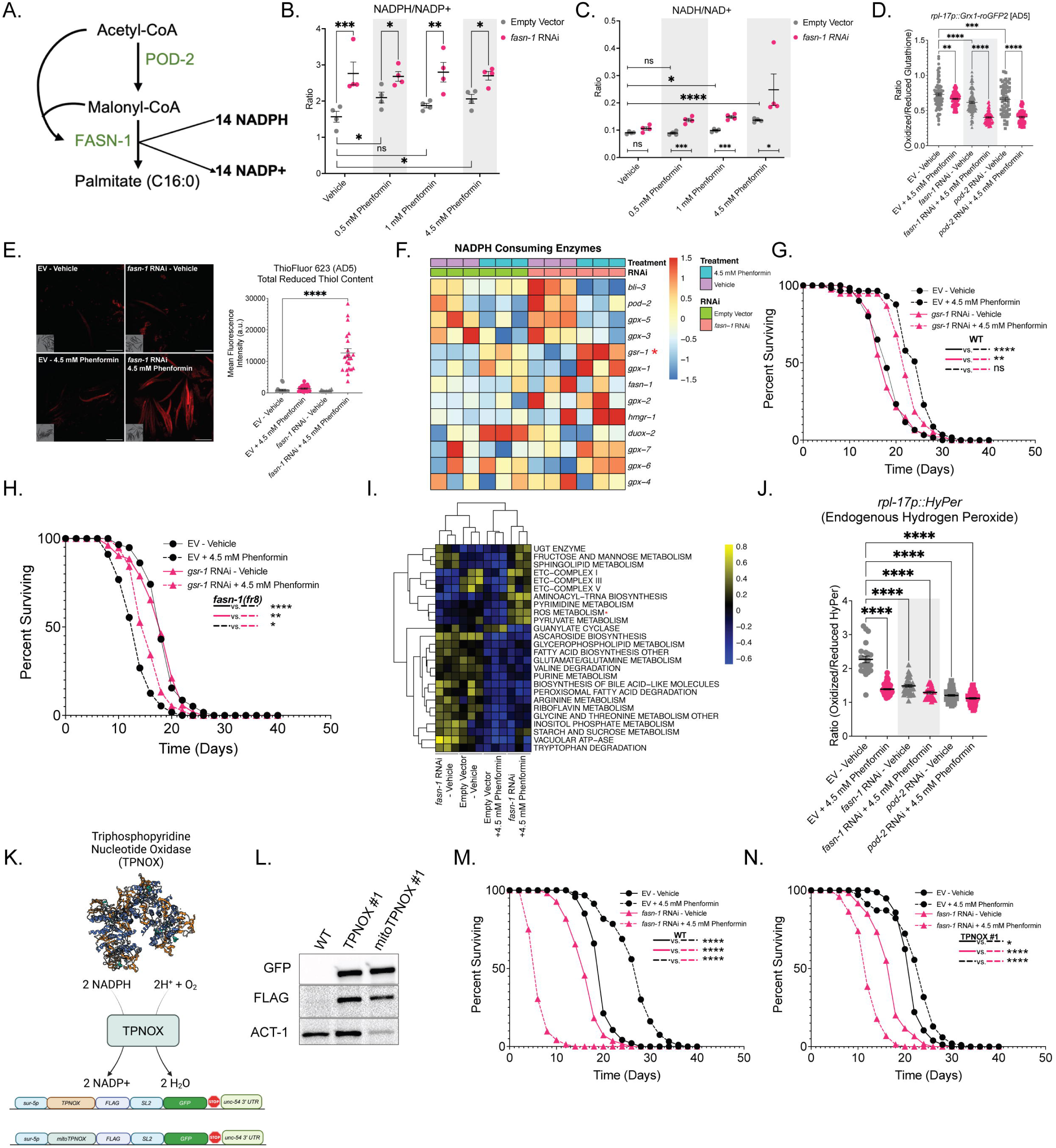
Inactivation of *fasn-1/pod-2* combinatorically enhances reductive stress with biguanide treatment. (A) Schematic of *de novo* lipogenesis indicating estimated molar consumption of NADPH per mole of palmitate produced. (B-C) LC-MS analysis of WT animals treated with *fasn-1* RNAi and varying doses of phenformin from L4 to AD3 with measurement of (B) NADPH/NADP+ and (C) NADH/NAD+. Note that the vehicle measurements in 4B and 4C are identical to those shown in 6I and 6J, respectively, as these quantifications were performed in parallel (n = 4 biological replicates). (D) Ratiometric expression of *rpl-17p::Grx1-roGFP2* animals treated with *fasn-1*/*pod-2* RNAi or 4.5 mM phenformin from L4 to AD5. (E) Representative micrographs and GFP quantification of WT animals treated with *fasn-1* RNAi or 4.5 mM phenformin from L4 to AD5, and then subsequently stained with reduced thiol dye ThioFluor 623 for 1 hour prior to imaging. Data represent the mean ± SEM of at least 20 animals per condition aggregated from 3 biologically independent experiments. Scale bar indicates 500 µM. (F) Z-score normalized expression of genes known to encode primary metabolic consumers of NADPH in their activity (n = 3 biological replicates). (G-H) Lifespan analyses of WT (L) and *fasn-1(fr8)* (M) animals treated with *gsr-1* RNAi or 4.5 mM phenformin. (I) Gene Set Variation Analysis (GSVA) pathway activity scores for metabolic gene sets extracted from WormClust^58^ in WT animals treated with *fasn-1* RNAi or 4.5 mM phenformin (BH adjusted *p* value < 0.05.) (J) Ratiometric expression of *rpl-17p::HyPer* animals treated with *fasn-1*/*pod-2* RNAi and with 4.5 mM phenformin from L4 to AD3. (K) Cartoon schematic of rationally engineered Triphosphopyridine Nucleotide Oxidase (TPNOX) quenching NADPH accumulation. (L). Representative western blot of WT animals bearing *sur-5p::TPNOX::FLAG::SL2::GFP::unc-54 3’ UTR* and *sur-5p::mitoTPNOX::FLAG::SL2::GFP::unc-54 3’ UTR* arrays at AD1. (M-N) Representative lifespan analyses of WT N2 (M) and *sur-5p::TPNOX::FLAG::SL2::GFP::unc-54 3’ UTR* overexpressing (N) animals treated with *fasn-1* RNAi or 4.5 mM phenformin. Lifespans are representative of 3 biological replicates (Table S3). Data represent the mean ± SEM. Imaging data are representative of at least 30 animals aggregated from 3 biologically independent experiments per condition, unless otherwise noted. Significance was assessed by (B-E, J) two-way ANOVA followed by correction for multiple comparisons, (G-H, M-N) log rank analyses, and (I) empirical Bayes tests with Benjamini-Hochberg correction using limma(). ns p > 0.05, * p < 0.05, ** p < 0.01, *** p < 0.001, **** p < 0.0001.

Given the coupled nature of NADPH-GSH and GSH-NADH pools, we hypothesized that a buildup of NADPH reducing equivalents could result in electron spillover, shuffling excess reducing equivalents across multiple redox pools^29^. Indeed, while biguanide treatment alone significantly increased the NADH/NAD+ ratio, consistent with established reports of biguanides as ETC inhibitors^22^, combinatorial *fasn-1* RNAi significantly increased the NADH/NAD+ ratio, suggestive of enhanced NADH reductive stress (Figure 4C). We also observed increased reduced glutathione pools, as measured by an *in vivo* GSH/GSSH redox response reporter and endogenous reduced thiol staining (Figure 4D-E)^60,61^. Affirming the link between elevated NADPH and GSH production, we observed an increase in *gsr-1* mRNA, the sole *C. elegans* glutathione reductase, with combinatorial *fasn-1* RNAi and phenformin treatment (Figure 4F). RNAi of *gsr-1* partially delayed the onset of BIG-SAD in *fasn-1(fr8)* animals (Figure 4G-H). We identified shared evolutionary history between *fasn-1* and *gsr-1* using clustering by inferred models of evolution (CLIME) analysis, suggesting an ancient, conserved compensatory link between fatty acid synthase activity and glutathione reduction (Figure S4B)^62^. Excess reducing equivalent production has been linked to an elevation in reactive oxygen species (ROS) scavenging systems^63,64^. Consistently, we observe enhanced expression of ROS metabolic genes, including elevation of catalases (*ctl-1*/*ctl-2*) and superoxide dismutases (*sod-1*/*sod-2*), and a decrease in overall endogenous hydrogen peroxide pools, with combinatorial *fasn-1* RNAi and phenformin treatment (Figure 4I-J, S4C)^61^.

Finally, to determine whether quenching NADPH accumulation would resolve the BIG-SAD, we overexpressed cytosolic or mitochondrial triphosphopyridine nucleotide oxidase (TPNOX)^65^, a genetically encoded tool which oxidizes NADPH to NADP+ (Figure 4K-L). Indeed, expression of cytoplasmic, but not mitochondrial, TPNOX was sufficient to delay the onset of *fasn-1*/*pod-2* RNAi induced BIG-SAD (Figure 4M-N, S4D-G). We additionally observed that overexpression of *LbNOX*, a genetically encoded NADH oxidase, was not sufficient to mitigate the BIG-SAD (Figure S4H-J)^66^. In aggregate, these data suggest that elevated reductive stress due to excess NADPH accumulation is causally linked to biguanide-induced toxicity, and that the enzymatic process of fatty acid biosynthesis acts as the major sink for reducing equivalents to prevent BIG-SAD and enable pro-longevity outcomes.

### Biguanide longevity regulators AAK-2/CeAMPK and SKN-1/CeNRF tune the onset and severity of reductive stress toxicity

Combinatorial *fasn-1* RNAi and phenformin treatment caused a dose-dependent collapse in adenylate energy charge and decreased TCA metabolites, reflecting reduced ETC activity and mitochondrial energetics (Figure 5A, S5A)^67^. RNA-sequencing revealed downregulation of genes linked to adenosine monophosphate-activated protein kinase (AAK-2/CeAMPK) activity in combined *fasn-1* RNAi and phenformin treated animals, indicating a paradoxical collapse of AMPK activity (Figure 5B). Indeed, combinatorial treatment with *fasn-1* or *pod-2* RNAi and phenformin treatment significantly reduced phospho-AAK-2^Thr172^ levels, suggesting an inability to compensate for perturbed ATP energy homeostasis (Figure 5C-D). Consistently, we noted that null mutation in *aak-2* accelerated the BIG-SAD, and, conversely, overexpression of a constitutively activated form of AAK-2 delayed the BIG-SAD (Figure 5E-G)^68^. Thus, biguanides induce mild ATP exhaustion that is exaggerated by concomitant *fasn-1* knockdown and is associated with accelerated death. If correct, it follows that ATP exhausting insults would require *de novo* fatty acid biosynthesis to buffer early death responses. Indeed, treatment of *C. elegans* with oligomycin, a potent inhibitor of Complex V that reduces ATP synthesis and promotes NADH accumulation, mirrors *fasn-1*/*pod-2* RNAi induced BIG-SAD (Figure 5H, S5B)^67^. We additionally observed that hypomorphic, longevity-promoting mutations in coenzyme-Q biosynthesis (*clk-1(qm30))* or ETC complex III activity (*isp-1(qm150)*) display significantly suppressed lifespans in response to *fasn-1* RNAi, corroborating that interventions which reduce ETC activity require *fasn-1* to buffer early death responses that accompany increased reducing equivalents and associated ATP exhaustion (Figure S5C).

**Figure 5.**
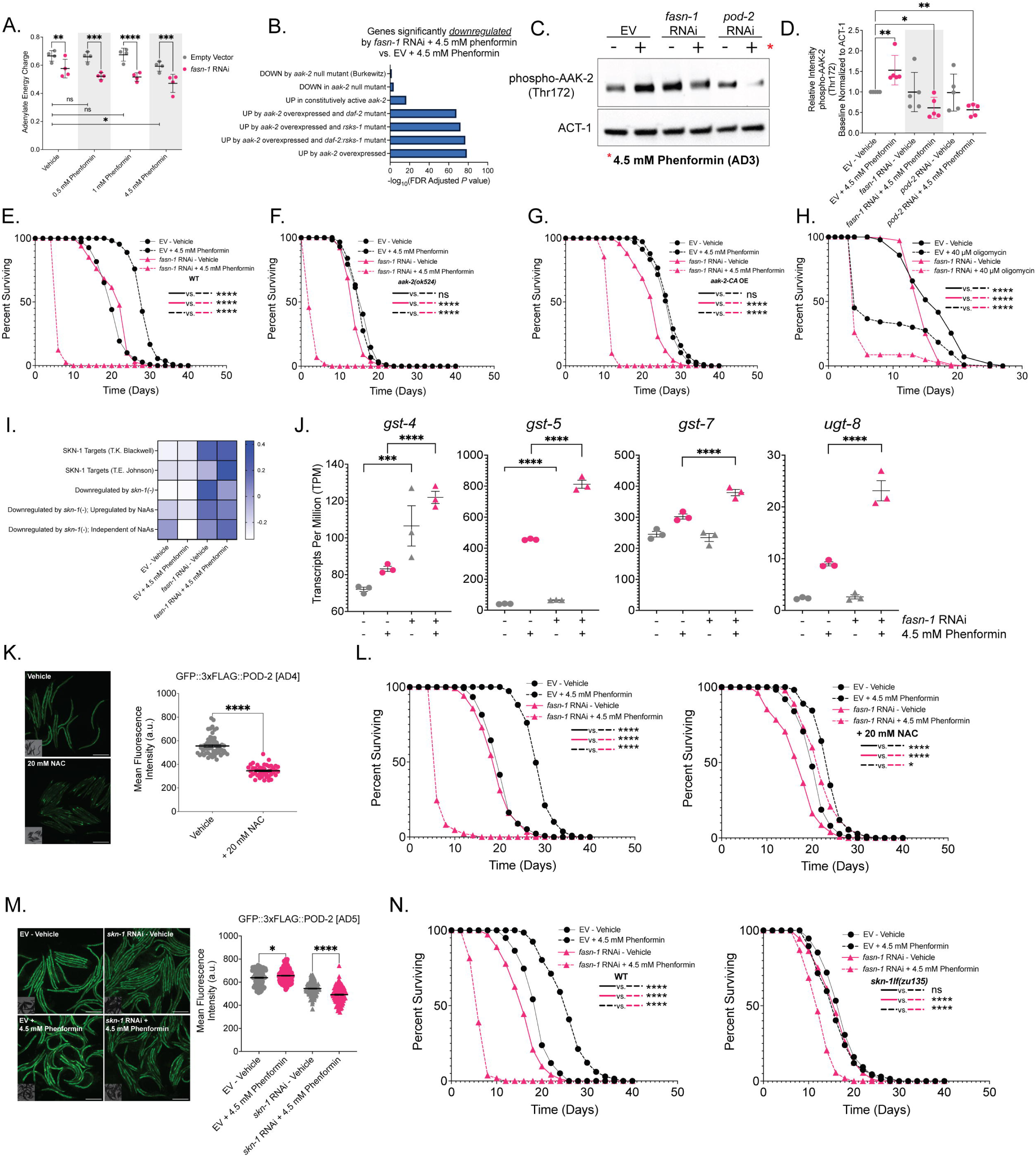
Manipulation of redox and ATP homeostasis fine tunes biguanide-induced toxicity responses. (A) LC-MS analysis of WT *C. elegans* treated with *fasn-1* RNAi and varying doses of phenformin from L4 to AD3 with measurement of adenylate energy charge (defined as the ratio of (ATP + 0.5 * ADP)/(ATP+ADP+AMP). Note that the vehicle measurements in 5A are identical to those shown in S6O, as these quantifications were performed in parallel (n = 4 biological replicates). (B) WormExp^109^ enrichment analysis for *C. elegans* mutant genotype gene sets in genes differentially expressed between *fasn-1* RNAi + 4.5 mM phenformin and +4.5 mM phenformin (DESeq2 adjusted *p* value < 0.05.) (C) Representative western blot of phospho-AAK-2^Thr172^ levels in WT animals treated with *fasn-1* RNAi or *pod-2* RNAi with 4.5 mM phenformin from L4 to AD3. (D) Quantification of western blots as described in (C), normalizing phospho-AAK-2^Thr172^ levels to ACT-1 (n = 5 biological replicates). (E-G) Lifespan analyses of WT (E), *aak-2(ok524)* null (F), and *aak-2CA OE* (G) animals treated with *fasn-1* RNAi or 4.5 mM phenformin. (H) Lifespan analyses of WT animals treated with *fasn-1* RNAi or 40 µM oligomycin treatment from L4 (n = 4 biological replicates). (I) Gene Set Variation Analysis (GSVA) pathway activity scores for SKN-1 related gene sets extracted from WormExp^109^ in WT animals treated with *fasn-1* RNAi or 4.5 mM phenformin from L4 to AD3 (BH adjusted *p* value < 0.05). (J) TPM levels for SKN-1 targets *gst-4*, *gst-5*, *gst-7*, and *ugt-8* with *fasn-1* RNAi or 4.5 mM phenformin from L4 to AD3 (n = 3 biological replicates). (K) Representative micrographs and GFP (M) quantification of *GFP::3xFLAG::POD-2* animals treated with 20 mM N-Acetyl Cysteine (NAC) from L4 to AD5. Note that the vehicle measurements in these graphs are identical to those shown in 1H and 6F, as these quantifications were performed in parallel. (L) Representative lifespan analyses of WT animals treated with *fasn-1* RNAi or 4.5 mM phenformin and supplemented with vehicle (left) or 20 mM NAC (right). (M) Representative micrographs and fluorescence quantification of *GFP::3xFLAG::POD-2* animals treated with *skn-1* RNAi or 4.5 mM phenformin from L4 to AD5. Data represent the mean ± SEM of at least 30 animals aggregated across 3 biologically independent experiments. (N) Lifespan analyses of WT (left) or *skn-1(zu135)* null (right) animals treated with *fasn-1* RNAi or 4.5 mM phenformin. Lifespans are representative of 3 biological replicates (Table S3). Data represent the mean ± SEM. Imaging data are representative of at least 30 animals aggregated from 3 biologically independent experiments per condition, unless otherwise noted. Scale bar indicates 500 µM. Significance was assessed by (A, D, M) two-way ANOVA followed by correction for multiple comparisons, (J) DESeq2 Wald tests with Benjamini-Hochberg FDR correction, (E-H, L, N) log rank analyses, and (I) empirical Bayes testing with Benjamini-Hochberg FDR correction using limma(). ns p > 0.05, * p < 0.05, ** p < 0.01, *** p < 0.001, **** p < 0.0001.

RNA-sequencing analyses also revealed hyperactivation of targets for the stress response transcription factor SKN-1/CeNRF, observing that validated antioxidant targets *gst-4*, *gst-5*, *gst-7*, and *ugt-8* are transcriptionally elevated upon combinatorial *fasn-1* RNAi and phenformin treatment relative to phenformin treatment alone (Figure 5I-J)^69^. SKN-1 activity has been implicated previously in the pro-longevity action of both phenformin and metformin, and activation of a mammalian ortholog, NRF2, has been shown to induce NADH reductive stress in a variety of cancers^9,11,14,15,31^. Thus, we hypothesized that FASN-1 may finely tune SKN-1 activation to mediate the balance between pro-longevity and reductive stress associated toxicity outcomes. We observed that inhibiting SKN-1 activity through the antioxidant compound N-acetyl cysteine (NAC)^60,70,71^ alone suppressed endogenous POD-2 levels and nearly completely abolished *fasn-1* RNAi-induced BIG-SAD (Figure 5K-L). Similarly, *skn-1* RNAi prevented biguanide-induced elevation of POD-2 levels, suggesting that limiting SKN-1 activity bypassed the need for enhanced POD-2 production and its subsequent buffering activity (Figure 5M). Concordantly, null mutation in *skn-1lf(zu135)* animals nearly completely abolished *fasn-1*/*pod-2* RNAi induced BIG-SAD (Figure 5N, S5D), whereas gain-of-function mutations in *skn-1gf(lax188)* accelerated the BIG-SAD (Figure S5E-F) Interestingly, both NAC supplementation and *skn-1lf(zu135)* mutation alone suppressed phenformin-mediated lifespan extension, suggesting that the requirement for SKN-1 in the response to biguanides in aging is complex, balancing pro-longevity responses and reductive stress toxicity (Figure 5L, 5N).

### Metabolic interventions that elevate NADPH production require FASN-1 to buffer against reductive stress

Flux balance analyses indicate that conversion of malate into pyruvate through malic enzyme 1 (MEN-1/CeME1), and glucose consumption into the pentose phosphate pathway through activity of GSPD-1/CeG6PD are both major NADPH producers (Figure 6A)^57^ .Thus, we hypothesized that malate or glucose supplementation in nematodes would mirror *fasn-1*/*pod-2* RNAi induced BIG-SAD. Compellingly, exogenous supplementation of malate (10 mM L-malic acid) or glucose (2% (w/v) D-(+)-glucose) mirrored *fasn-1* RNAi induced BIG-SAD, a phenotype not observed with supplementation of other glycolytic or TCA intermediary metabolites (Figure 6B-D, S6A-E). Supplementation with palmitate and stearate failed to rescue malate or glucose-mediated accelerated death, akin to phenformin treatment, despite palmitate supplementation elevating somatic lipid stores and average lipid droplet diameter (Figure S6F-M). We then hypothesized, like phenformin, that malate and glucose supplementation may protect protein synthesis of FASN-1 and/or POD-2 to sustain NADPH buffering capacity. Indeed, malate and glucose supplementation significantly elevated levels of endogenously tagged FASN-1::GFP (Figure 6E), and malate significantly elevated intestinal levels of GFP::3xFLAG::POD-2 (Figure 6F). Glucose supplementation paradoxically reduced levels of GFP::3xFLAG::POD-2, suggesting a more complex regulation of POD-2, or a pleiotropic effect of dietary glucose on POD-2 activity (Figure 6F). Because malate supplementation more closely mimicked BIG-SAD, we tested supplemented animals for expression of *egl-45* and *eif-3.B*, which translationally protected POD-2 expression upon biguanide treatment. Akin to phenformin treatment, and suggesting a conserved mechanism by which POD-2 protein levels were increased, malate supplementation increases expression of *egl-45* and *eif-3.B* transcripts (Figure 6G). We affirmed elevated *de novo* fatty acid biosynthesis of stearate with malate treatment, leveraging a radiolabeled acetate tracing strategy that we validated using *daf-2(e1370)* mutants, previously known to display enhanced *de novo* fatty acid synthesis (Figure 6H, S6N)^72^.

**Figure 6.**
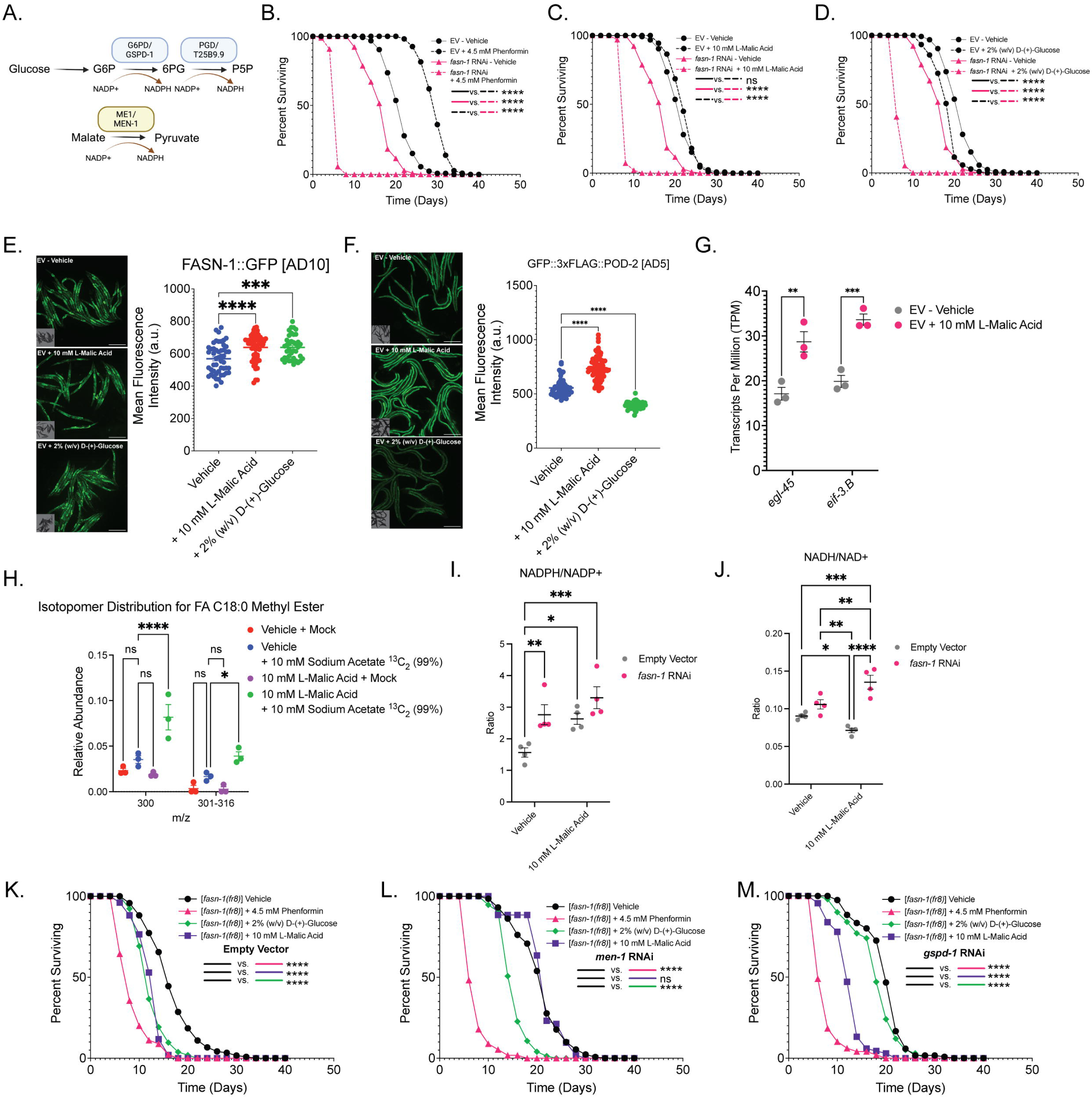
NADPH-generating metabolic interventions require *de novo* fatty acid biosynthesis to buffer reductive stress toxicity. (A) Schematic of metabolic pathways that exogenously increase NADPH production. (B-D) Representative lifespan analyses of WT animals treated with *fasn-1* RNAi or (B) 4.5 mM phenformin, (C) 10 mM L-malic acid (malate), or (D) 2% (w/v) D-(+)-glucose (glucose). Note that these lifespan analyses were performed in tandem and thus contain identical EV - Vehicle curves. (E) Representative micrograph (left) and GFP quantification (right) of FASN-1::GFP animals treated with 10 mM L-Malic Acid or 2% (w/v) D-(+)-Glucose from L4 to AD10. (F) Representative micrograph (left) and GFP quantification (right) of GFP::3xFLAG::POD-2 animals treated with 10 mM L-Malic Acid or 2% (w/v) D-(+)-Glucose from L4 to AD5. Note that the vehicle measurements in these graphs are identical to those shown in 1H and 5K, as these quantifications were performed in parallel. (G) TPM expression values for *egl-45* and *eif-3.B* with *fasn-1* RNAi or 4.5 mM phenformin L4 to AD3. Note that the vehicle measurements in 2D are identical to those in 6G, as these RNA-sequencing experiments were performed in tandem (n = 3 biological replicates). (H) Relative mass count abundance in the 300 and 301-316 m/z mass spectrum of FA C18:0 stearate FAMEs from WT animals treated with 10 mM L-Malic Acid or 10 mM sodium acetate ^13^C_2_ (99%) (n = 3 biological replicates). (I-J) LC-MS analysis of *C. elegans* treated with *fasn-1* RNAi or 10 mM L-Malic Acid with measurement of (I) NADPH/NADP+ or (J) NADH/NAD+. Note that the vehicle measurements in 6I and 6J are identical to those shown in 4B and 4C, respectively, as these quantifications were performed in parallel (n = 4 biological replicates). (K-M) Lifespan analyses of hypomorphic *fasn-1(fr8)* animals treated with vehicle (black), 4.5 mM phenformin (pink), 10 mM L-Malic Acid (purple) or 2% (w/v) D-(+)-Glucose (green) and with the following RNAi: EV control (K), *men-1/CeME1* (L), or *gspd-1/CeG6PD* (M). Lifespans are representative of 3 biological replicates (Table S3). Data represent the mean ± SEM. Imaging data are representative of at least 30 animals aggregated from 3 biologically independent experiments per condition, unless otherwise noted. Scale bar indicates 500 µM. Significance was assessed by (E-F) one-way ANOVA followed by correction for multiple comparisons, (H-J) two-way ANOVA followed by correction for multiple comparisons, (G) DESeq2 Wald tests with Benjamini-Hochberg FDR correction, and (B-D, K-M) log rank analyses. ns p > 0.05, * p < 0.05, ** p < 0.01, *** p < 0.001, **** p < 0.0001.

As we hypothesized, malate supplementation in combination with *fasn-1* RNAi enhanced NADPH/NADP and NADH/NAD ratios and decreased adenylate energy charge, suggesting elevated reductive stress and ATP exhaustion akin to phenformin treatment (Figure 6I-J, S6O). Finally, directly inhibiting the activity of the enzyme predicted to enhance NADPH production downstream of malate (through *men-1* RNAi) or glucose (through *gspd-1* RNAi) suppressed the accelerated death of *fasn-1(fr8)* hypomorphic animals, suggesting that the NADPH-generating activities of malic enzyme and the pentose phosphate pathway, respectively, are required for their early demise (Figure 6K-M). Combined, these data suggest that post-transcriptional rewiring of *de novo* fatty acid biosynthesis acts as a major buffer against NADPH accumulation in diverse metabolic insults to prevent reductive stress and accelerated death.

### Inhibition of human and murine FASN sensitizes cells to reductive stress generating interventions

Observing a synthetic lethal relationship between impaired fatty acid biosynthesis and biguanide treatment in *C. elegans*, we hypothesized a similar role for FASN in mammals to mitigate biguanide-induced reductive stress. Indeed, reanalysis of previously described CRISPR-Cas9 screens revealed that *FASN* and *ACACA* deficiency sensitized human Jurkat T-cells to low-dose phenformin treatment (Figure 7A)^73^. Moreover, metabolomics and functional genomics data from the Cancer Cell Line Encyclopedia suggest that altered accumulation of redox cofactor metabolites associate with increased cellular sensitivity to loss of *FASN* and *ACACA*, consistent with a role for fatty acid biosynthesis in modulating redox homeostasis in humans (Figure 7B-C)^74^. In aggregate, these data suggest the possibility that *de novo* fatty acid biosynthesis may play a conserved role to buffer biguanide-induced reductive stress.

**Figure 7.**
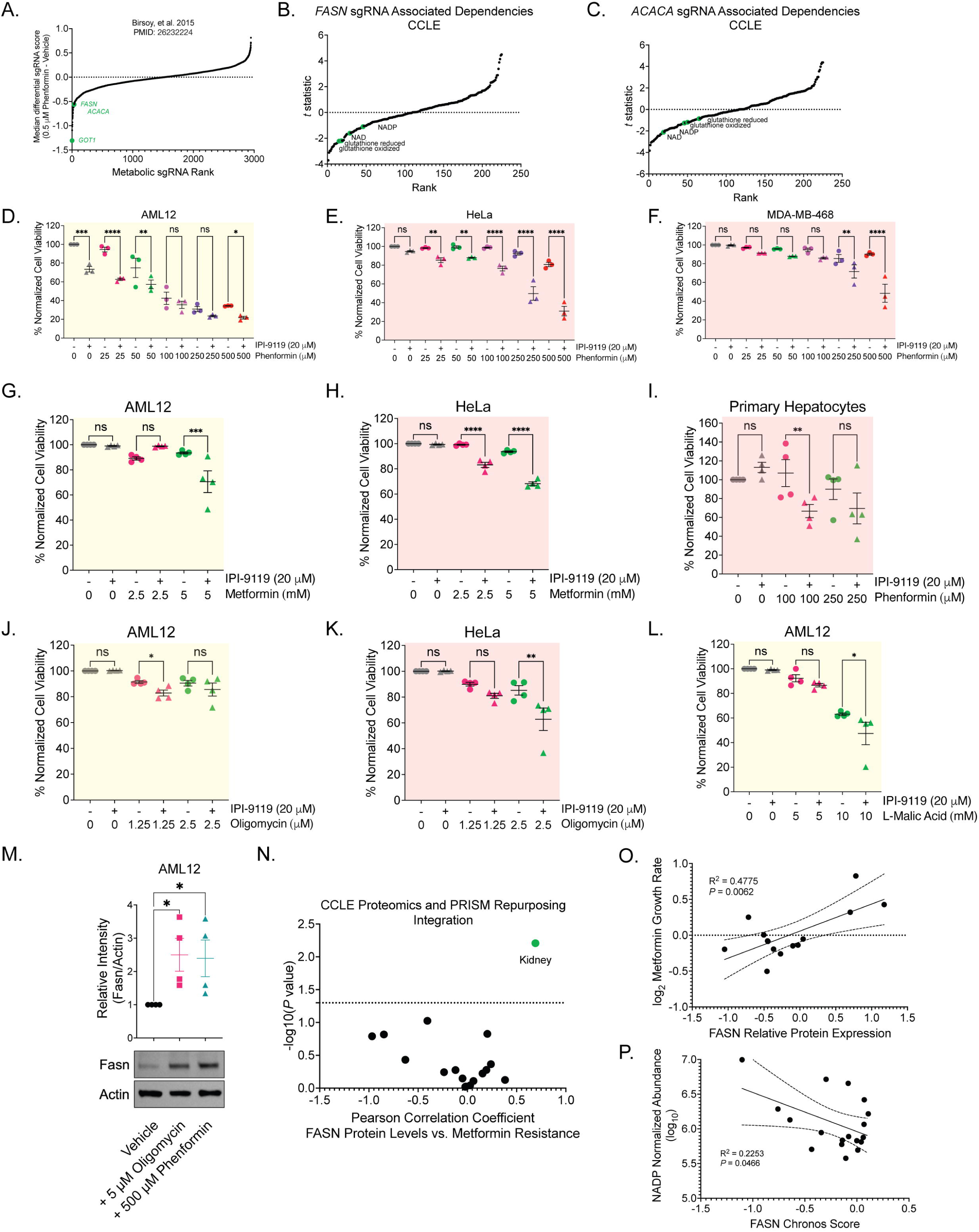
FASN inhibition in murine and human cellular models synergistically enhances sensitivity to biguanides. (A) Metabolic sgRNA rank data from Birsoy, et al.^73^ (PMID: 26232224) indicating that FASN and ACACA deficiency are sensitizers to low dose phenformin treatment in Jurkat T-cells. (B-C) *t*-statistic analysis of Cancer Cell Line Encyclopedia (CCLE) human cell lines associating metabolite variance with deficiencies in *FASN* (B) or *ACACA* (C) from DepMap CRISPR-Cas9 pooled screens. Associations were extracted from the CCLE metabolomics project^74^. Redox associated metabolites are highlighted in green. (D-F) Crystal violet staining (CVS) intensity of murine hepatocyte AML12 (D), human cervical carcinoma HeLa (E), and human breast adenocarcinoma MDA-MB-468 (F) cells treated for 48 hours prior to analysis with the indicated doses of IPI-9119 and phenformin (n = 3 biological replicates). (G-I) CVS intensity of AML12 (G), HeLa (H), and isolated primary human liver hepatocytes (I) treated for 48 hours with the indicated doses of IPI-9119 and metformin (G-H) or phenformin (I) (n = 4 biological replicates). (J-L) CVS intensity of AML12 (J,L) and HeLa (K) cells treated for 48 hours with the indicated doses of IPI-9119 and oligomycin (J-K) or L-Malic Acid (L) (n = 4 biological replicates). (M) Representative western blot and quantification for FASN in AML12 cells treated with oligomycin or phenformin for 48 hours (n = 4 biological replicates). (N) Pearson correlation coefficients of lines from the CCLE^110^, comparing FASN protein levels with relative resistance to metformin-mediated growth inhibition (Drug Repurposing Study) and stratified by cellular tissue lineage. Data were extracted and analyzed using Data Explorer v2.0 on the DepMap^76^. Kidney lineage is highlighted in green. (O) Pearson correlation coefficients of kidney lineage cancer cell lines from the analysis above. (P) Pearson correlation coefficients of kidney lineage cancer cell lines^74^, comparing NADP metabolite abundance with their respective growth sensitivity to FASN deficiency via CRISPR-Cas9 Chronos Score (CCLE)^110^. Cells were passaged/thawed and plated for 2 days prior to drug treatment for cells to reach at least 70% of confluency. Data represent the mean ± SEM. All non-IPI-9119 treated cells were treated with an equivalent percentage (v/v) of DMSO to control for off-target toxicity. Significance was assessed by (D-L) two-way ANOVA followed by correction for multiple comparisons, (M) one-way ANOVA, or (B-C, N-P) extracted from DepMap Data Explorer v2.0. ns p > 0.05, * p < 0.05, ** p < 0.01, *** p < 0.001, **** p < 0.0001.

To formally test if fatty acid biosynthesis buffering biguanide-induced reductive stress is a conserved requirement, we leveraged a potent, irreversible inhibitor of human and murine fatty acid synthase, IPI-9119, in combination with phenformin treatment in a panel of cancer and immortalized cells lines^75^. We observed that combinatorial IPI-9119 and phenformin administration in murine AML12 (hepatocyte), human HeLa (cervical adenocarcinoma) and human MDA-MB-468 (breast adenocarcinoma) cells synergistically reduced cellular viability in a dose dependent manner (Figure 7D-F, S7A-C). We additionally observed a similar synthetic lethality relationship between FASN inhibition and metformin treatment, consistent with a conserved role for FASN in buffering biguanide-mediated toxicity (Figure 7G-H). Compellingly, this synthetic lethal interaction extends to primary human hepatocytes, suggesting that fatty acid biosynthesis may prevent systemic, medically relevant toxicity to biguanide exposure (Figure 7I). Consistent with our hypothesis that FASN buffers against biguanide-induced reductive stress and ATP exhaustion, we observed a similar synthetic lethal relationship between FASN inhibition and oligomycin or malate treatment (Figure 7J-L). In line with the translational protection of *fasn-1* and *pod-2* in *C. elegans*, we noted that phenformin treatment increases FASN and ACC levels in HeLa, AML12, and MDA-MB468 cells, and increased FASN levels in AML12 cells treated with oligomycin, indicating a conserved, post-transcriptional response to reductive stress generating insults (Figure 7M, S7D).

Lastly, we hypothesized that combinatorial FASN inhibition and biguanide treatment may represent an untapped therapeutic strategy to mitigate growth of reductive stress-sensitive cancers. We noted a significant association between FASN protein expression and resistance to metformin-mediated growth inhibition, specifically in kidney cancer subtypes from the CCLE and DepMap (Figure 7N-O, S7E), as well as strong association between enhanced accumulation of redox associated metabolites NADP and GSH with sensitivity to FASN inhibition (Figure 7P, S7F)^76^. We finally noted that kidney renal cell and papillary cell carcinomas display elevated levels of FASN relative to control, and that elevated FASN levels are associated with poorer prognoses (Figure S7G-H). This suggested that reducing *de novo* fatty acid biosynthesis in tandem with biguanide treatment may create a new metabolic vulnerability in these redox-sensitive cancers. Combined, these results suggest that enhanced *de novo* fatty acid biosynthesis is an ancient, conserved mechanism that protects metazoans from the deleterious effects of biguanides and other reductive stress-associated interventions.

## DISCUSSION

Here, we identified an ancient, conserved, post-transcriptional mechanism by which metazoans alleviate reductive stress via activation of *de novo* fatty acid biosynthesis. While phenformin reduces global mRNA translation in nematodes through attenuation of polysome engagement, key fatty acid biosynthetic enzymes FASN-1 and POD-2 are translationally protected to prevent biguanide-mediated synthetic lethality. We find that translational protection of *pod-2* is partly mediated by eIF3 complex members EGL-45/CeEIF3A and EIF-3.B/CeEIF3B, loss of which mirrors BIG-SAD with *fasn-1* and *pod-2* depletion. Importantly, BIG-SAD is not due to reduced biogenesis of a fatty acid product, but instead through failure to consume excess NADPH, resulting in electron spillover into NADH and GSH pools and subsequent reductive stress toxicity. We show that reductive stress exerts its toxicity by hyperactivating SKN-1/CeNRF and dampening AAK-2/CeAMPK function, resulting in a reversal of biguanide mediated pro-longevity outcomes and accelerated ATP exhaustion. Importantly, this protective mechanism is required by diverse NADPH-generating metabolic interventions, as *de novo* fatty acid biosynthesis is broadly required to buffer toxicity resulting from unchecked reductive stress. Finally, this requirement for fatty acid biosynthesis is conserved to mammals and even humans, as sublethal FASN inhibition sensitizes primary human hepatocytes and a variety of immortalized and cancerous cell lines to the effects of biguanides and associated reductive stress.

### mRNA translation fine tunes biguanide-induced toxicity responses

Multiple mechanisms have been proposed to underly metformin’s geroprotective effects, including activation of AMPK through lysosomal v-ATPases, altered gut microbiome metabolism, enhanced mTORC1 inhibition, and altered ether lipid production^8,9,11–15^. Here, we suggest cytosolic translation inhibition as an additional hallmark of biguanide action in aging. This link between biguanide action and translation inhibition builds upon a burgeoning literature in intraorganellar communication between the mitochondrion and cytoplasm, raising a question as to what signals mediate this crosstalk^35,38,42,46,67^. Previous studies suggest that ETC inhibition activates the integrated stress response (ISR), which results in dampened cytosolic mRNA translation concomitant with activation of a cytoprotective transcriptional program^67,77,78^. Activation of the ISR transcription factor ATF4/Gcn4 extends organismal lifespan, and ATF-4 is required for the pro-longevity effects of mRNA translation and mTORC1 inhibition paradigms in *C. elegans* ^40,79^. Indeed, the loss of polysome occupancy coupled with post-transcriptional protection of key regulators seen in our study mirrors ISR activation; however, we find that biguanide-mediated lifespan extension in nematodes does not require activation of the ISR or ATF-4, and biguanide treatment enhances the TE of transcripts not protected during canonical ISR activation (data not shown). It remains of interest to determine if biguanides induce a mitochondrial metabolite or signaling-related factor to elicit a global decrease in cytosolic polysomes while protecting translation of a select few transcripts.

Despite a global decrease in polysome occupancy, our studies have identified 26 genes with enhanced translational efficiency upon phenformin treatment. Sequence specificity has been shown to critically influence the rate of mRNA translation, through targeting of the 5’ untranslated region (UTR) and 5’ terminal CDS of mRNA transcripts^80^. Several sequence motifs are involved in enhancement of translation efficiency, including the presence of Kozak sequences and upstream open reading frames^81^. Additionally, *trans*-splicing of spliced leader 1 (SL1) sequences has been shown to enhance translational efficiency of essential *C. elegans* genes, mechanistically hypothesized through shortening of native 5’ UTRs and improved RNA binding protein (RBP) binding potential^82^. Translational regulation through the 3’ UTR has been described in neuronal dendrites, in part through cooperatively enhancing RBP recruitment and subsequent translation initiation machinery^83^. We note that *fasn-1* and *pod-2* both contain SL1 target sites, and analyses from modENCODE datasets have revealed that *fasn-1* and *pod-2* mature mRNA transcripts predominantly contain SL1 sites ^84^. However, approximately 62% of the *C. elegans* transcriptome is SL1 *trans*-spliced, suggesting that *trans*-splicing alone is not sufficient to explain why only 26 genes display enhanced translation efficiency with phenformin treatment^82,84,85^. A broader interrogation of RBPs and *trans*-splicing factors that may cooperatively bind to these 26 transcripts is required to understand how these transcripts are protected during biguanide exposure.

We identified EGL-45/CeEIF3A and EIF-3.B/CeEIF3B as factors that directly mediate the translational protection of *pod-2*. The eIF3 complex acts as the major assembly scaffold for interactions between ribosomal subunits, target mRNA species, RBPs, and other eIFs. Intriguingly, recent studies have shown that select eIF3 subunits can differentially alter translation efficiency of specific transcripts^52,86–90^. Moreover, inhibition of specific eIF3 subunits have been shown to extend *C. elegans* lifespan through mechanisms distinct from canonical pro-longevity paradigms, suggesting that individual eIF3 subunits may have diverse roles in stress resolution^86^. *C. elegans* modENCODE transcription factor (TF) binding site analysis of the promoters of *egl-45* and *eif-3.B* suggest that a number of TFs may coordinate enhanced transcription of these factors with biguanide treatment^91^. It will be of great interest to both validate the concert of TFs that mediate enhanced *egl-45* and *eif-3.B* transcription, and define how these TFs are activated by biguanide-specific signaling mechanisms. Additional work will be required to determine how EGL-45 and EIF-3.B selectively recognize *pod-2* transcripts, which we speculate may be through specific interactions with the 5’ or 3’ UTRs of the transcript. Lastly, we note that post-developmental inactivation of *egl-45* and *eif-3.B* has previously been shown to robustly extend organismal lifespan, an observation that we corroborate in this study^33^. How knockdown of these genes can display differential longevity responses upon biguanide treatment will be important to explore, as observations we present imply that the longevity promoting signals that are beneficial during eIF3 subunit knockdown must be diverted to compensate for biguanide exposure.

### De novo fatty acid biosynthesis detoxifies NADPH-driven reductive stress

We demonstrate that biguanides enhance NADPH/NADP+, NADH/NAD+, and GSH/GSSG redox ratios, and that the increased accumulation of NADPH during inactivation of *fasn-1* leads to redox spillover, resulting in global elevation of reductive stress driven toxicity and organismal demise. Our study unexpectedly finds that NADPH accumulation drives biguanide-induced reductive stress, suggesting that previously established reductive stress outcomes of biguanide exposure rely principally upon NADPH, but not NADH, homeostasis to prevent catastrophic outcomes^31,92–94^. Reductive stress has been relatively understudied as a redox imbalance disorder, leading to disrupted pathophysiology, cardiomyopathies, cancers, and alcohol-associated liver disease^29–31,93,95–97^. Our data suggest that fatty acid biosynthesis buffers against NADPH-induced reductive stress during biguanide exposure, adding to a number of mechanisms that act as redox cofactor sinks, including inhibition of purine salvage during NADH reductive stress, and proteasomal degradation of FNIP1 to reverse toxicity due to low ROS from persistent antioxidant signalling^63,98–100^. The generality of this mechanism is revealed during malate or glucose treatment, in which fatty acid synthase acts as a sink for NADPH reductive stress. Thus, our findings suggest that multiple NADPH producing paradigms absolutely require enhanced fatty acid synthase activity to consume potentially toxic reducing equivalents^57^. We also demonstrate that the fatty acid synthase-mediated consumption of NADPH yielded during malate incorporation is strictly required to establish the pro-longevity benefits of malate supplementation in *C. elegans*^101^. We hypothesize that there may exist analogous consumption mechanisms to alleviate NADH or GSH reductive stress, respectively, in specific contexts during aging.

How elevating NADPH reducing equivalents “spill over” into NADH and GSH reducing pools is poorly understood. Previous studies indicate that these pools are spatially compartmentalized, with the mitochondrial inner membrane preventing diffusion of mitochondrial NAD(P)H into the cytoplasm^29,102^. However, exchange of cytoplasmic and mitochondrial NAD(P)H pools are possible through the activity of isocitrate-KG and malate-aspartate shuttles^29,103,104^. Cytoplasmic GSH is generated by glutathione reductase activity using NADPH as an electron donor, suggesting that enzymatic activity can also enable redox pool shuttling^29^. It is possible that biguanide treatment may redirect excess reducing equivalents into cytoplasmic NADPH, with fatty acid biosynthesis ultimately consuming the excess NADPH to ensure redox homeostasis. It is also possible that this inability to prevent spillover results in ATP exhaustion and aberrant antioxidant gene expression programs, as suggested by the impaired AAK-2/CeAMPK and hyperactivated SKN-1/CeNRF activities established in this study. It will be of interest to identify the shuttling mechanism(s) that directly exchange reducing equivalents between the NADH/NAD+, GSH/GSSG, and NADPH/NADP+ redox couples, and in which organelles these effects are primarily executed. Our experiments utilizing TPNOX suggest that the cytoplasmic consumption of NADPH, akin to the localization of fatty acid synthase, can significantly mitigate reductive-stress associated toxicity, but compartment specific visualization of redox status in metazoan organelles and tissues are needed to confirm these findings.

### Dialing down the stress: manipulating metabolic pathways to rebalance redox homeostasis

Recent studies have shown that cleavage of FASN-1 by CED-3 caspase is required to signal pan-stress resolution^105,106^. We propose that in addition to this, the conserved NADPH consuming properties of *de novo* fatty acid biosynthesis limit aberrant stress responses to biguanide treatment. Thus, *de novo* fatty acid biosynthesis may act as a tunable rheostat to buffer reducing equivalents and limit consequent stress response programs upon redox-altering insults. Given the evidence for these metabolic pathways to be regulated post-transcriptionally, the potential to manipulate *de novo* fatty acid biosynthesis via altered translation factor activity and polysomal engagement presents as an attractive node to maximize the potential geroprotective properties of biguanides.

### Manipulating redox homeostasis to target metabolic vulnerabilities in neoplasia

Recent studies have intricately defined the differential capacity for cancer subtypes to buffer reductive stress, suggesting that heretofore underappreciated or unknown genetic and pharmacologic modalities may be combinatorically repurposed to exploit vulnerabilities in selective carcinomas^31,92,93^. We propose that in addition to established NADH and GSH driving interventions, inducing NADPH reductive stress may serve as a metabolic vulnerability in neoplasia, motivating systematic dissection of the cancer subtypes that may benefit from this therapeutic approach.

### Limitations of the study

While our current study identified critical translation factors that regulate *pod-2* translation efficiency during biguanide treatment, we did not identify a factor that mediates translational protection of *fasn-1*. Additional genetic screening strategies to identify such factors will be critical to determine how FASN-1/CeFASN protein levels are modulated during reductive stress-inducing interventions. We additionally did not identify which sequence(s) in *fasn-1* or *pod-2* transcripts are required for polysome engagement, which will require future ribosome footprinting and enhanced cross-linked RNA immunoprecipitation^107,108^ approaches.

## Supporting information

Data S1 - Source Data and Statistics

Data S2 - Unprocessed Immunoblot Images

## RESOURCE AVAILABILITY

### Lead contact

Further information and requests for resources and reagents should be directed to and will be fulfilled by the Lead Contact, Alexander A. Soukas

### Materials availability

Plasmids, oligonucleotides, worm strains, or cells utilized in this study are available through the Lead Contact upon request.

### Data and code availability

- Raw values for the metformin genome-wide RNAi screen are included in this manuscript as **Table S1**. All RNA-sequencing raw data has been deposited to the NCBI Gene Expression Omnibus (NCBI GEO) and is available under accession number GSE277946. RNA-sequencing processed data are provided in this manuscript as **Table S2** (polysome profiling) and **Table S4** (BIG-SAD). All lifespan statistics presented in this manuscript have been included as **Table S3**. Metabolomics and lipidomics raw data have been deposited to Mendeley and can be accessed at **DOI:10.17632/28gvnkd8r4.1**. Metabolomics and lipidomics processed data are provided in this manuscript as **Table S5**. All other raw quantification values for graphs and unprocessed immunoblotting images in this manuscript have been provided as **Data S1**. All other original microscopy images will be shared by the Lead Contact upon request.
- This study does not report original code.
- Any additional information required to reanalyze the data reported in this manuscript will be made available by the Lead Contact upon request.

## ACKNOWLEDGEMENTS

We thank members of the Soukas Laboratory for their helpful feedback and discussions related to this manuscript. We thank T. Keith Blackwell (Joslin Diabetes Center), Sean P. Curran (University of Southern California), Martin S. Denzel (Altos Labs), Dennis H. Kim (Boston Children’s Hospital), William B. Mair (Harvard T.H. Chan School of Public Health), Kunihiro Matsumoto (Nagoya University), and Read Pukkila-Worley (University of Massachusetts Chan Medical School) for providing strains and additional reagents. We thank Richard Bouley and the Program in Molecular Biology (PMB) Microscopy Core at Mass General Brigham for confocal imaging assistance. Some strains were provided by the CGC, which is funded by the NIH Office of Research Infrastructure Programs (P40 OD010440). This work was funded by the NIH (R01 AG058259 and R01 AG69677 to A.A.S.), the Nutrition Obesity Research Center at Harvard (NORCH) (P30 DK040561 to A.A.S.), the NIH/NIDDK-funded Boston-Area DERC (P30 DK057521 to A.A.S.), the NIH/NIDDK T32 Harvard Training Program in Bioinformatics Applied to Diabetes, Obesity, and Metabolism (5T32DK110919-02 to D.P.), and the Weissman Family MGH Research Scholar Award (to A.A.S.). F.M.A. was supported by the PhD Program in Biological and Biomedical Sciences at Harvard Medical School, Division of Medical Sciences and the Albert J. Ryan Foundation Fellowship. Visualizations in this manuscript were generated by BioRender.com.

## AUTHOR CONTRIBUTIONS

Conceptualization, F.M.A and A.A.S.; Methodology, F.M.A, J.F.R., A.I.Y., S.W.E., N.L.S., J.A.A-S., W.W., D.J.B., D.P., O.S.S., M.D.B. and A.A.S.; Formal Analysis, F.M.A, J.F.R., A.I.Y., S.W.E., N.L.S., J.A.A-S., W.W., D.J.B., D.P., and O.S.S.; Investigation, F.M.A, J.F.R., A.I.Y., S.W.E., N.L.S., J.A.A-S., W.W., D.J.B., D.P., and O.S.S.; Resources, W.W., O.S.S., M.D.B., and A.A.S.; Writing – Original Draft, F.M.A. and A.A.S.; Writing – Reviewing & Editing, F.M.A. and A.A.S.; Visualization, F.M.A., J.F.R., D.P., O.S.S. and M.D.B.; Supervision, A.A.S.; Funding Acquisition, A.A.S.

## KEY RESOURCES TABLE

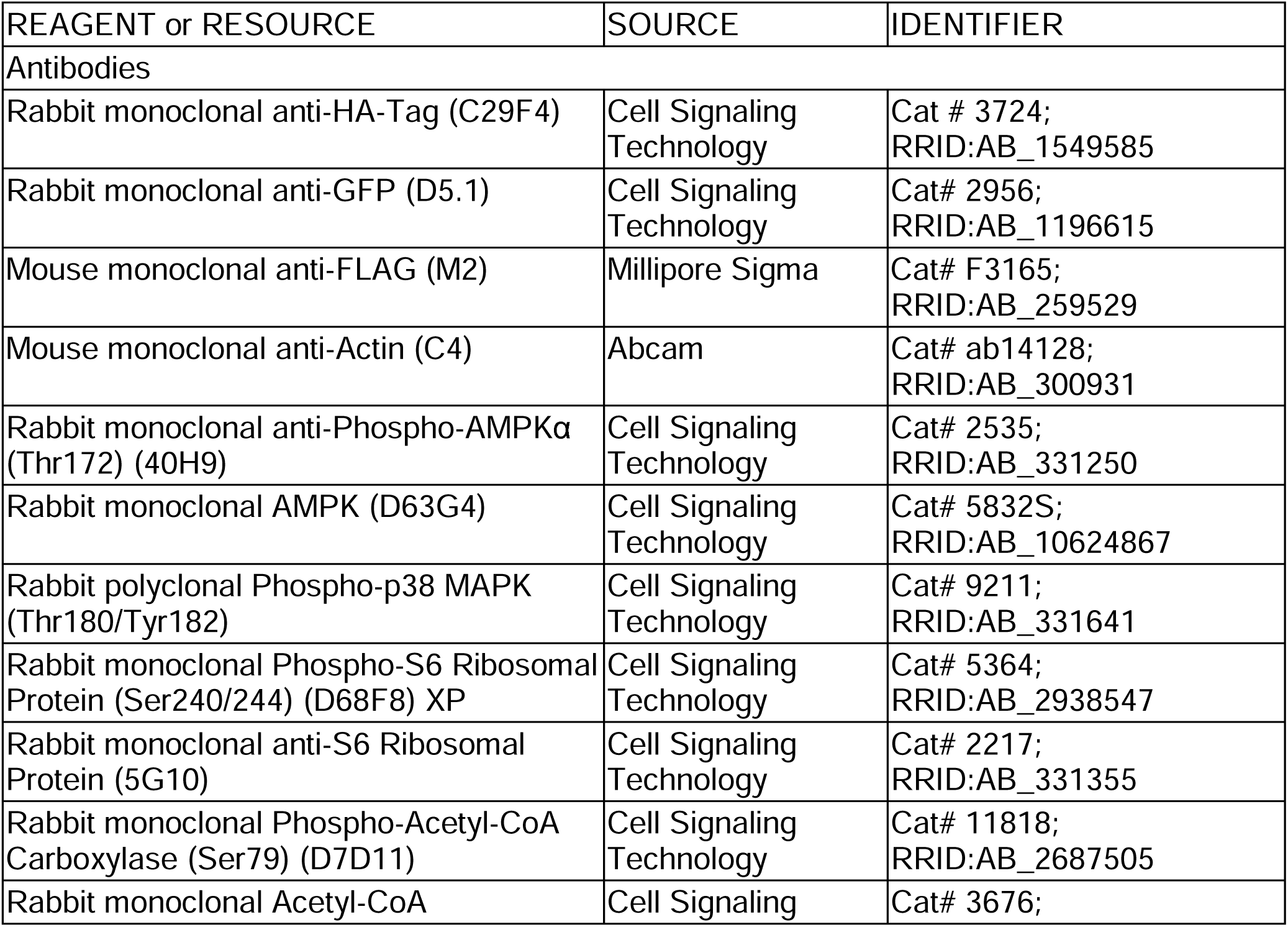
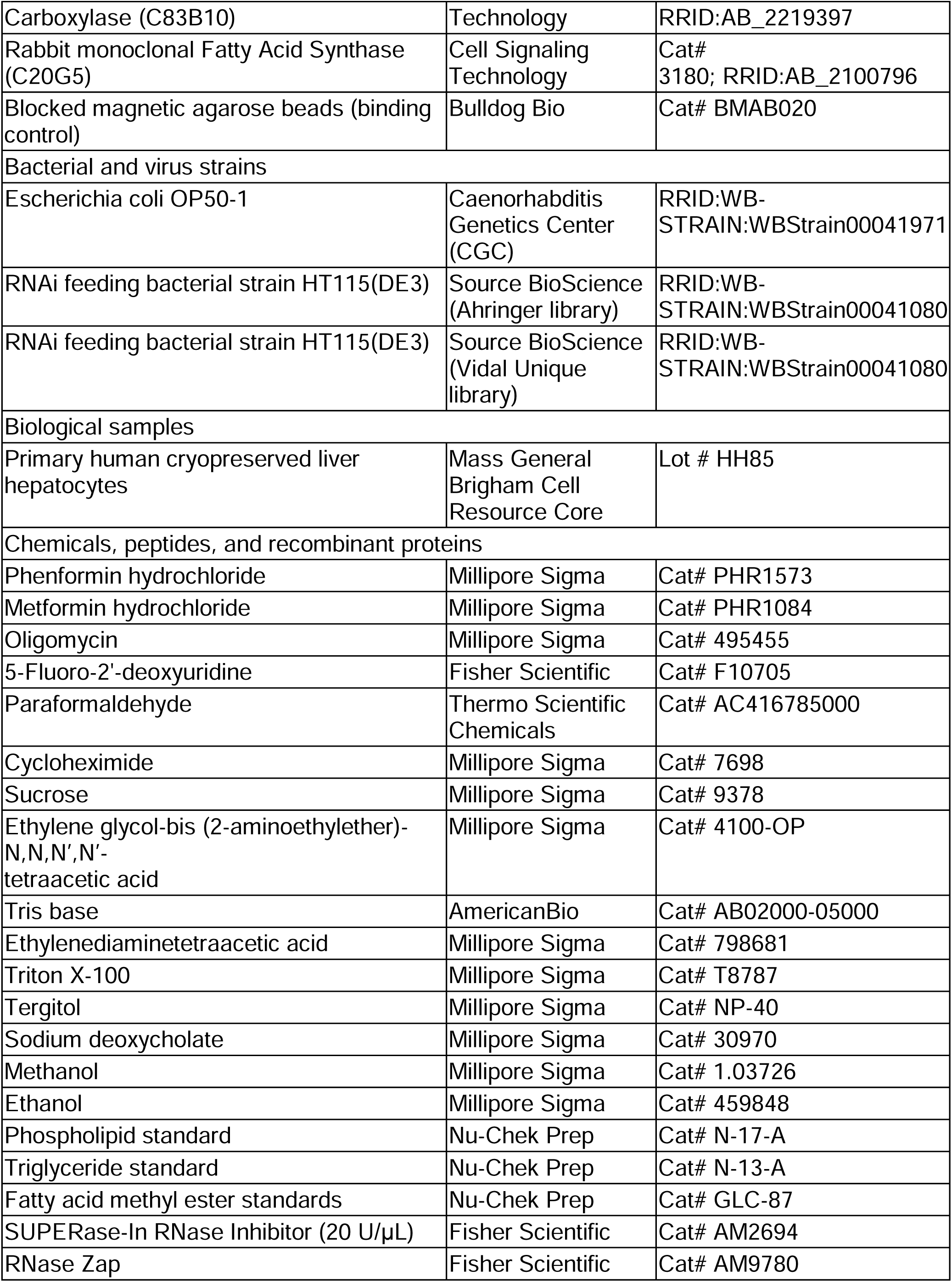
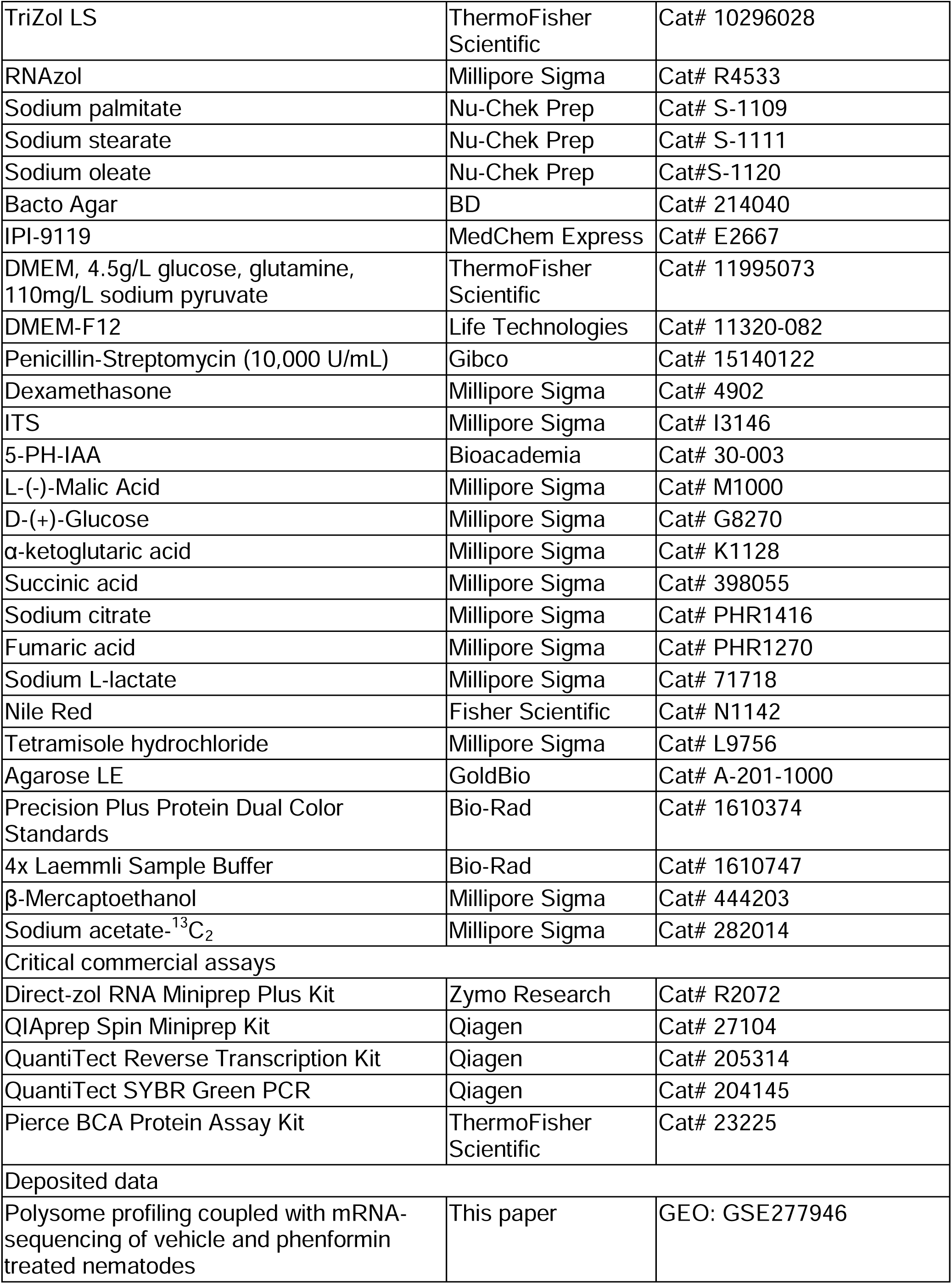
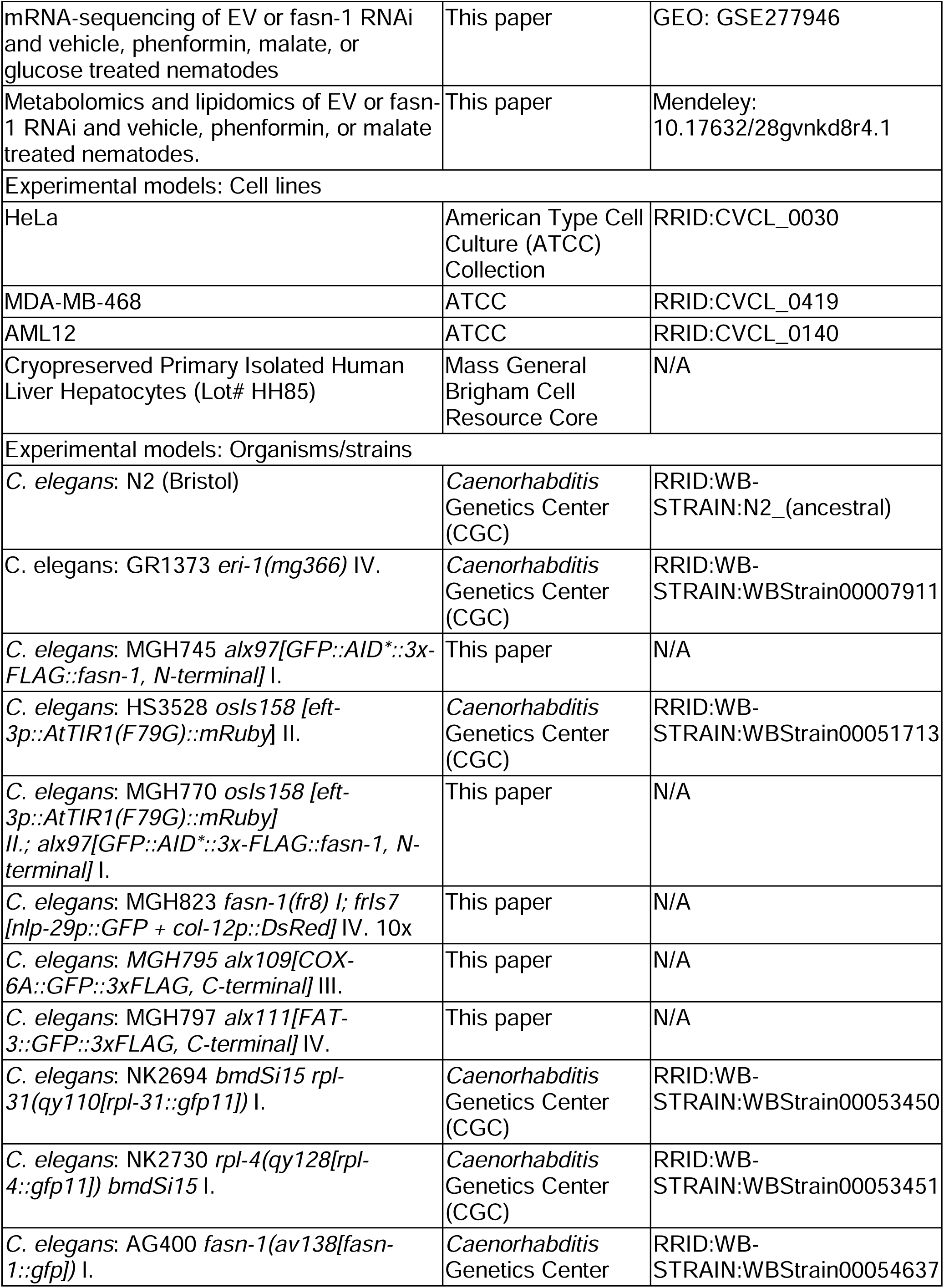
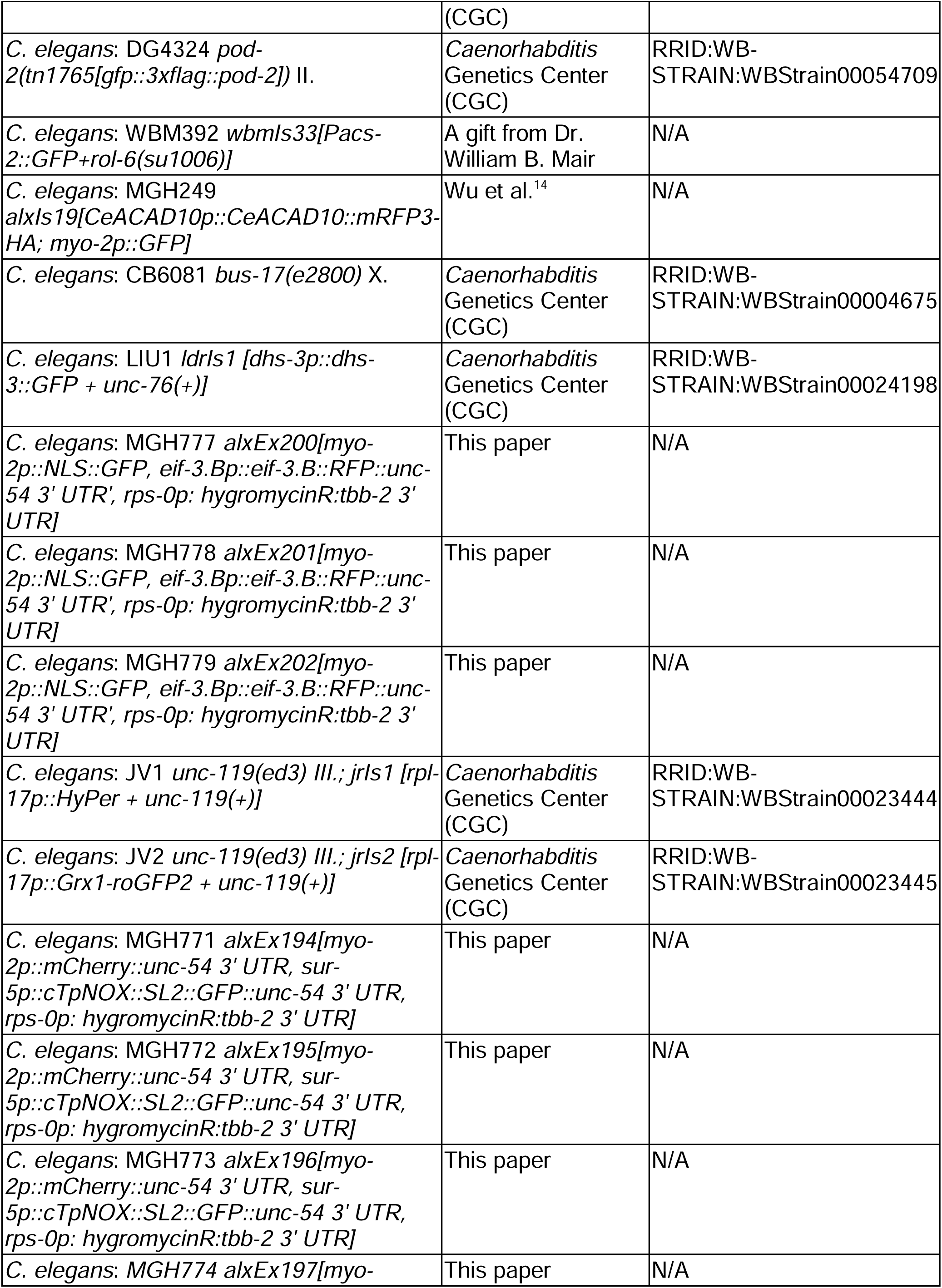
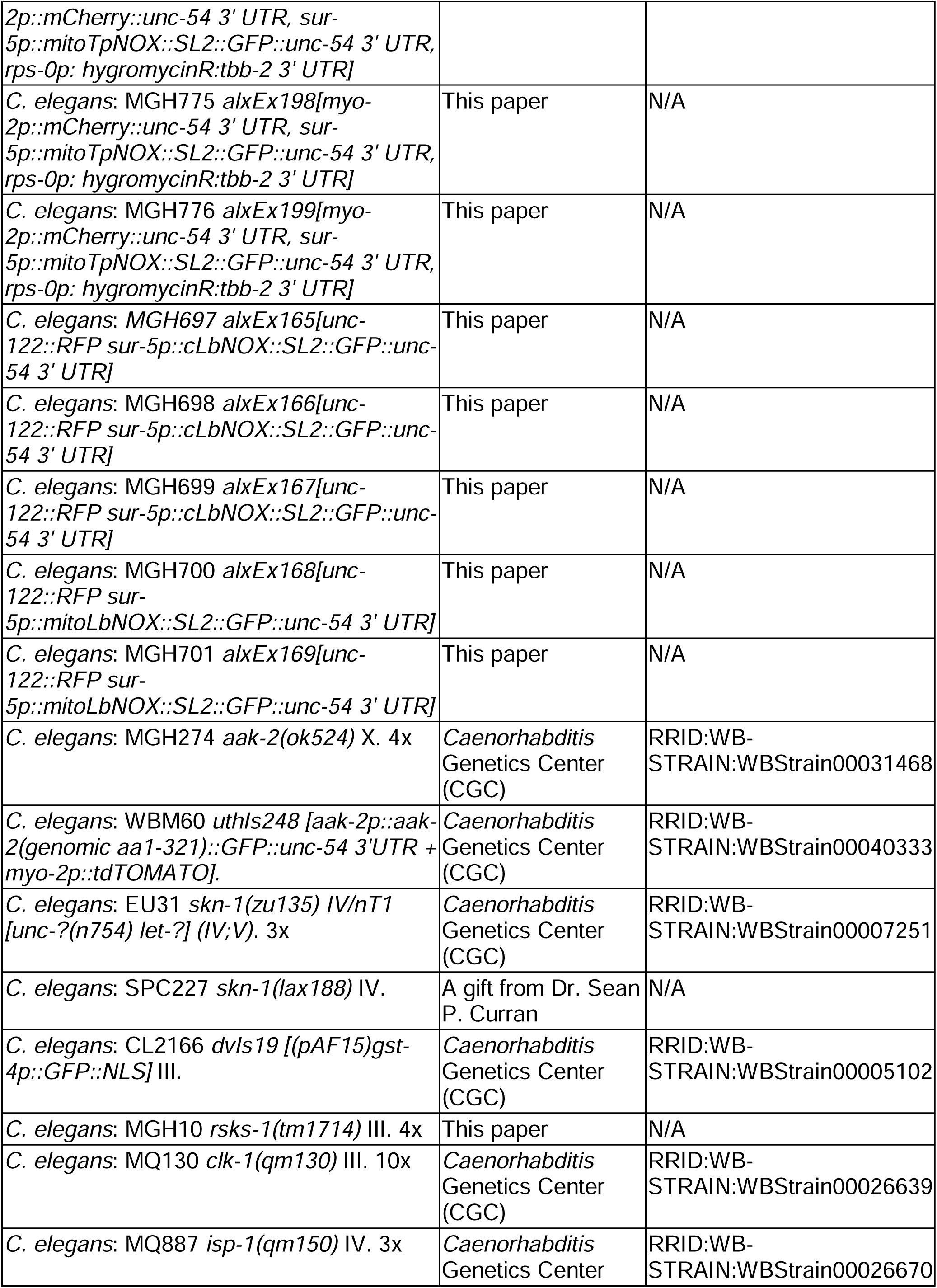
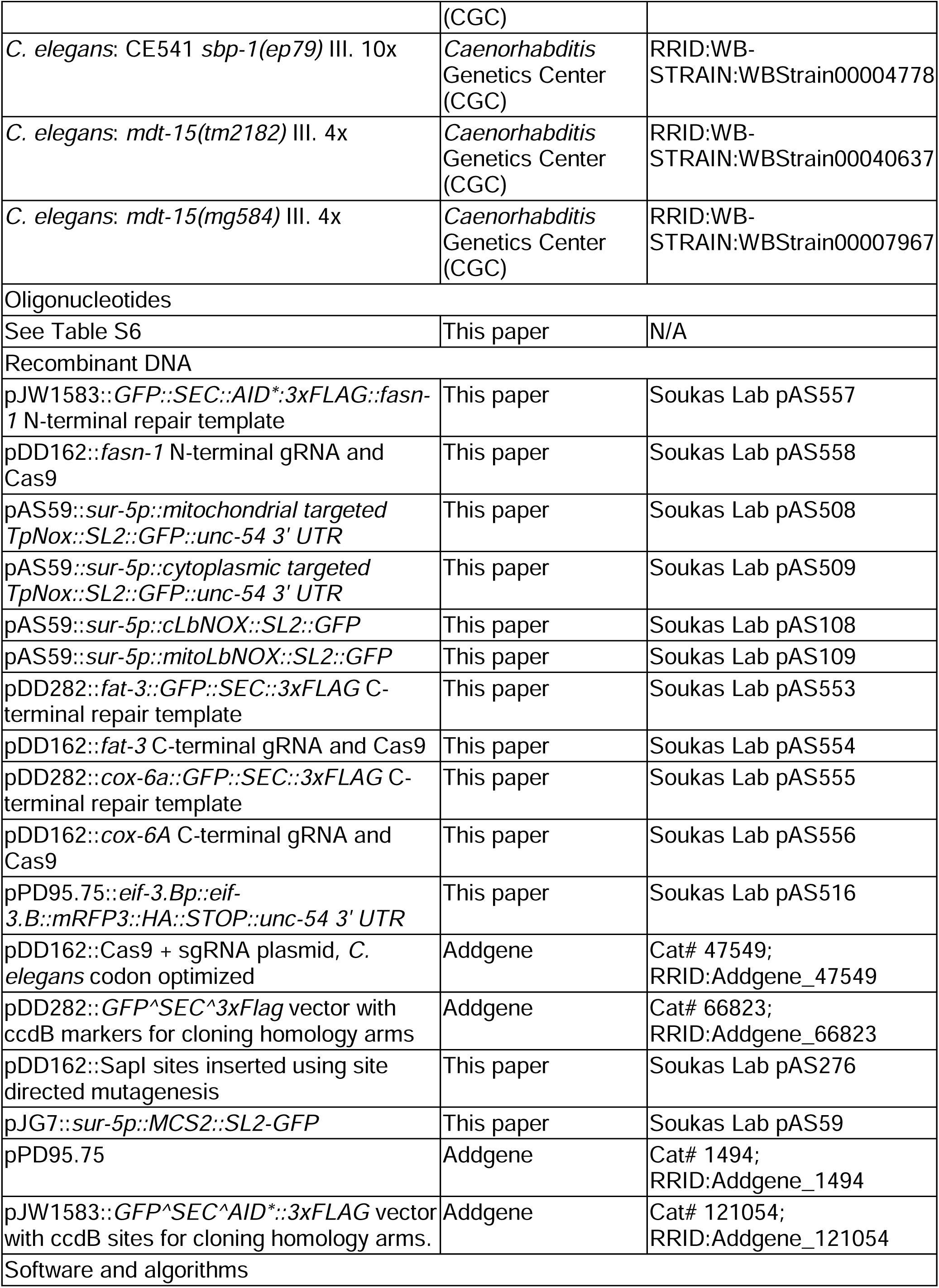
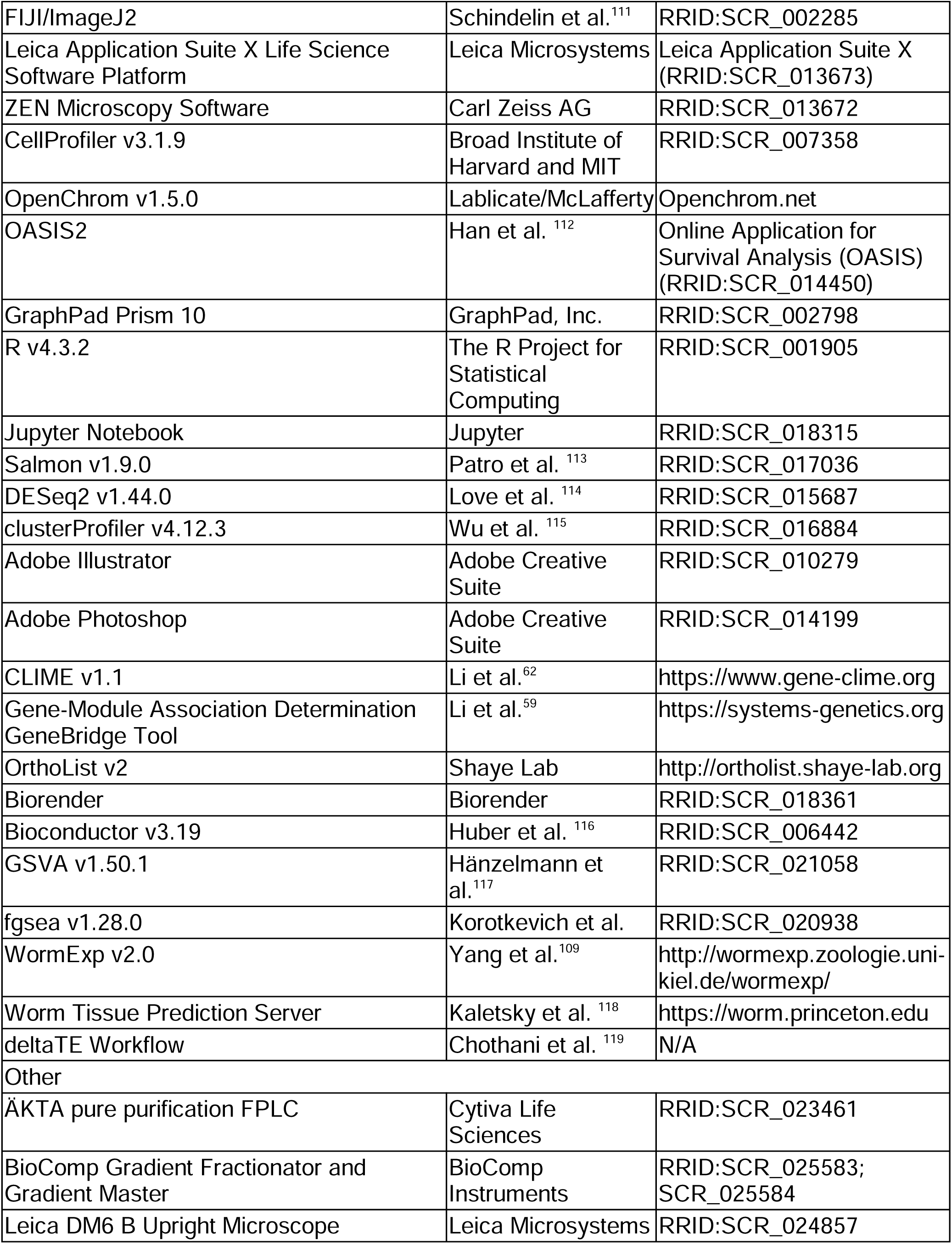

## STAR METHODS

### EXPERIMENTAL MODEL AND STUDY PARTICIPANT DETAILS

#### C. elegans nematode strains and maintenance

All nematode strains used in this study are included in the Key Resources Table. All strains were genotype/transgene expression validated and backcrossed to ancestral N2 Bristol [RRID: WB-STRAIN:N2_(ancestral)] wild-type nematodes at least 3 times (unless otherwise noted) prior to use in these studies. Strains were maintained on nematode growth medium (NGM) agar plates seeded with *E. coli* OP50-1 and maintained non-starved for at least 2 generations prior to start of studies^120^. Nematode strains were grown at 20°C in a temperature and humidity controlled Thermo Scientific Forma Environmental Chamber incubator, unless otherwise noted. Ancestral strains used in this study were freshly thawed every 3 months to prevent genetic drift.

Animals were synchronized in these studies using alkaline hypochlorite bleach treatment from gravid adults. Animals were first collected in M9 medium, centrifuged at 3300xg for 1 minute, and then resuspended in 6 mL of a 1.3% household bleach, 250 mM NaOH supplemented M9 solution with 1 minute of vigorous shaking by hand. Bleach solution exposure and shaking were repeated once more following an M9 wash to ensure cuticle degradation and egg release. After the second bleaching, eggs were resuspended gently in M9 media and washed four times to remove bleach solution. The eggs were then resuspended in 10 mL M9 solution and rotated overnight for 18h at 20°C. Hatched and arrested L1 larva were washed and resuspended in 10 mL M9 to remove excess dauer hormone, counted, and dispensed onto NGM or RNAi plates, as indicated below.

#### E. coli bacterial strains and maintenance

All non-RNAi experiments were conducted on NGM Bacto agar or NGM agarose LE (for use in metformin-related experiments) plates supplemented with *E. coli* OP50-1 as the food source^121^. Cultures of *E. coli* OP50-1 were grown in LB broth supplemented with streptomycin for 48h at 37°C without shaking and seeded directly onto NGM plates. Seeded cultures were dried under ambient conditions for 2 days prior to use, and plates were used within 3-7 days after seeding for nematode strain maintenance or experimental work.

#### Mammalian tissue culture cell lines and culture conditions

All tissue and cell culture lines used in these studies are listed in the Key Resources Table. All lines were routinely tested for mycoplasma contamination via PCR prior to experimentation. AML12 hepatocytes were grown in a 1:1 mixture of Dulbecco’s Modified Eagle’s Medium (DMEM) and Ham’s F12 medium (DMEM-F12, Thermo Fisher) supplemented with 0.005 mg/mL insulin, 0.005 mg/mL transferrin, 5 ng/mL selenium, 40 ng/mL dexamethasone, 10% (v/v) fetal bovine serum (FBS) (GeminiBio), and 1% (v/v) penicillin-streptomycin (Life Technologies). Primary cryopreserved human liver hepatocytes were obtained from the Mass General Brigham Cell Resource Core and grown in C+H media obtained from the Cell Resource Core. Primary hepatocytes were maintained for no more than 4 days for the duration of experimentation. All other cell lines were grown in DMEM supplemented with 4500 mg/L D-glucose, 584 mg/L L-glutamine, 110 mg/L sodium pyruvate, 15 mg/L phenol red (Gibco), 10% (v/v) FBS, and 1% (v/v) penicillin-streptomycin. All cells were grown in a 37°C tissue culture incubator in a 5% CO_2_ water humidified environment.

### METHOD DETAILS

#### Genome-wide RNAi screening for metformin response regulators

96-well high throughput microscopic assessment of body growth size following 72 hours of 150 mM metformin treatment was performed as previously described^53,122^. Briefly, *eri-1(mg366)* animals were bleach synchronized for 18 hours, and L1s were subsequently dropped onto 96-well RNAi agarose plates pretreated with 150 mM metformin and seeded with 18,179 individual RNAi clones (corresponding to 86% of all *C. elegans* CDS) and dried under laminar flow. Animals were then incubated for 72 hours at 20°C prior to microscopic analysis. Experiments were performed in biological duplicate. All images were then extracted, animals identified using a custom-built machine learning object identification algorithm using CellProfiler v3.1.9, and individual worm lengths in pixels were calculated. Average ΔZ (calculated as the 150 mM metformin worm length – vehicle control) were calculated for each replicate and well, and missing values were imputed using the Multivariate Imputation by Chained Equations (MICE) strategy. RNAi clones with an average ΔZ score greater than - 0.395 were selected for downstream Gene Ontology and Gene Set Enrichment analyses. Gene set overrepresentation analyses were performed using R package clusterProfiler v4.12.3 for GO term-based pathway annotations^115^, and Gene Set Enrichment Analyses was performed using R package fgsea v1.28.0 and plotted using R package ggplot2 v3.5.0.

#### Polysome profiling and density ultracentrifugation coupled with bulk RNA-sequencing

Polysome profiling using *C. elegans* lysates was performed as previously described with minor modifications^123^. For each sample, 30,000 animals of the indicated stage, genotype, strain, or treatment were harvested from 10 cm NGM or RNAi agar plates in M9 and washed three times via gravity settling with fresh M9 to remove bacteria. Worm slurries were then concentrated and solubilized in worm solubilization buffer containing 50 mM Tris-HCl (pH 8.0), 300 mM NaCl, 10 mM MgCl2, 1 mM EGTA, 0.1 mg/ml cycloheximide, 1% Triton X-100, 0.1% sodium deoxycholate,1x cOmplete EDTA-free protease inhibitor, and 125 U/µl SUPERase•In RNAse inhibitor. Worms were then homogenized using a motorized pellet pestle with 120 strokes on ice. Homogenates were solubilized at 4°C on a tube rotator for 1 hour and then cleared via centrifugation at 14,000 x g at 4°C for 5 minutes. 100 microliters of each lysate were retained as a total RNA control. The remainder of the lysate was then loaded into a 5% (w/v) – 50%(w/v) sucrose gradient prepared in high salt resolving buffer containing 25 mM Tris-HCl (pH 8.0), 0.14 M NaCl, and 10 mM MgCl_2_ using the Gradient Master (Biocomp Instruments), and centrifuged in a Beckman L8 Ultracentrifuge in a SW41 rotor at 41,000 RPM for 1.5 hours at 4°C, with maximum acceleration and braking. For OP50-1 phenformin experiments, gradients were then loaded onto an ÄKTA pure purification FPLC connected with an in-line UV fluorescence sensor coupled to a Bio-Rad chromatography fraction collector, and 10 1mL fractions were collected per gradient. For *egl-45* and *eif-3.B* RNAi experiments, gradients were loaded and fractionated using a Piston Gradient Station with an Triax Flow Cell (Biocomp Instruments) coupled to an automatic 96 tube rack fraction collector (Gilson) with continuous absorbance recording at 260 nm. In either use case, fractions corresponding to the monosome (80S peak) or polysomes (corresponded to all peaks following the monosome) were pooled together, mixed with TriZol LS (Invitrogen), and flash frozen in liquid nitrogen. RNA was extracted from each pooled fraction and the total lysate using the Direct-Zol RNA Miniprep Plut Kit (Zymo Research) and concentrated using the RNA Clean and Concentrator Kit (Zymo Research). rRNA integrity in each fraction was verified using an RNA Tapestation 4000 (Agilent). RNA extracts from the total lysate, monosomal, or polysomal fractions of each sample were then subjected to bulk RNA-sequencing or qRT-PCR analysis, as indicated below.

#### Bulk RNA-sequencing

For bulk RNA-sequencing analyses, 1000 worms of the indicated strain, genotype, or treatment per sample and replicate was harvested using M9, washed three times to remove bacteria, and worm pellets were resuspended into 500 microliters of RNAzol RT. Total RNA was subsequently extracted and the genomic DNA was removed using the Direct-Zol RNA Miniprep Plus Kit (Zymo Research) following manufacturer’s instructions. Total extracted RNA was evaluated for quality control using a NanoDrop One C Microvolume UV-Vis Spectrophotometer (Thermo Fisher). Samples were sent to Azenta (Genewiz) for additional quality control, library preparation, and mRNA sequencing. Samples were validated for RNA integrity with a RIN score > 9 and DV200 > 70 using an RNA Tapestation 4200 (Agilent). Illumina library preparation was performed using polyA selection for mRNA species. Approximately 20 million paired-end 150bp reads were generated per sample, with ≥ 80% of bases passing a Phred quality score ≥ Q30.

#### Lifespan analyses

Lifespans were performed at 20°C except where indicated and performed as previously published^124^. Unless otherwise noted, synchronized L1 animals were seeded onto NGM plates and allowed to grow until the L4/YA stage. On day 0 as indexed in the figure legends, approximately 50 L4 animals were transferred onto fresh NGM or RNAi plates, as indicated. These plates were supplemented with 100 µM FUDR to suppress progeny production (unless otherwise noted). Unless otherwise noted, all lifespan experiments were performed with post-developmental exposure to RNAi or drug treatment (vehicle, phenformin, metformin, malate, glucose, or oligomycin). All drug treatments or FUDR supplementations were performed the same day as L4 transfer, with vehicle or drugs added to the top of prepared NGM or RNAi plates and allowed to dry for 1 hour under laminar flow. For metformin treatment experiments, metformin was added to NGM or RNAi agarose LE plates, as indicated previously^121^. For experiments performed with the use of FUDR, animals were manually transferred to freshly seeded RNAi and/or drug supplemented plates every 2 days between day 0 and day 10 of adulthood, ensuring no crossover contamination of progeny or laid eggs until the animals ceased reproduction. Dead worms were scored and removed every other day (as indicated by the data points), and scoring investigators were blinded to the experimental group/treatment until the conclusion of each experiment. All lifespans performed included matched, same day N2 wild-type controls examined simultaneously with experimental test subjects in each study. All experiments involved a minimum of 2-3 biologically independent experiments, with each biological replicate containing 3 plates of approximately 50 animals each for each condition within the experiment. All lifespan statistics and number of animals scored are included in this paper as **Table S3**.

#### Quantitative fluorescence analysis

For all fluorescence imaging studies, animals of the indicated stage, genotype, or treatment were manually transferred to a droplet of 1 mg/mL levamisole in M9 buffer on top of a 2% (w/v) agarose pad embedded on a microscope slide. Animals were observed for cessation of thrashing and then covered with a coverslip immediately prior to imaging. For whole worm fluorescence imaging studies, slides were imaged at 5x magnification on a Leica DM6 B microscope with Thunder Imaging. For lipid droplet imaging studies, slides were imaged on a Zeiss LSM 800 confocal microscope equipped with an Airyscan detector, centering the posterior intestinal cell nuclei under DIC imaging. Animals were paralyzed and mounted immediately prior to imaging to prevent fluorescence bleaching or excessive levamisole exposure.

#### Fatty acid methyl ester GC/MS lipidomics

Lipid extraction and GC/MS of extracted, acid-methanol-derivatized lipids was performed as described previously ^53,125^. Briefly, 10,000 *C. elegans* of the indicated strain, genotype, stage, or treatment were harvested using M9, and washed three times with fresh M9 to remove bacteria. Worm pellets were then sonicated in minimal amounts of M9 at 70% amplitude using a QSonica Q800R water batch sonicator at 4°C for 20 minutes using a 30 second on, 30 second off cycle. 10 microliters of each sample were then retained for protein concentration analysis. The remainder of the lysate was then combined in a chloroform:methanol mixture, spiked with a FA C17:0 phospholipid and FA C13:0 triglyceride standard mixture, organic phase extracted and dried under nitrogen gas, and then separated into triacylglyceride, cholesteryl esters, and phospholipids using silica-gel solid phase extraction in a gravity manifold. Triglyceride and phospholipid fractions were retained and dried under nitrogen gas. Acidified methanol was added to each lipid fraction and fatty acid methyl esters were subsequently derivatized in a 75°C water bath for 1 hour. Derivativized lipids were then extracted into a hexane and frozen at -80°C for subsequent gas chromatography/mass spectrometry (GC/MS) analysis. Samples were loaded onto an Agilent GC-MS model 6890/5973N outfitted with a Supelcowax-10 (24079) 30 M, 250 μm fused silica capillary column. Samples were injected into a splitless inlet set at 250 °C and 13.33 pounds per square inch. Ultrapure helium was used a carrier gas, with a 1 mL per minute flow rate. Protocol used was held 2 minutes at 150°C, ramp rate at 10°C per minute until 200°C, held for 4 minutes, ramped 5°C per minute until 240°C, held for 3 minutes, ramped 10°C per minute until 270°C, and then held for 5 minutes. A 3 minute solvent delay was used, following detection of masses between 50-550 Da with a MS quadrupole set at 150°C and a MS source detection set at 230°C, with an auxiliary temperature of 270°C. Mass spectra and retention times were extracted, annotated, and area under the curves quantified using the OpenChrom/Eclipse software, using the GLC-87 (Nu-Check Prep) standard mixture as a reference. Fatty acids were quantified as the relative percentage of the total quantified fatty acid pool.

#### Isotopic fatty acid ^13^C_2_ acetate labeling

For hot acetate labeling experiments, animals were treated from L1 of hatching with mock or 10 mM sodium ^13^C_2_ acetate (99% atomic labeling) as indicated in the **Key Resources Table**. 10,000 animals of the indicated stage, genotype, strain, or treatment were harvested in M9 from 10 cm NGM agar plates. Animals were rinsed three times with fresh M9 to remove residual bacteria. Worm pellets were then sonicated in minimal amounts of M9 at 70% amplitude using a QSonica Q800R water batch sonicator at 4°C for 20 minutes using a 30 second on, 30 second off cycle. Following sonication, lipids were extracted in 3:1 methanol:methylene chloride following the addition of acetyl chloride in borosilicate glass tubes, which were then derivatized in a 75°C water bath for 1 hr. Derivatized fatty acids were neutralized with 7% potassium carbonate, extracted with hexane, and washed with acetonitrile prior to evaporation under nitrogen. Lipids were resuspended in 200 µL of hexane and analyzed with GC/MS as described above. Mass spectra and retention times were extracted, annotated, and area under the curves quantified using the OpenChrom/Eclipse software, using the GLC-87 (Nu-Check Prep) standard mixture as a reference. The m/z and mass counts for identified palmitate (FA C16:0) and stearate (FA C18:0) peaks were extracted, and mass counts corresponding to 270-286 m/z were isolated for palmitate peaks, and 298-316 for stearate peaks. Given the natural abundance of ^13^C in both the worm and bacterial derived lipid stores, representation of isotopically labeled carbon labeling in stearate was primarily analyzed and relative abundances calculated as previously described^72^.

#### Intracellular LC-MS metabolomics

Intracellular unlabeled liquid chromatography-mass spectrometry (LC-MS) metabolomics was performed as previously described^126^. Briefly, exactly 1000 worms for each replicate of the indicated genotype, stage, or treatment conditions were counted, manually picked and harvested in M9 buffer, and washed three times to remove residual metabolite treatment or bacterial food source from the plates. Worms were then transferred into a prechilled microcentrifuge tube and metabolites were extracted using a solution containing HPLC-grade 40% (v/v) methanol, 40% (v/v) acetonitrile, and 20% (v/v) ultrapure distilled water. Samples were sonicated at 70% amplitude using a QSonica Q800R water batch sonicator at 4°C for 20 minutes using a 30 second on, 30 second off cycle. Samples were then spun down at 14,000 RPM at 4°C for 20 minutes to pellet cuticle and cellular debris. The supernatant was then transferred into a fresh microcentrifuge tube, and 5 µL of each sample was loaded onto a ZIC-pHILIC column (Millipore). Mass spectra were acquired on a Thermo Q-Exactive Plus run in polarity switching mode with a scan range of 70-1000 m/z and a resolving power of 70,000 at 200 m/z. For each run, the total flow rate was 0.15 mL/min and the samples were loaded at 80% B. The gradient was held at 80% B for 0.5 min, then ramped to 20% B over the next 20 min, held at 20% B for 0.8 min, ramped to 80% B over 0.2 min, then held at 80% B for 7.5 min for re-equilibration. Data were subsequently extracted using Xcalibur (v.4.1.31.9) and analyzed using TraceFinder (v.4.1) and Progenesis (v.2.3.6275.47961).

#### RNAi interference (RNAi)

All RNAi experiments were conducted using *E. coli* HT115(DE3) (Ahringer and Vidal Unique RNAi Libraries) as the food source on RNAi agar or RNAi agarose LE (for use in metformin related experiments) 6 cm plates supplemented with 5 mM IPTG and 200 µg/mL carbenicillin^121^. Cultures of *E. coli* HT115(DE3) were grown in LB broth for 18h at 37°C with 220 RPM shaking incubation and subsequently concentrated to 1/5^th^ of the volume via centrifugation and replenishment with fresh LB broth supplemented with 200 µg/mL carbenicillin. 300 µL of concentrated RNAi bacterial cultures were then seeded onto RNAi plates, dried under laminar flow for 2 hours, and subsequently used for experiments within 1-3 days after seeding. All RNAi clones used in this study were sequence verified prior to use in experiments using either Sanger sequencing through the MGH DNA Core or whole plasmid sequencing through Plasmidsaurus.

#### Quantitative RT-PCR (qRT-PCR)

qRT-PCR was performed as previously described ^15,124^. Approximately 1000 synchronized animals of the stage and treatment conditions indicated in the figure legends were harvested using M9 buffer and washed an additional 3x times to remove residual feeding bacteria. If not processed immediately, nematodes were flash frozen in liquid nitrogen and store at -80°C until RNA isolation. Animals were homogenized using a Qiagen TissueLyzer II with metal beads, centrifuged for 10 minutes at 12,000xg to pellet cuticle and cellular debris, and RNA was extracted from the supernatant using RNAzol RT (Molecular Research Center) according to manufacturer instructions. RNA was treated with RNase free DNAse prior to reverse transcription with the Quantititect Reverse Transcription kit (Qiagen). qRT-PCR was conducted using a Quantitect SYBR Green PCR reagent (Qiagen) following manufacturer instructions on a Bio-Rad CFX96 Real-Time PCR system (Bio-Rad). Expression levels of tested genes were presented as normalized fold changes to the abundance control genes (as indicated in the Figure Legends) using the ΔΔCt method^127^. Primers use for qRT-PCR as included in **Table S6**.

#### Immunoblotting

Immunoblotting was performed as previously described ^124^. Worm and cell lysates were collected into 1x RIPA buffer (containing 50 mM Tris-HCl [pH 7.4], 150 mM NaCl, 1% (v/v) Triton X-100 detergent, 1% sodium deoxycholate, 0.1% sodium dodecyl sulfate (SDS), 1 mM EDTA) supplemented with EDTA-free protease inhibitor tablets (Roche) and a phosphatase inhibitor cocktail containing sodium fluoride, sodium pyrophosphate, sodium orthovanadate, β-glycerophosphate, and okadaic acid. Samples were sonicated at 40% amplitude in a QSonica Q800R3 water bath sonicator at 30 s ON/30 s OFF intervals for a total of 10 minutes at 4°C. Lysates were cleared by centrifugation at 21,000xg at 4°C for 10 minutes and supernatants were retained in clean microcentrifuge tubes for protein analysis. Protein concentration quantification was performed using a Pierce BCA Protein Assay Kit (Thermo Fisher) and samples were diluted to equal concentration using RIPA lysis buffer. Lysates were diluted in 4x Laemmli Sample Buffer (Bio-Rad) supplemented with 10% (v/v) β-mercaptoethanol and boiled for 5 minutes in a 95°C dry bath prior to SDS-PAGE. SDS-PAGE was performed using the Bio-Rad Mini Protean Tetra Cell gel system with 4-15% Mini-PROTEAN TGX precast gels at 150V, using Bio-Rad Precision Plus Dual Color standards. Electrophoretic transfer to nitrocellulose membranes was performed at 100V for 1 hour at 4°C using a 20% (v/v) methanol-based transfer solution. Membranes were then stained for protein abundance using Ponceau S solution, cut using clean scissors, and rinsed twice with 1x TBST (containing 0.1% (v/v) Tween 20 detergent) to remove excess stain. Membranes were blocked using a 5% non-fat dry milk solution in 1x TBST for 1 hour rocking at room temperature, rinsed 3 times with 1x TBST, and then incubated with primary antibody, rocking for 18 hours at 4°C and diluted according to manufacturer’s guidelines. Membranes were then washed 3 times with 1x TBST. For primary antibody immunodetection, a 1:5000 dilution of Goat anti-Rabbit HRP conjugate or Goat anti-Mouse HRP conjugate (GE Heathcare) in a 5% bovine serum albumin (BSA) 1x TBST solution was used, incubating membranes for 1 hour rocking at room temperature. Membranes were washed 3 times with 1x TBST, and HRP detection was performed using the SuperSignal West Pico PLUS chemiluminescent substrate. Chemiluminscence was detected and imaged on membranes using the iBright imaging system (Thermo Fisher Scientific). Images were obtained using a chemiluminescence and membrane overlay and compiled using Adobe Illustrator and Microsoft Office PowerPoint.

#### Ribonucleoprotein (RNP) immunoprecipitation and qRT-PCR analysis (RIP-qRT-PCR)

Immunoprecipitation was performed using 30,000 animals per sample bearing a transgene expressing *myo-2p::NLS::GFP, eif-3.Bp::eif-3.B::RFP::unc-54 3’ UTR’, rps-0p: hygromycin:tbb-2 3’ UTR* synchronized to the L4/young adult stage, treated with or without 4.5 mM phenformin from L1 of hatching. Animals were harvested in M9 and washed three times with fresh M9 to remove excess bacteria. Worm pellets were aspirated of excess M9 and resuspended in 200 microliters of lysis buffer containing 10 mM Tris/Cl (pH 7.5), 150 mM NaCl, 0.5 mM EDTA, 0.5 % Nonidet-P40, 125 U/microliter of SUPERaseIn RNAse inhibitor, and supplemented with EDTA-free protease inhibitor tablets (Roche) and a phosphatase inhibitor cocktail described above. Samples were then sonicated at 40% amplitude in a QSonica Q800R3 water bath sonicator at 30 s ON/30 s OFF intervals for a total of 10 minutes at 4°C. Samples were then cleared by centrifugation at 21,000xg at 4°C for 5 minutes and supernatants were retained in clean microcentrifuge tubes for protein analysis by BCA as described above. All samples were diluted in lysis buffer to equivalent protein concentration, and 50 microliters of each sample were saved as a total IP control for comparison. RFP-Trap and Blocking Control magnetic agarose beads (Bulldog Bio) were equilibrated for each sample in ice cold dilution buffer containing 10 mM Tris/Cl (pH 7.5), 150 mM NaCl, 0.5 mM EDTA, 125 U/microliter of SUPERaseIn RNAse inhibitor, and supplemented with EDTA-free protease inhibitor tablets (Roche) and a phosphatase inhibitor cocktail described above. Equilibrated beads were then incubated with half of each sample (to ensure a Blocking Control was performed for each sample incubated with RFP-Trap), rotating overnight at 4°C. 50 microliters of each sample (Blocking/RFP-Trap) were retained as a flow-through control. Beads were washed three times using ice cold wash buffer containing 10 mM Tris/Cl (pH 7.5), 150 mM NaCl, 0.5 mM EDTA, 0.05% Nonidet-P40, 125 U/microliter of SUPERaseIn RNAse inhibitor, and supplemented with EDTA-free protease inhibitor tablets (Roche) and a phosphatase inhibitor cocktail described above. Beads were then split into two pools for immunoblotting or qRT-PCR. For immunoblotting, protein was eluted from beads with resuspension in 40 microliters of 4x Laemmli Sample Buffer (Bio-Rad) supplemented with 10% (v/v) β-mercaptoethanol and boiled for 5 minutes in a 95°C dry bath prior to SDS-PAGE as indicated above. For qRT-PCR analysis, beads were eluted in RNAzol RT according to manufacturer’s instructions. cDNA synthesis and qRT-PCR plate preparation was performed as indicated above. Data was normalized to *pmp-3* as the reference expression control and the blocking IP and vehicle treated sample as the reference sample control using the ΔΔCt method^127,128^. Primers used for RNA immunoprecipitation/qRT-PCR analysis as included in **Table S6**.

#### Fixative Nile red staining

Fixative-based Nile red staining was conducted as indicated previously ^129^. Briefly, approximately 1000 synchronized animals of the indicated treatment, genotype, or condition were washed off plates using M9 buffer supplemented with 0.01% Triton X-100 detergent (M9T) and washed 3 times with fresh M9T to remove excess bacteria. Animals were then fixed in a 40% isopropanol solution for 3 minutes under rotation and then stained with a dilute 0.5 mg/mL Nile Red stock solution in 40% isopropanol (6 uL of stock solution in 1 mL of 40% isopropanol). Samples were incubated in staining for 2 hours, shaking, in the dark. Samples were destained using M9T for 30 minutes shaking in the dark, washed once more with M9T, and then worms were mixed and dispensed onto microscope slides and sealed under a coverslip with clear nail polish, and dried for 5 minutes in the dark. Approximately 50 animals per replicate and per condition were then imaged at 10x with brightfield, DIC and GFP (AlexFluor4888-Green) filters using a Leica DM6 B Upright LED microscope with Thunder Imaging capability.

#### ThioFluor 623 staining

Staining of endogenous reduced thiols in living *C. elegans* nematodes was performed as previously described^60^ with the following modifications. Animals were manually picked from and immersed in sterile M9 buffer. Animals were washed three times with fresh M9 to remove bacteria and then incubated in 25 µM ThioFluor 623 staining solution prepared in M9. Animals were rotated in staining solution for 1 hour at 20°C, and then subsequently washed of excess staining solution with fresh M9 three times. Animals were then transferred to a 2% (w/v) agarose pad on a microscope slide, immobilized with levamisole, sealed with a coverslip, and then immediately imaged at 10x with brightfield, DIC, and RFP filters using a Leica DM6 B Upright LED microscopy with Thunder Imaging capability.

#### Generation of C. elegans transgenics

Transgenic overexpression of mitochondrial and cytoplasmic LbNOX was performed by subcloning each LbNOX sequence from pUC57-LbNOX or pUC57-mitoLbNOX plasmids into pAS59[sur-5p::MCS2_::SL2::GFP::unc-54 3’ UTR] via Gibson assembly using NheI and KpnI restriction digests and oligonucleotides AS-3260, AS-3261, and AS-3262. Extrachromosomally arrayed animals were generated via injection of 5 ng/µL of each construct (pAS108 or pAS109) with 10 ng/µL of coel::RFP injection marker (Addgene Cat# 8938), and 85 ng/µL of ultrapure salmon sperm DNA (Thermo Fisher) for a final concentration of 100 ng/µL of total DNA material. Mixtures were injected into the gonads of wild-type animals, and progeny were assessed for array retention using GFP and RFP fluorescence.

Transgenic overexpression of mitochondrial and cytoplasmic TPNOX was performed by subloning an GENEFragment of mitoTPNOX and cytoTPNOX (AS-8603 and AS-8604) individually into pAS59 via Gibson assembly using NheI and KpnI restriction digests and oligonucleotides AS-8600, AS-8601, and AS-8602. Extrachromosomally arrayed animals were generated via injection of 10 ng/uL of each construct (pAS508 or pAS509) with 5 ng/µL of pCFJ90 (*myo-2p::mCherry*) co-injection marker (Addgene Cat# 19327), 10 ng/µL of pAS351 (*rps-0p: hygromycinR:tbb-2 3’ UTR*) and 75 ng/µL of salmon sperm DNA for a final concentration of 100 ng/µL of total DNA material. Mixtures were injected into the gonads of wild-type animals, and progeny were assessed for array retention using GFP and RFP fluorescence.

Transgenic overexpression of EIF-3.B was performed via the PCR fusion stitch method^130^. The entire genomic locus of EIF-3.B was amplified via PCR using primers AS-8818 and AS-8819, and stitched in-frame with RFP::HA::STOP::unc-54 3’ UTR from modified Fire vector pPD95.79 with a second round of PCR. PCR fragments were extracted and purified using the QIAquick Gel Extraction Kit (Qiagen). 10 ng/µL of the PCR stitch product along with 5 ng/µL of pAS5 (*myo-2p::NLS::GFP*) co-injection marker, 10 ng/µL of pAS351 and 75 ng/µL of salmon sperm DNA (100 ng/µL total concentration was injected into the gonads of wild-type animals, and progeny were assessed for array retention using GFP and RFP fluorescence.

For generation of the 3xFLAG::AID*::GFP::FASN-1 animal, plasmids bearing *C. elegans* codon optimized Cas9 with an sgRNA targeting the N-terminus of *fasn-1* (pAS558, derived from pAS276 and pDD162, AS-8232 and AS-8233) and a 3xFLAG::AID*::GFP repair template (pAS557, derived from pJW1583, AS-8235 to AS-8238) were cloned via SapI digestion and Gibson assembly. 50 ng/µL of pAS558, 50 ng/µL of pAS557, 10 ng/µL of pGH8 (*rab-3p::mCherry*) co-injection marker (Addgene Cat# 19359), 5 ng/µL of pCFJ104 (*myo-3p::mCherry*) co-injection marker (Addgene Cat# 19328) and 2.5 ng/µL of pCFJ90 co-injection marker were injected into the gonads of wild-type animals. Knock-in editing selection and validation was performed as previously described ^131^.

For generation of COX-6A::GFP::3xFLAG animals, plasmids bearing *C. elegans* codon optimized Cas9 with an sgRNA targeting the C-terminus of *cox-6a* (pAS556, derived from pAS276 and pDD162, AS-8913 and AS-8914) and a GFP::3xFLAG repair template (pAS555, derived from pDD282, AS-8927 to AS-8930) were cloned via SapI digestion and Gibson assembly. 50 ng/µL of pAS556, 50 ng/µL of pAS555, 10 ng/µL of pGH8 co-injection marker, 5 ng/µL of pCFJ104 co-injection marker and 2.5 ng/µL of pCFJ90 co-injection marker were injected into the gonads of wild-type animals. Knock-in editing selection and validation was performed as previously described ^131^.

For generation of FAT-3::GFP::3xFLAG animals, plasmids bearing *C. elegans* codon optimized Cas9 with an sgRNA targeting the C-terminus of *fat-3* (pAS554, derived from pAS276 and pDD162, AS-8907 and AS-8908) and a GFP::3xFLAG repair template (pAS553, derived from pDD282, AS-8915 to AS-8918) were cloned via SapI digestion and Gibson assembly. 50 ng/µL of pAS554, 50 ng/µL of pAS553, 10 ng/µL of pGH8 co-injection marker, 5 ng/µL of pCFJ104 co-injection marker and 2.5 ng/µL of pCFJ90 co-injection marker were injected into the gonads of wild-type animals. Knock-in editing selection and validation was performed as previously described ^131^.

Two to three independently injected transgenic lines were generated for each extrachromosomal array and independently validated for transgene and co-injection marker expression. Animals with hygromycin resistance transgene expression were maintained on NGM plates freshly supplemented with 50 µg/mL hygromycin B and passaged once off hygromycin prior to synchronization and experimentation. All plasmids were sequence verified through Plasmidsaurus, Inc. and all animals were genotype verified prior to experimentation. For all oligonucleotides used during strain generation and validation, refer to **Table S6** for sequence details.

#### Generation of metabolically inactive E. coli for lipidomic and lifespan studies

PFA killing of OP50-1 E. coli was performed as previously described with slight modifications ^15,51^. 50 mL aliquots of OP50-1 liquid cultures grown overnight in LB media supplemented with 25 μg/mL streptomycin were dispensed into 250 mL Erlenmeyer flasks. Either 1× PBS (Life Technologies) for mock treatment or 4% PFA (Millipore Sigma) diluted in 1× PBS was added to each flask for a final concentration of 1% (vol/vol). Bacteria were then shaken in 37°C at 210 rpm for 2 hr to enable PFA inactivation. Cultures were then aseptically transferred into 50 mL conical centrifuge tubes and then washed 10× with sterile PBS to remove residual PBS or PFA solution. After the final wash, bacterial pellets were then 10× concentrated in LB media supplemented with 25 μg/mL streptomycin, and 300 μL seeded onto freshly prepared NGM plates. Plates were allowed to dry for 2 days prior to use for GC/MS or lifespan analyses. A standard culture of OP50-1 grown overnight was similarly 10× concentrated and seeded as a ‘live OP50-1’ control to compare to mock-treated and PFA-treated bacterial conditions.

#### Cellular viability analysis using crystal violet staining

Crystal violet staining of mammalian cell culture was performed as described by the Xin Chen Lab (UCSF) with minor modifications (https://pharm.ucsf.edu/xinchen/protocols/cell-enumeration). Approximately 100,000 cells for each cell line described were seeded in each well of a 24-well plate and allowed to grow 48 hours under tissue culture conditions described above until they reached approximately 70% confluency. Cells were then treated with compounds as described for 48 hours prior to staining. Cells were aspirated of media and rinsed once with PBS, then incubated for 15 minutes with crystal violet staining solution (0.125g crystal violet in 50 mL of 20% (v/v) methanol). Staining solution was aspirated, and cells were rinsed 5 times with PBS to remove excess stain. Cells were lysed using a 0.1 M sodium citrate solution in 50% (v/v) ethanol (pH 4.2) for 30 minutes with gentle rocking and visually inspected for complete lysis. Stained lysates were then diluted 4x with sterile water, loaded into 96-well plates, and optical density read using a Spectramax plate reader at an emission wavelength of 590 nm. Normalization of results to baseline and statistical testing was performed using Graphpad Prism 10.

### QUANTIFICATION AND STATISTICAL ANALYSIS

#### Lifespan analyses

Lifespan analyses was performed using OASIS2 ^112^ to generate mean lifespan days, standard error, 95% confidence interval (CI), and log rank analyses. Percent survival values were extracted from tabular results and plotted using Graphpad Prism 10. For all tabular lifespan statistical results, refer to **Table S3**.

#### Quantitative fluorescence analyses

Whole worm fluorescence imaging was performed using FIJI/ImageJ2. Raw intensity images were extracted using Leica LAS X software, and FIJI was used to manually outline and trace individual worms. Mean grey values were extracted for each individual worm (normalized to worm surface area) and plotted/statistically analyzed using Graphpad Prism 10. For analysis of lipid droplet diameters, Biodock was used, utilizing a machine leaning algorithm trained on images randomly selected throughput the entire imaging dataset (including all biological and technical replicates). Diameters for each lipid droplet identified were extracted from Biodock and plotted using Graphpad Prism 10. All representative images shown were equivalently scaled using FIJI. Biological replicate assessment and statistical tests used are as noted in the Figure Legends.

#### CLIME and Gene-Module Association Determination analyses

CLIME^62^ was performed using ‘*fasn-1*’ and ‘*pod-2*’ as input *C. elegans* query, with 200 predictions per ECM requested with default parameters utilized otherwise. The top 25 paralogs were selected based on the log likelihood ratio of ECM membership and plotted using Graphpad Prism 10. Gene-Module Association Determination^59^ was performed on GeneBridge (https://systems-genetics.org/) using ‘*fasn-1*’ as the C. elegans query set. Gene-Module Association Scores (GMAS) were calculated using all available transcriptomic datasets. Manhattan plots were extracted from the server and replotted using Graphpad Prism 10.

#### Immunoblotting densitometry analyses

Densitometry analysis was performed using FIJI/ImageJ2. Raw chemiluminescent images for each membrane were extracted using iBright imaging software (Thermo Fisher) and band intensities were inverted using FIJI. Equal rectangular traces were collected for each band (ensuring equivalent area sample) and mean grey values were extracted and plotted using Graphpad Prism 10. Biological replicate assessment and statistical tests used are as noted in the Figure Legends.

#### RNA-sequencing analyses

Fastq read quality control, adapter trimming, quality score filtering, and quasi-alignment were all performed using custom UNIX/Bash shell scripts on the Mass General Brigham ERISTwo Scientific Computing CentOS 7 Linux Cluster. Reads were analyzed for quality control using FastQC v0.11.8^132^ (http://www.bioinformatics.babraham.ac.uk/projects/fastqc) and MultiQC v1.19^133^. Reads were then filtered for Illumina adapter contamination, truncated short reads or low-quality base calls using BBDuk (BBTools)^134^. The subsequently trimmed and cleaned reads were then quasi-aligned to the Caenorhabditis elegans reference transcriptome annotation (WBcel235, Ensembl Release 105) using Salmon v1.9.0^113,135^, correcting for GC content and sequencing bias using the command parameters ‘--gcBias’ and ‘--seqBias’. All statistical analyses and visualizations were generated using the R v4.3.2 (https://www.r-project.org) Bioconductor v3.18 statistical environment on a local machine through Jupyter Notebook v6.4.10 (https://jupyter.org)^116^. Quasi-aligned transcript quantification files for each sample were collapsed into gene-level count matrices using R package tximport v1.30.0^136^.

Analysis of polysome profiling associated RNA-sequencing data was performed using the deltaTE workflow^119^, generating a list of differentially translated genes (DTEGs) using R package DESeq2 v1.42.1101 with a design formula ‘∼ Replicate + Treatment + Fraction + Treatment:Fraction’. DTEGs were selected with an FDR adjusted P value < 0.05. Quantification of the CDS and UTR lengths for upregulated and downregulated DTEGs was calculated using the WBcel235 GTF from Ensembl Release 105^135^. Quantification of gene essential scores was extracted from Campos et al.^137^ Volcano plots and density distribution plots were generated using R package ggplot2 v3.5.0. Protein complex binning of DTEGs was performed using an annotation map of *C. elegans* proteins belonging to established orthologous mammalian protein complexes^48^. Binning of ribosomal subunit and ETC complex proteins was performed using WormPaths and WormBase GO term membership^58^. Cleveland plots were generated using R package ggplot2 v3.5.0.

For analysis of *fasn-1* RNAi treatment experiments, paired differential expression was calculated using R package DESeq2 v1.42.1x with a design formula of ‘∼ RNAi_Treatment’^114^. Genes were considered differentially expressed with a Benjamini-Hochberg False Discovery Rate (FDR) corrected P value < 0.05 and an absolute log_2_ transformed fold change of 1^138^. Metabolism gene set variation analysis (GSVA) was performed using R package GSVA v1.50.1 using metabolic gene sets extracted from WormPaths^58,117^. Comparison of differentially expressed genes with prior *C. elegans* mutant datasets was performed using WormExp v2.0^109^. Tissue expression prediction analysis of differentially expressed genes as indicated in the Figure Legends was performed using the Worm Tissue Prediction Server and data plotted using Graphpad Prism 10^118^. Heatmaps were generated using R package pheatmap v1.0.12. Gene set overrepresentation analyses for genes identified as differentially expressed (with thresholds set as indicated above) were performed using R package clusterProfiler v4.12.3 for GO term-based pathway annotations^115^. Visualizations were post-edited for font, sizing, and appearance using Adobe Illustrator and the Adobe Creative Cloud Suite. All custom bash scripts, Jupyter Notebook files, and processed fastq and transcript count files used in these analyses will be provided upon reasonable request by the corresponding author.

#### Metabolomics analyses

Heatmaps of z-score normalized relative abundances of select metabolites were generated using R package pheatmap v1.0.12. Relative abundance or ratiometric metabolite values were otherwise plotted using Graphpad Prism 10. Biological replicate assessment and statistical tests used are as noted in the Figure Legends.

#### DepMap and CCLE metabolomics integrative analyses

Integrative DepMap^76^ and CCLE metabolomics^74^ data detailing the genetic dependencies most closely linked to metabolomic variance were extracted from a prior publication and replotted using Graphpad Prism 10. Lineage stratified analysis of FASN protein levels (using CCLE proteomics^110^) and growth resistance to metformin treatment (using the Drug Repurposing Study) was performed using Data Explorer v2.0 in the DepMap Portal (https://doi.org/10.25452/figshare.plus.25880521.v1) and replotted using Graphpad Prism 10. FASN expression level alterations in KIRC and KIRC cancers with their associated link to overall survival outcomes were extracted from GEPIA^139^ and replotted using Adobe Illustrator.

#### General statistical analyses

Unless otherwise noted, all values are presented with mean ± SEM and the number (n) of biological replicates performed is indicated in the Figure Legends. All statistical analyses unless otherwise noted was performed in Graphpad Prism 10. The statistical differences between control and a single experiment group were determined by Student’s test, and control and multiple experimental groups were determined by one-way ANOVA, with corrected p values < 0.05 considered significant. Two-way ANOVA with Bonferroni correction for multiple hypothesis testing was performed to determine significant difference where genetic interferences/differential genotypes and drug treatments were applied together, with corrected p values < 0.05 considered significant. Asterisks denote corresponding statistical significance ∗p < 0.05; ∗∗p < 0.01; ∗∗∗p < 0.001 and ∗∗∗∗p < 0.0001.

## SUPPLEMENTAL FIGURE TITLES AND LEGENDS

**Figure S1.**
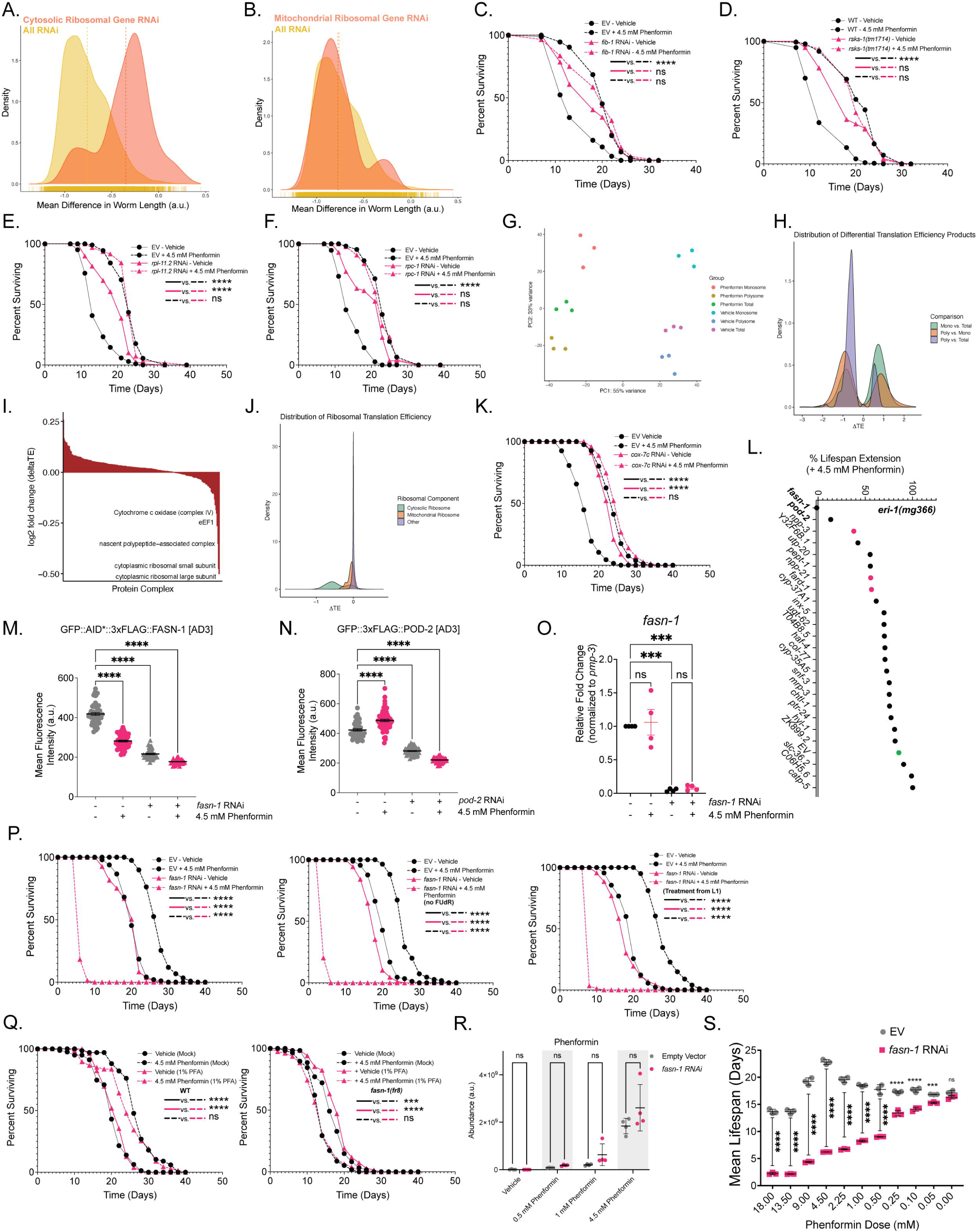
Biguanides partly enforce their growth inhibitory and pro-longevity effects through reduction of mRNA translation. Related to Figure 1. (A-B) Distribution plot of all RNAi clones and their respective impact on metformin-mediated growth inhibition. RNAi distributions were stratified into cytoplasmic (A) or mitochondrial (B) ribosome GO term groups (orange) relative to the average distribution of all RNAi clones assayed (yellow). (C) Lifespan analyses of WT animals treated with *fib-1* RNAi or 4.5 mM phenformin from L4. (D) Lifespan analyses of WT animals and *rsks-1(tm1714)* null animals treated with 4.5 mM phenformin from L4. (E-F) Lifespan analyses of WT animals treated with *rpl-11.2* (E) or *rpc-1* (F) RNAi and 4.5 mM phenformin from L4. (G) PCA analysis of polysomal, monosomal, and total RNA fractions collected with vehicle or phenformin treated WT animals at L4/YA (n = 3 biological replicates). (H) Translation efficiency (TE) shifts upon treatment with 4.5 mM phenformin when comparing polysomal occupancy to total RNA expression (purple), polysomal to monosomal RNA occupancy (orange), and monosomal occupancy to total RNA expression (green) (n = 3 biological replicates). (I) Genes ranked on TE shifts with phenformin treatment, with values combined based on known protein complex membership^48^. (J) TE shifts with phenformin treatment in cytoplasmic (green) versus mitochondrial (orange) ribosomal genes. (K) Representative lifespan analysis of WT *C. elegans* animals treated from L4 with *cox-7C* RNAi and 4.5 mM phenformin. (L) Curated lifespan screen using RNAi hypersensitive *eri-1(mg366)* animals, targeting genes with increased TE upon 4.5 mM phenformin treatment relative to EV control (green). Genes highlighted in red were used as positive controls from previously established studies^14,15^. (M-O) Validation of RNAi knockdown efficiency for *fasn-1* (M, O) or *pod-2* (N) RNAi using endogenously GFP-tagged FASN-1 (M), POD-2 (N), or qRT-PCR RNA analysis of WT animals treated with *fasn-1* RNAi or 4.5 mM phenformin from L4 to AD3 (N). Note that the EV measurements in S1M are identical to those shown in S2C, and EV measurements in S1N are identical to those shown in 2I, as these quantifications were performed in parallel. For (O), n = 4 biological replicates. (P) Lifespan analyses for WT animals treated with *fasn-1* RNAi or 4.5 mM phenformin from L4 with FUdR (left), from L4 without FUdR (middle), or from L1 with FUdR (right). (Q) Lifespan analyses for WT (top) or hypomorphic *fasn-(fr8)* (bottom) animals treated with 4.5 mM phenformin from L4, fed with *E. coli* OP50-1 pre-treated with mock or 1% (v/v) PFA for 2 hours. (R) LC-MS analysis of phenformin levels in animals treated with *fasn-1* RNAi or increasing doses of phenformin (n = 4 biological replicates). (S) Mean lifespan of animals treated with *fasn-1* RNAi and subjected to varying doses of phenformin from L4. Lifespans are representative of 3 biological replicates (Table S3). Data represent the mean ± SEM. Imaging data are representative of at least 30 animals aggregated from 3 biologically independent experiments per condition, unless otherwise noted. Significance was assessed by (C-F, K, P, Q) log rank analyses and (M-O, R-S) two-way ANOVA followed by correction for multiple comparisons. ns p > 0.05, * p < 0.05, ** p < 0.01, *** p < 0.001, **** p < 0.0001.

**Figure S2.**
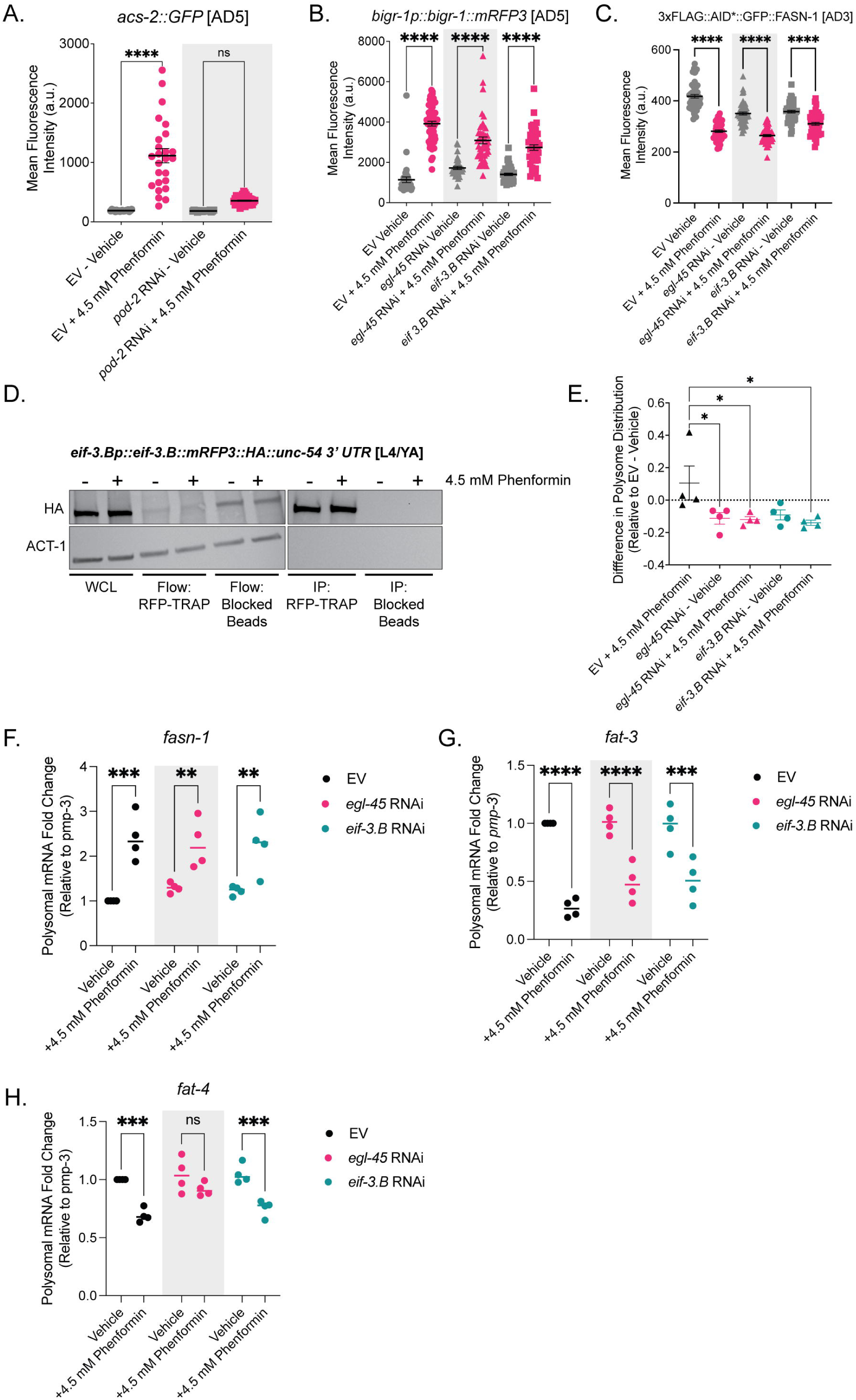
EGL-45 and EIF-3.B depletion does not act as a transgene suppressor, and does not regulate FASN-1 protein synthesis or polysome occupancy. Related to Figure 2. (A) GFP quantification of *acs-2p::GFP* animals treated with *pod-2* RNAi or 4.5 mM phenformin from L4 until AD5. (B-C) GFP quantification of *bigr-1p::bigr-1::mRFP3* (B) and 3xFLAG::AID*::GFP::FASN-1 (C) animals treated with *egl-45* or *eif-3.B* RNAi or 4.5 mM phenformin from L4 until AD5 (B) or AD3 (C). Note that the EV measurements in S2C are identical to those shown in S1M, as these quantifications were performed in parallel. (D) Representative western blot validation of RNA immunoprecipitation strategy shown in 2N. Left blot represents EIF-3.B::mRFP3 enrichment in whole cell lysates that is significantly depleted in RFP IP conditions relative to blocking bead controls. Purity of EIF-3.B mRFP3 pull down is shown in the right blot. Representative of 3 biological replicates. (E) Difference in distribution of ribosomes on polysomes relative to EV - Vehicle of WT animals treated with *egl-45* or *eif-3.B* RNAi or 4.5 mM phenformin from L4 to AD3 (n = 4 biological replicates). (F-H) qRT-PCR expression analysis of *fasn-1, fat-4,* and *fat-3* from pooled polysomal fractions of WT *C. elegans* treated as shown in (E) (n = 4 biological replicates). Lifespans are representative of 3 biological replicates (Table S3). Data represent the mean ± SEM. Imaging data are representative of at least 30 animals aggregated from 3 biologically independent experiments per condition, unless otherwise noted. Significance was assessed for (A-C, E-H) using two-way ANOVA followed by correction for multiple comparisons. ns p > 0.05, * p < 0.05, ** p < 0.01, *** p < 0.001, **** p < 0.0001.

**Figure S3.**
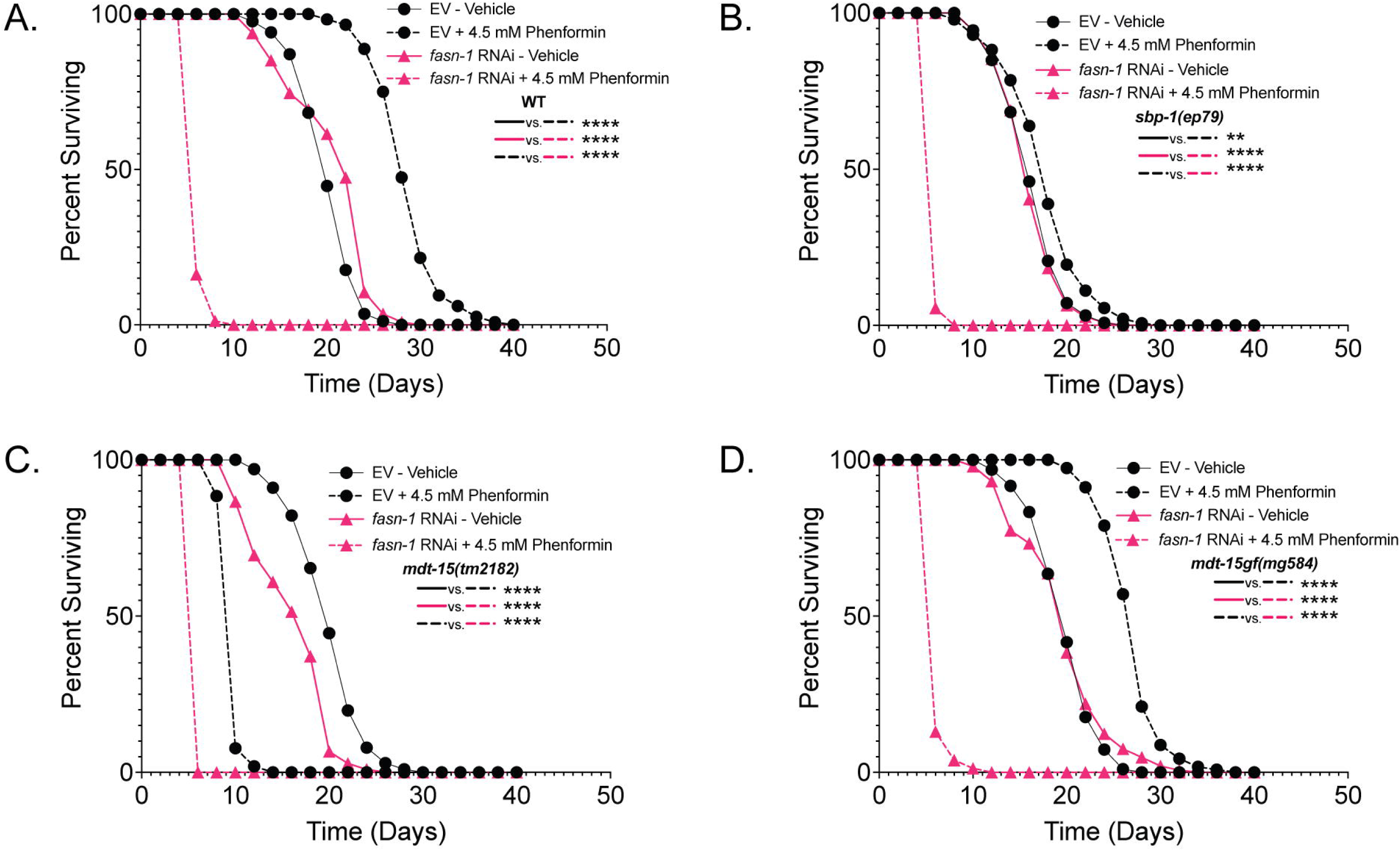
Established lipid synthesis inhibition paradigms do not mimic *fasn-1*/*pod-2* RNAi-induced BIG-SAD. Related to Figure 3. (A-D) Lifespan analyses of WT (A), *sbp-1(ep79)* (B), *mdt-15(tm2182)* (C), and *mdt-15gf(mg584)* (D) animals treated with *fasn-1* RNAi or 4.5 mM phenformin from L4. Lifespans are representative of 3 biological replicates (Table S3). Significance was assessed for (A-D) using log rank analysis. ns p > 0.05, * p < 0.05, ** p < 0.01, *** p < 0.001, **** p < 0.0001.

**Figure S4.**
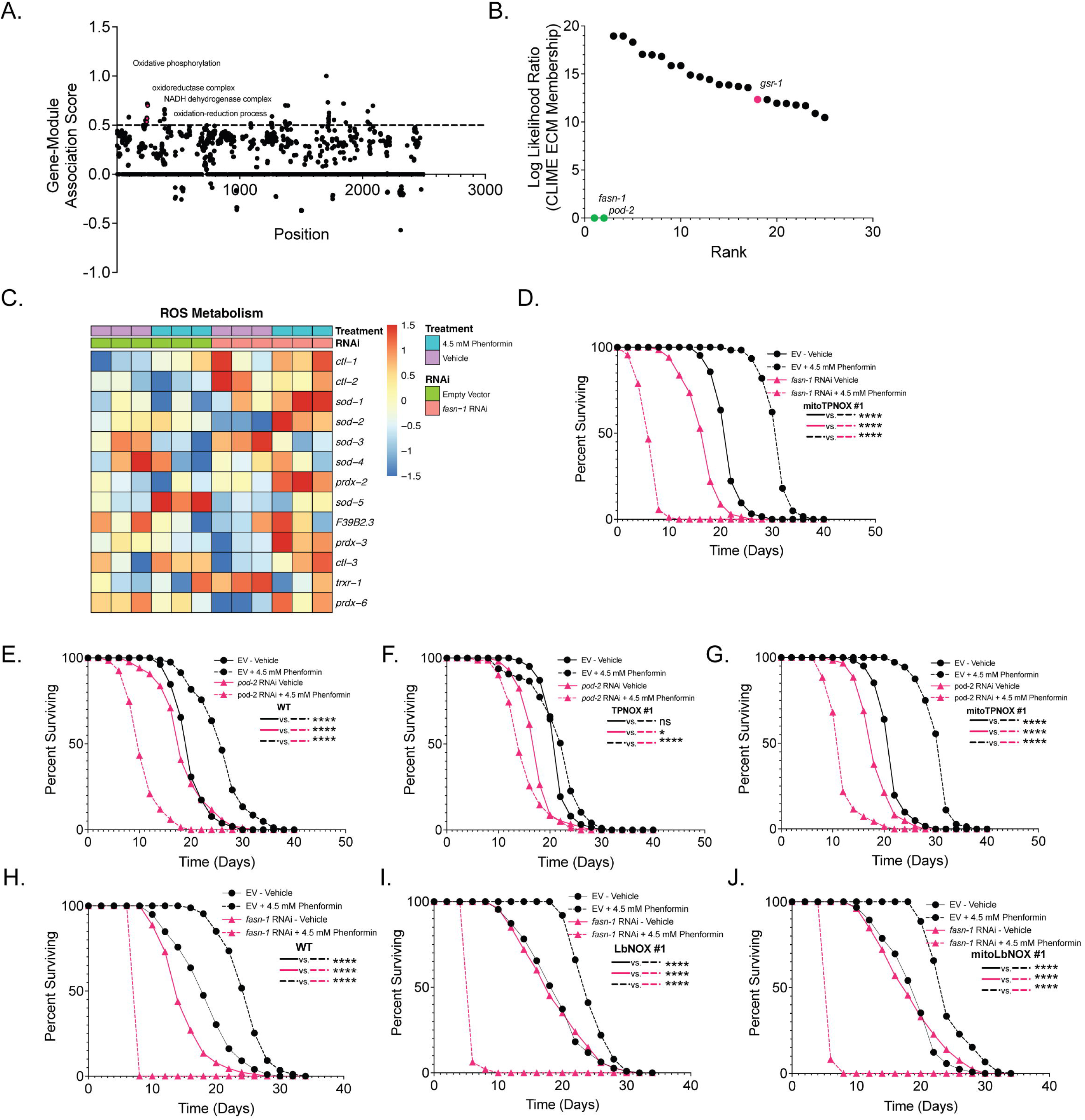
FASN-1 fine tunes redox homeostasis and ROS metabolism during biguanide response. Related to Figure 4. (A) GMAS of FASN-1 expression with alterations in GO Molecular Function terms. Data were analyzed using the GeneBridge analysis platform. Dotted line represents a significant gene-module association score above 0.5. (B) CLIME^62^ evolutionary conservation association analysis of *fasn-1* and *pod-2*. The top 25 genes associated by ECM membership and ranked by log likelihood score are presented here. *gsr-1* is highlighted in red.(C) Gene expression of transcripts related to ROS metabolism from WT animals treated with *fasn-1* RNAi or 4.5 mM phenformin from L4 to AD3 (n = 3 biological replicates). ROS metabolism gene sets were extracted from WormClust^58^. (D-G) Lifespan analysis of mitoTPNOX (D,G), WT (E), or cytoplasmic TPNOX (F) overexpressing animals with *fasn-1* RNAi or *pod-2* RNAi or 4.5 mM phenformin treatment from L4. (H-J) Lifespan analyses of WT (H), *sur-5p::LbNOX::FLAG::SL2::GFP::unc-54 3’UTR* (I) or *sur-5p::mitoLbNOX::FLAG::SL2::GFP::unc-54 3’UTR* (J) animals treated with *fasn-1* RNAi or 4.5 mM phenformin from L4. Lifespans are representative of 3 biological replicates (Table S3). Significance was assessed for (D-J) using log rank analyses. ns p > 0.05, * p < 0.05, ** p < 0.01, *** p < 0.001, **** p < 0.0001.

**Figure S5.**
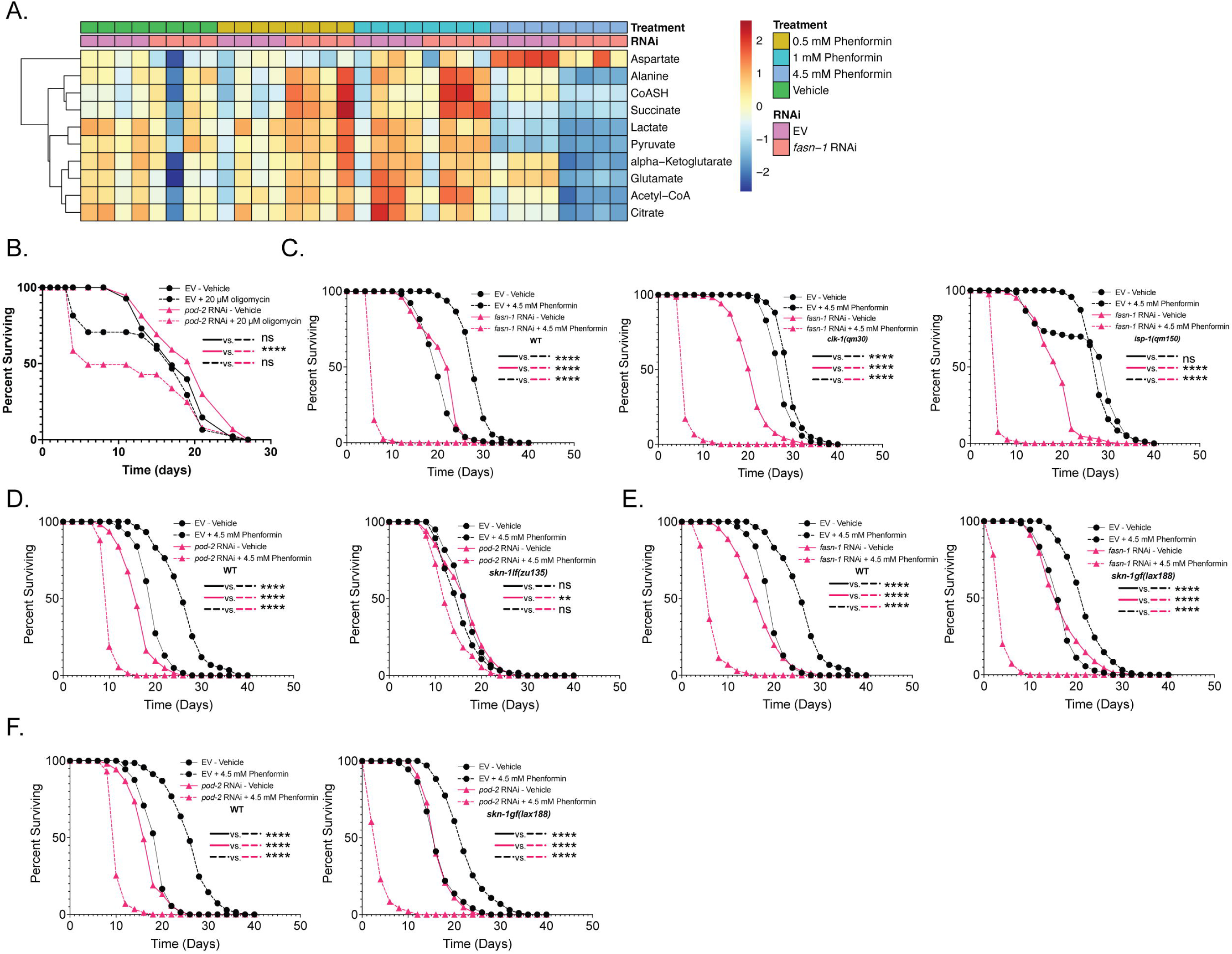
Manipulating the response to reductive stress controls the onset of BIG-SAD. Related to Figure 5. (A) Visualization of select TCA cycle and amino acid metabolites from LC-MS analysis of WT animals treated with *fasn-1* RNAi and varying doses of phenformin (n = 4 biological replicates). (B) Lifespan analyses of WT animals treated with *pod-2* RNAi or 20 µM oligomycin from L4. Lifespan is representative of 4 biological replicates. (C) Lifespan analyses of WT (left), *clk-1(qm30)* (middle), or *isp-1(qm150)* (right) animals treated with *fasn-1* RNAi or 4.5 mM phenformin from L4. (D) Lifespan analysis of WT (left), or *skn-1lf(zu135)* (right) animals treated with *pod-2* RNAi or 4.5 mM phenformin from L4. (E-F) Lifespan analyses of WT (E-F left) or *skn-1gf(lax188)* (E, F right) animals treated with *fasn-1* (E) or *pod-2* RNAi (F) or 4.5 mM phenformin from L4. Lifespans are representative of 3 biological replicates (Table S3). Significance was assessed for (B-F) using log rank analyses. ns p > 0.05, * p < 0.05, ** p < 0.01, *** p < 0.001, **** p < 0.0001.

**Figure S6.**
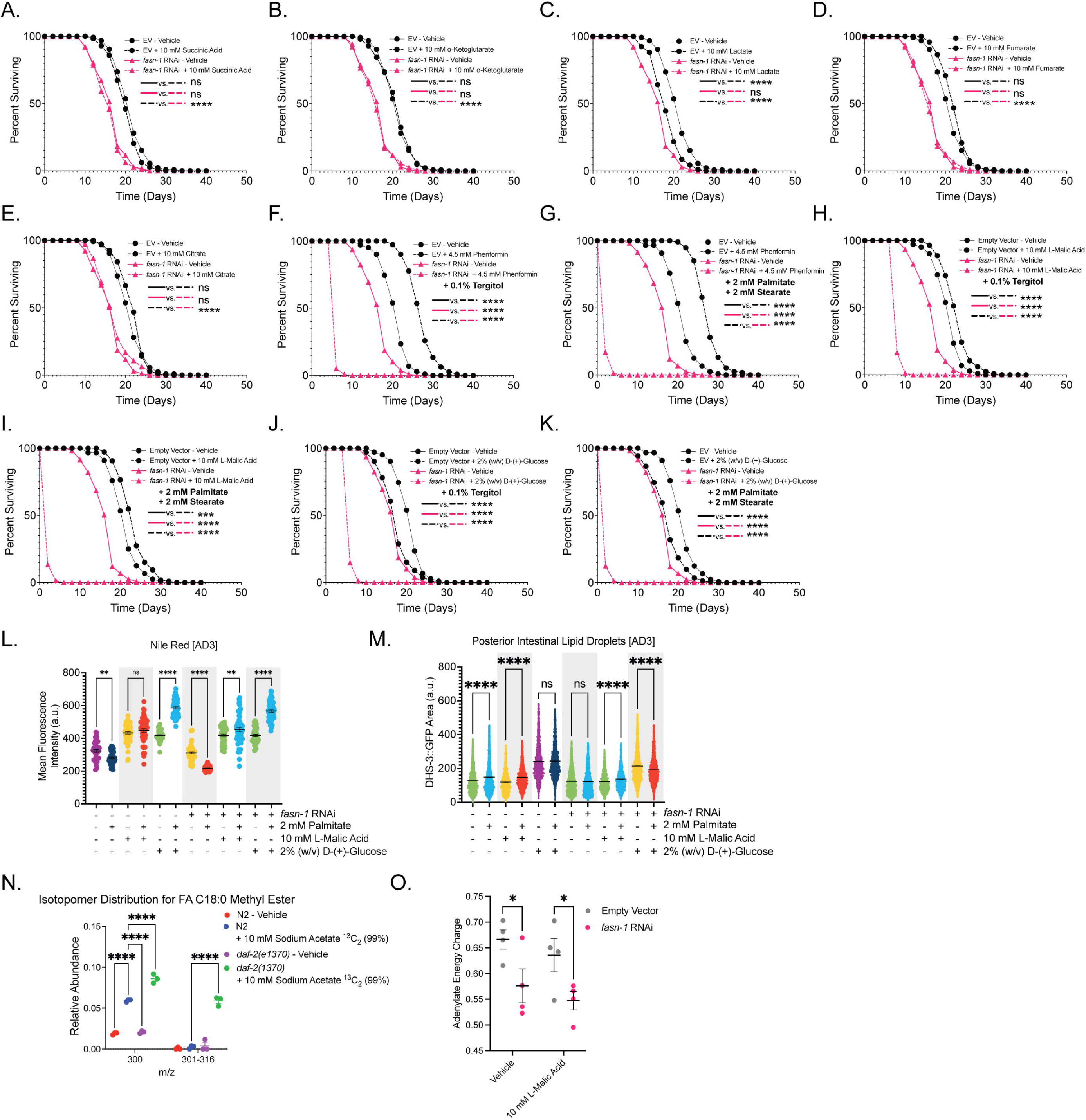
Exogenous malate and glucose supplementation is not rescued by supplementation with downstream lipid products. Related to Figure 6. (A-E) Lifespan analyses of WT animals treated with *fasn-1* RNAi or 10 mM succinic acid (A), 10 mM alpha-ketoglutarate (B), 10 mM lactate (C), 10 mM fumarate (D), or 10 mM citrate (E). Note that these lifespan analyses were performed in tandem and thus contain identical vehicle treatment curves in these representative plots. (F-K) Lifespan analyses of WT animals treated with *fasn-1* RNAi or 4.5 mM phenformin (F-G), 10 mM L-malic acid (H-I), or 2% (w/v) D-(+)-glucose (J-K) and supplemented with either 0.1% tergitol control (F, H, J) or 2 mM palmitate and 2 mM stearate (G, I, K). (L-M) Quantification of Nile Red stained neutral lipid stores (L) or DHS-3::GFP labeled posterior intestinal lipid droplet area (M) of animals treated with *fasn-1* RNAi and 10 mM L-malic acid or 2% (w/v) D-(+)-glucose on plates supplemented with 2 mM palmitate. Note that the EV/*fasn-1* RNAi ± 2 mM palmitate measurements in S6M are identical to those shown in 3N, as these quantifications were performed in parallel. (N) Relative abundance of mass counts in the 300 and 301-316 m/z mass spectrum of FA C18:0 stearate FAMEs from WT or *daf-2(e1370)* animals treated with 10 mM labeled sodium acetate ^13^C_2_ (99%) (n = 3 biological replicates). (O) LC-MS analysis of WT *C. elegans* treated with *fasn-1* RNAi or 10 mM L-Malic Acid with measurement of adenylate energy charge. Note that the vehicle measurements in S6O are identical to those shown in 5A, as these quantifications were performed in parallel (n = 4 biological replicates). Lifespans are representative of 3 biological replicates (Table S3). Data represent the mean ± SEM. Imaging data are representative of at least 30 animals aggregated from 3 biologically independent experiments per condition, unless otherwise noted. Significance was assessed by (A-K) log rank analysis, and (L-O) two-way ANOVA with correction for multiple comparisons. ns p > 0.05, * p < 0.05, ** p < 0.01, *** p < 0.001, **** p < 0.0001.

**Figure S7.**
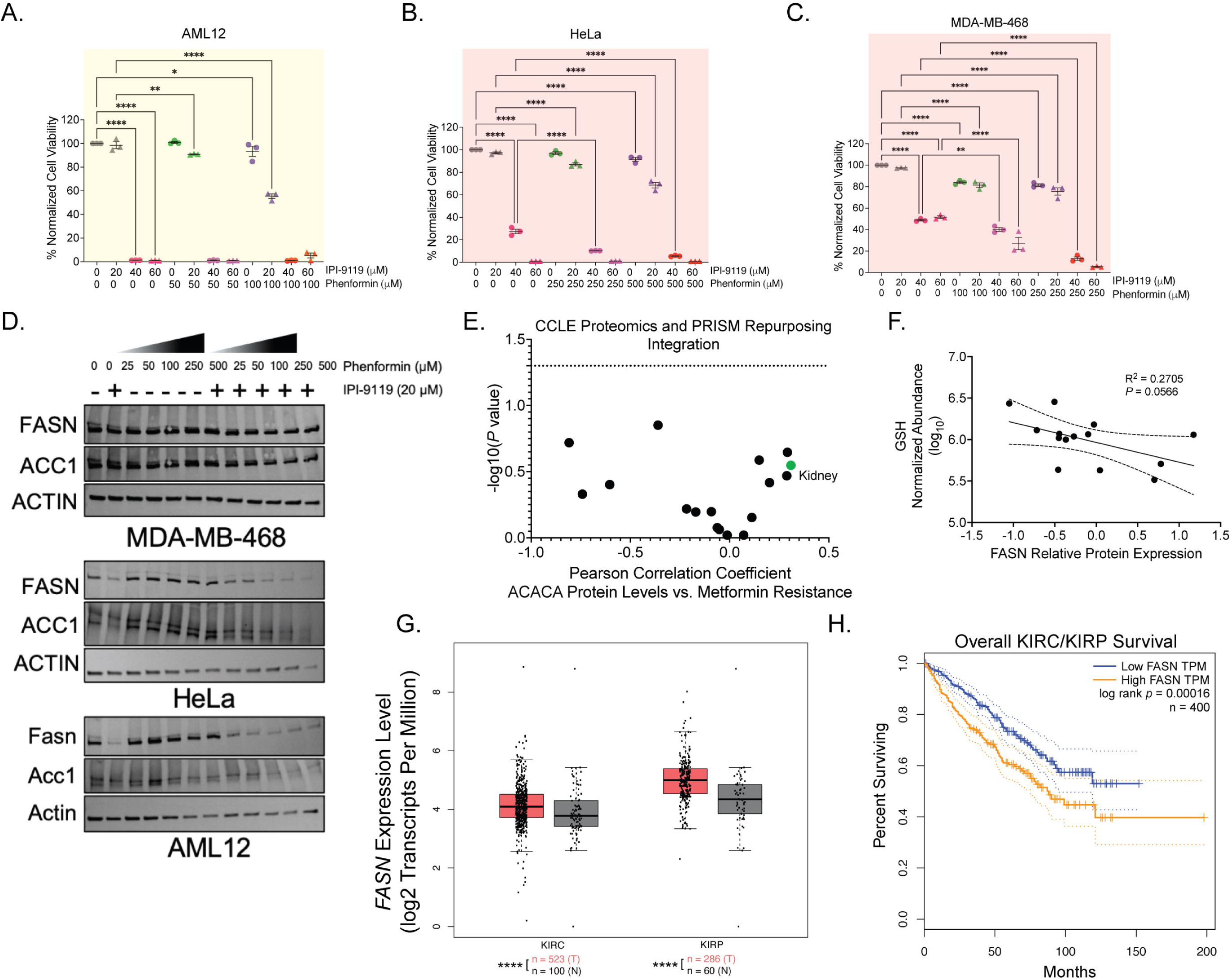
FASN activity buffers biguanide-induced energetic stress response in human and murine cancer cells. Related to Figure 7. (A-C) CVS intensity of AML12 (A), HeLa (B), and MDA-MB-468 (C) cells treated for 48 hours with the indicated doses of IPI-9119 and phenformin (n = 3 biological replicates). (D) Representative western blots for FASN/Fasn, ACC1/Acc1, and ACTIN/Actin in MDA-MB-468 (top), HeLa (middle), and AML12 (low) cells treated for 48 hours prior to analysis with the indicated doses of IPI-9119 and phenformin (representative of n = 3 biological replicates). (E) Pearson correlation coefficients of lines from the CCLE^110^, comparing ACC1 protein levels with their relative resistance to metformin-mediated growth inhibition and stratified by cellular tissue lineage. Data were extracted and analyzed using Data Explorer v2.0 on the DepMap^76^. Kidney lineage is highlighted in green. (F) Pearson correlation coefficients of kidney lineage cancer cell lines (CCLE)^74^, comparing their respective GSH metabolite levels with their FASN protein levels. (G) Relative *FASN* mRNA transcript expression in Kidney Renal Cell Carcinoma (KIRC) and Kidney Renal Papillary Cell Carcinoma (KIRP) tumors compared to tissue-matched normal controls extracted from the TCGA and GTEx databases. Data were analyzed using GEPIA^139^. (H) Overall disease survival analysis for individuals with KIRC/KIRP, stratified by high versus low *FASN* mRNA transcript expression. Data were extracted using GEPIA^139^. Cells were passaged/thawed and plated for 2 days prior to drug treatment for cells to reach at least 70% of confluency. Data represent the mean ± SEM. All non-IPI-9119 treated cells were treated with an equivalent percentage (v/v) of DMSO to control for off-target toxicity. Significance was assessed by (A-C) two-way ANOVA with correction for multiple comparisons, or (E-F) extracted from DepMap Data Explorer v2.0 and (G-H) GEPIA. ns p > 0.05, * p < 0.05, ** p < 0.01, *** p < 0.001, **** p < 0.0001.

**Data S1. Source values, statistical test result, and all uncropped western blots displayed in this manuscript. Related to Figures 1-7, Figures S1-S7, and STAR Methods.**

**Table S1. Results of genome-wide RNAi screen for effectors of 150 mM metformin-mediated growth inhibition. Related to Figures 1 and S1.**

**Table S2. Genes displaying differential translation efficiency (**ΔTE) in WT *C. elegans* treated with 4.5 mM phenformin. Related to Figures 1-2 and S1.

**Table S3. Tabular statistics for all *C. elegans* lifespan analyses and survival data included in this manuscript. Related to Figures 1-6, S1, and S3-S6.**

**Table S4. Genes displaying differential expression upon *fasn-1* RNAi in concert with 4.5 mM phenformin, 10 mM L-malic acid, or 2% (w/v) D-(+)-glucose treatment. Related to Figures 2, 4, 5, 6, and S4.**

**Table S5. Raw and processed GC-MS lipidomics and LC-MS metabolomics data presented in this study. Related to Figures 3, 4, 5, 6, S5 and S6.**

**Table S6. All oligonucleotides used during this study. Related to Figures 1-7, Figures S1-7, and STAR Methods.**

