## Supplementary material for "Reductive death is averted by an ancient metabolic switch": Data S2 - Unprocessed Immunoblot Images

Data S2, Ahsan et al.

Figure 2L

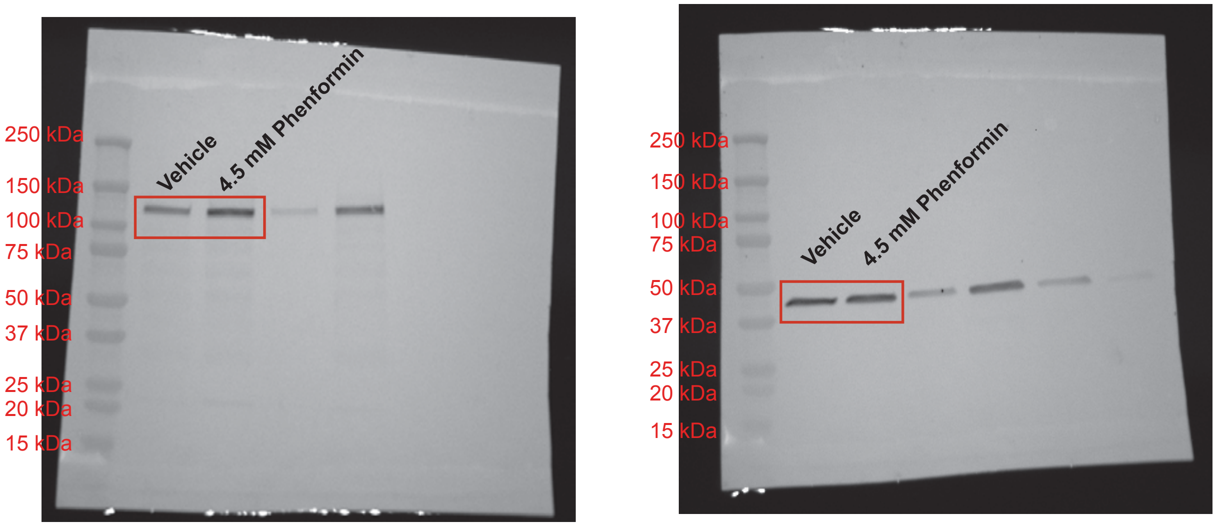


anti-HA (1:1000)

anti-Actin (1:2500)

Figure S2D

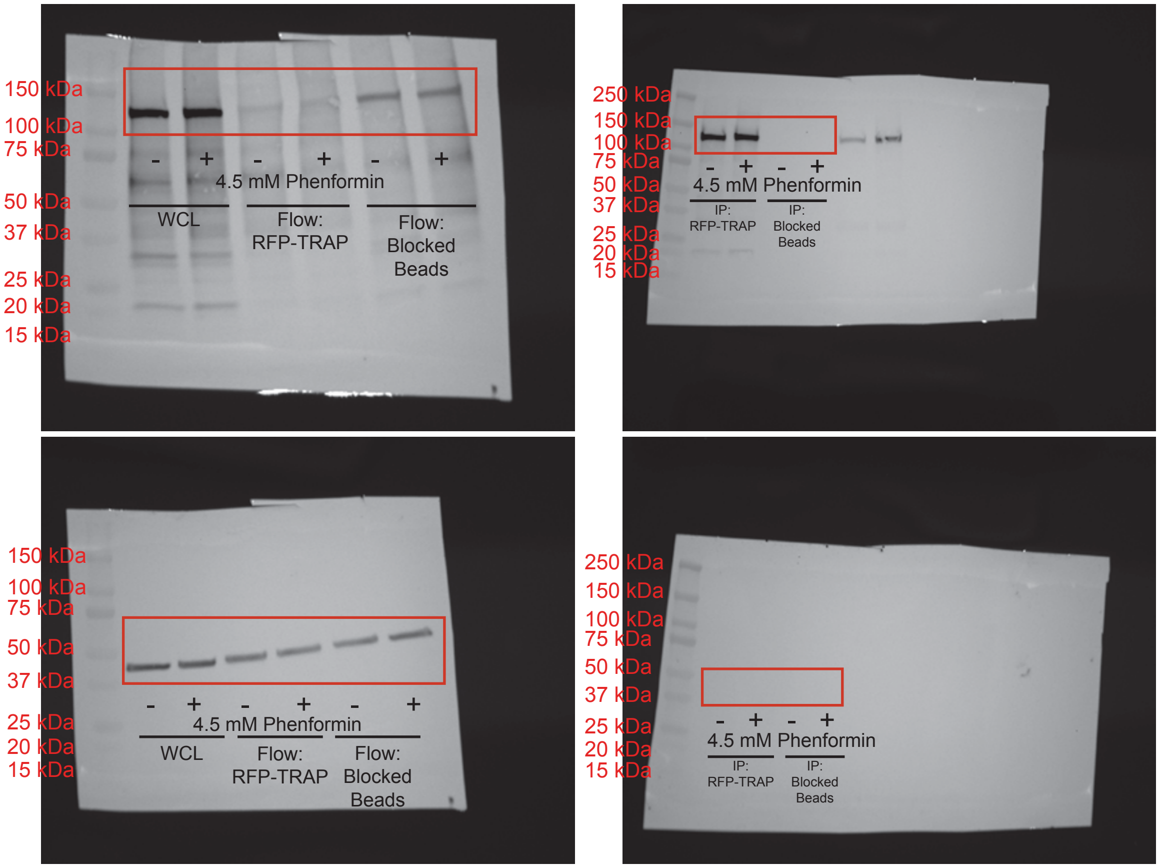


anti-HA (1:1000)

anti-Actin (1:2500)

Figure 4L

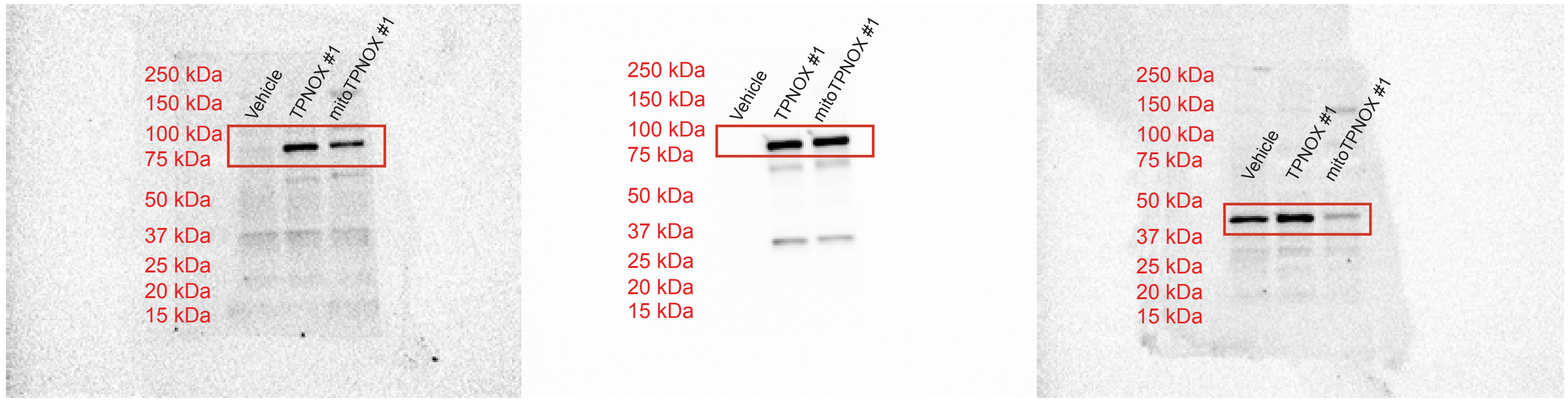


anti-FLAG (m2) (1:1000)

anti-GFP (D5.1) (1:1000)

anti-Actin (1:2500)

Data S2, Ahsan et al.

### Figure 5C

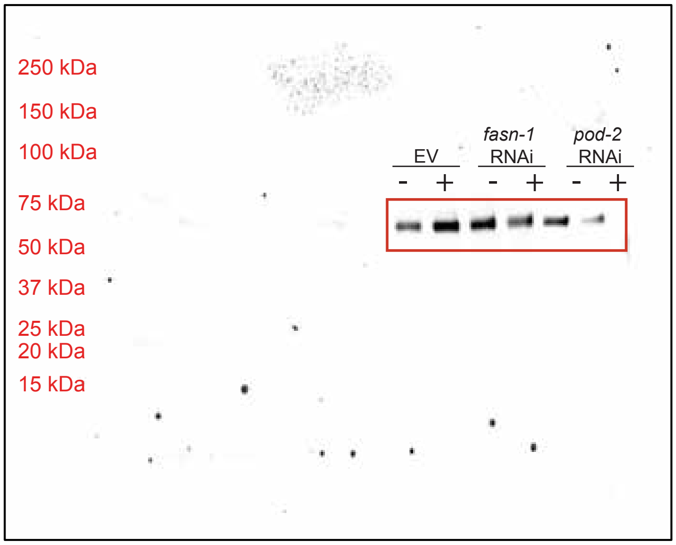


anti-phospho-AAK-2 (Thr172) (1:1000)

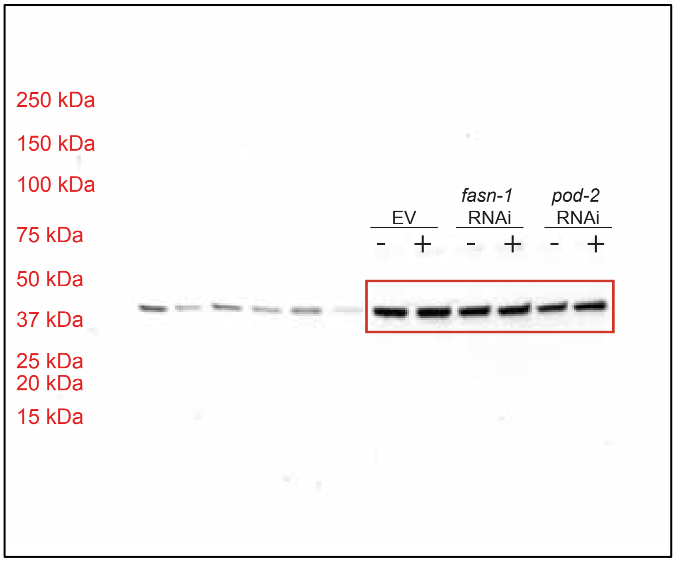


anti-ACT-1 (1:2500)

### Figure 7M

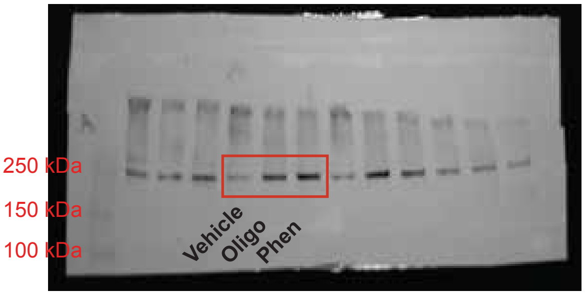


anti-Fasn (1:1000)

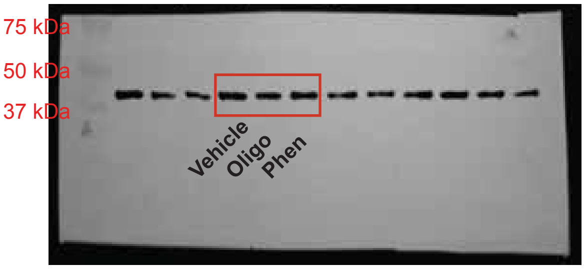


anti-Actin (1:2500)

### Figure S7D

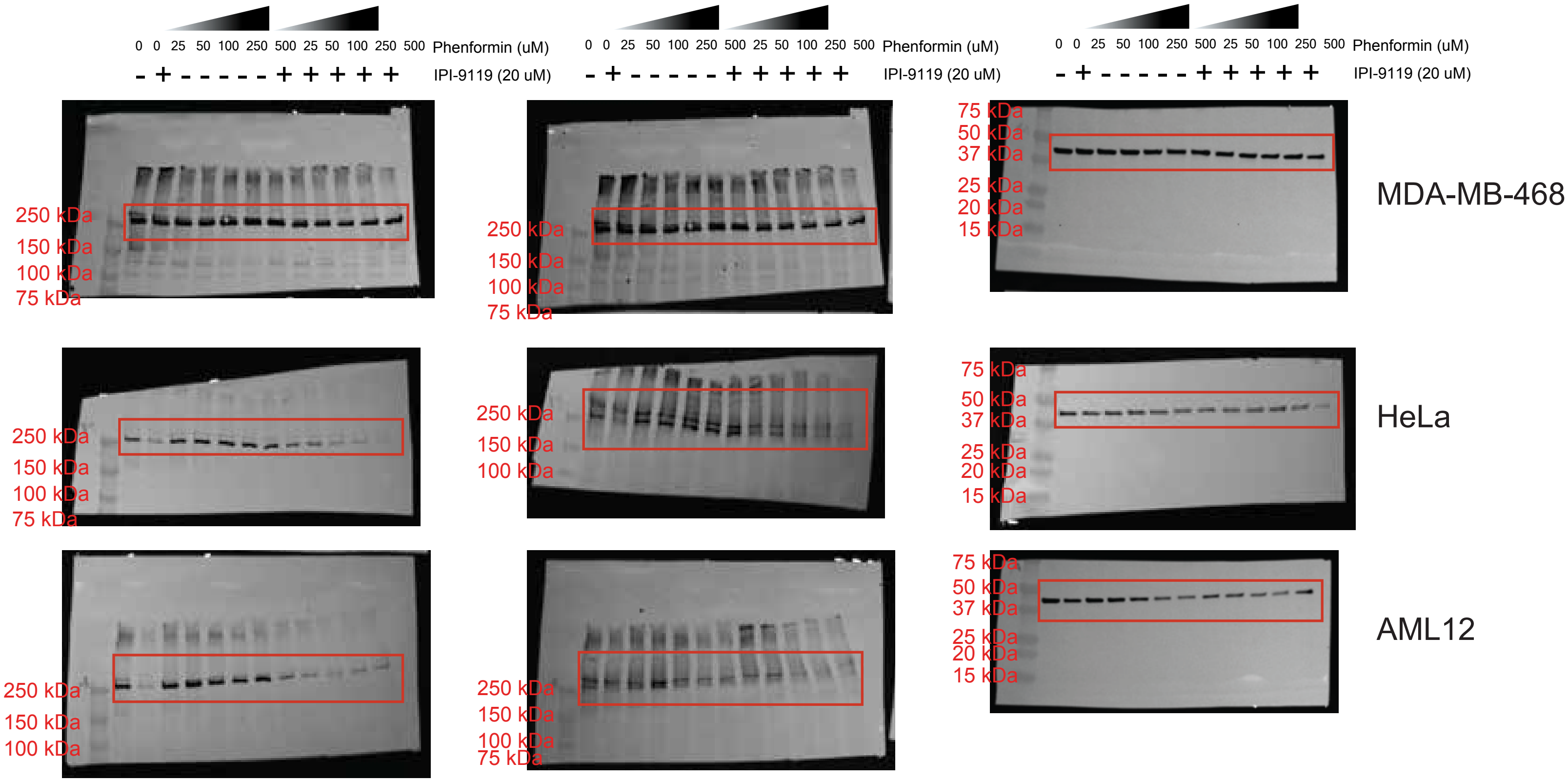


anti-FASN (1:1000) anti-ACC1 (1:1000) anti-ACTIN (1:2500)
